# A “Deep Dive” into the SARS-Cov-2 Polymerase Assembly: Identifying Novel Allosteric Sites and Analyzing the Hydrogen Bond Networks and Correlated Dynamics

**DOI:** 10.1101/2020.06.02.130849

**Authors:** Khaled Barakat, Marawan Ahmed, Yasser Tabana, Minwoo Ha

## Abstract

Replication of the SARS-CoV-2 genome is a fundamental step in the virus life cycle and inhibiting the SARS-CoV2 replicase machinery has been proven recently as a promising approach in combating the virus. Despite this recent success, there are still several aspects related to the structure, function and dynamics of the CoV-2 polymerase that still need to be addressed. This includes understanding the dynamicity of the various polymerase subdomains, analyzing the hydrogen bond networks at the active site and at the template entry in the presence of water, studying the binding modes of the nucleotides at the active site, highlighting positions for acceptable nucleotides’ substitutions that can be tolerated at different positions within the nascent RNA strand, identifying possible allosteric sites within the polymerase structure and studying their correlated dynamics relative to the catalytic site. Here, we combined various cutting-edge modelling tools with the recently resolved SARS-CoV-2 cryo-EM polymerase structures to fill this gap in knowledge. Our findings provide a detailed analysis of the hydrogen bond networks at various parts of the polymerase structure and suggest possible nucleotides’ substitutions that can be tolerated by the polymerase complex. We also report here three “druggable” allosteric sites within the nsp12 RdRp that can be targeted by small molecule inhibitors. Our correlated motion analysis shows that the dynamics within one of the newly identified sites are linked to the active site, indicating that targeting this site can significantly impact the catalytic activity of the SARS-CoV-2 polymerase.

## Introduction

SARS-CoV-2 is an enveloped, positive-sense RNA virus belonging to the Nidovirales family and has been identified as the cause of COVID-19 disease that began in Wuhan, China^1^. The virus is closely related to the severe acute respiratory syndrome coronavirus (SARS-CoV) and to several other bat coronaviruses^2^. The genome of SARS-CoV-2 is composed of 26-32 kilobases (kb), making it among the largest RNA genomes known to date. Replication of this genome is a fundamental step in the virus life cycle and inhibiting this mechanism has been proven recently as a promising approach in combating and controlling SARS-CoV-2 infection. A successful proof of concept for this approach is remdesivir, a nucleoside analog (NA) drug that has been repurposed recently against COVID-19^*3*^.

The SARS-CoV-2 replicase machinery involves several non-structural proteins (nsp)s encoded by the viral genome and assembled around the RNA-dependent RNA polymerase (RdRp), nsp12^4^. This includes the nsp7 and nsp8 RdRp cofactors; the dimer forming RNA binding protein, nsp9; the cofactor in viral replication, nsp10; and the nsp14 exoribonuclease (ExoN)^4-7^. The presence of the latter proofreading ExoN adds an additional challenge in blocking the replication of SARS-CoV-2. This ExoN is a CoV-specific intrinsic mechanism, which does not exist in other RNA viruses, and can simply remove the incorporated nucleoside analog (NA) and restore the viral polymerase function^8-9^. Although the interaction between all these proteins is important for the proper replication of the viral genome, several studies identified the nsp7-nsp8-nsp12 complex as the minimal functional assembly required to study this process. Additional studies showed also that the nsp12 on its own, although still functional, has low processivity, compared to the nsp7-nsp8-nsp12 complex^3^. Other studies argued that the nsp8 protein is capable of de novo initiating the replication process and has been proposed to function as a primase^10^.

The recent SARS-CoV-2 polymerase cryo-EM structures provide a detailed picture for the organization of the polymerase complex^5, 11^. A functional nsp7-nsp8-nsp12 complex involves a single nsp7 and two nsp8 monomers attached to the nsp12 RdRp (see Figure 1). In this structure, nsp12 adopts the conserved “right hand” shaped fold associated with all RNA viruses. This fold includes three distinct subdomains that can be clearly identified as fingers (S397 to A581 and K621 to G679), palm (T582 to P620 and T680 to Q815) and thumb (H816 to E920). In addition, the CoV-2 nsp12 includes an n-terminal extension domain (D60 to R249), which adopts a nidovirus RdRp-associated nucleo-tidyltransferase (NiRAN) domain^12^. This NiRAN subdomain is connected to the rest of the nsp12 structure through an interface formed by residues A250 to R365. Furthermore, residues D29-K50 form an additional n-terminal β-hairpin loop that is bound into a groove close to the NiRAN domain. The polymerase active site (i site) is located within the palm subdomain and is made up from the conserved RdRp A-G motives. Motif A comprises residues T611 to M626, which include the classical aspartate residue, D618. This residue helps in coordinating two catalytic divalent magnesium ions at the active site along with the two other aspartate residues, D760 and D761, located in motif C. Motif B includes residues T680 to T710, motif C includes residues F753 to N767, motif D contains residues L775 to E796, motif E includes residues H810 to V820, motif F includes residues K912 to E921, and finally, motif G includes residues K500 to S518^5^.

**Figure 1.**
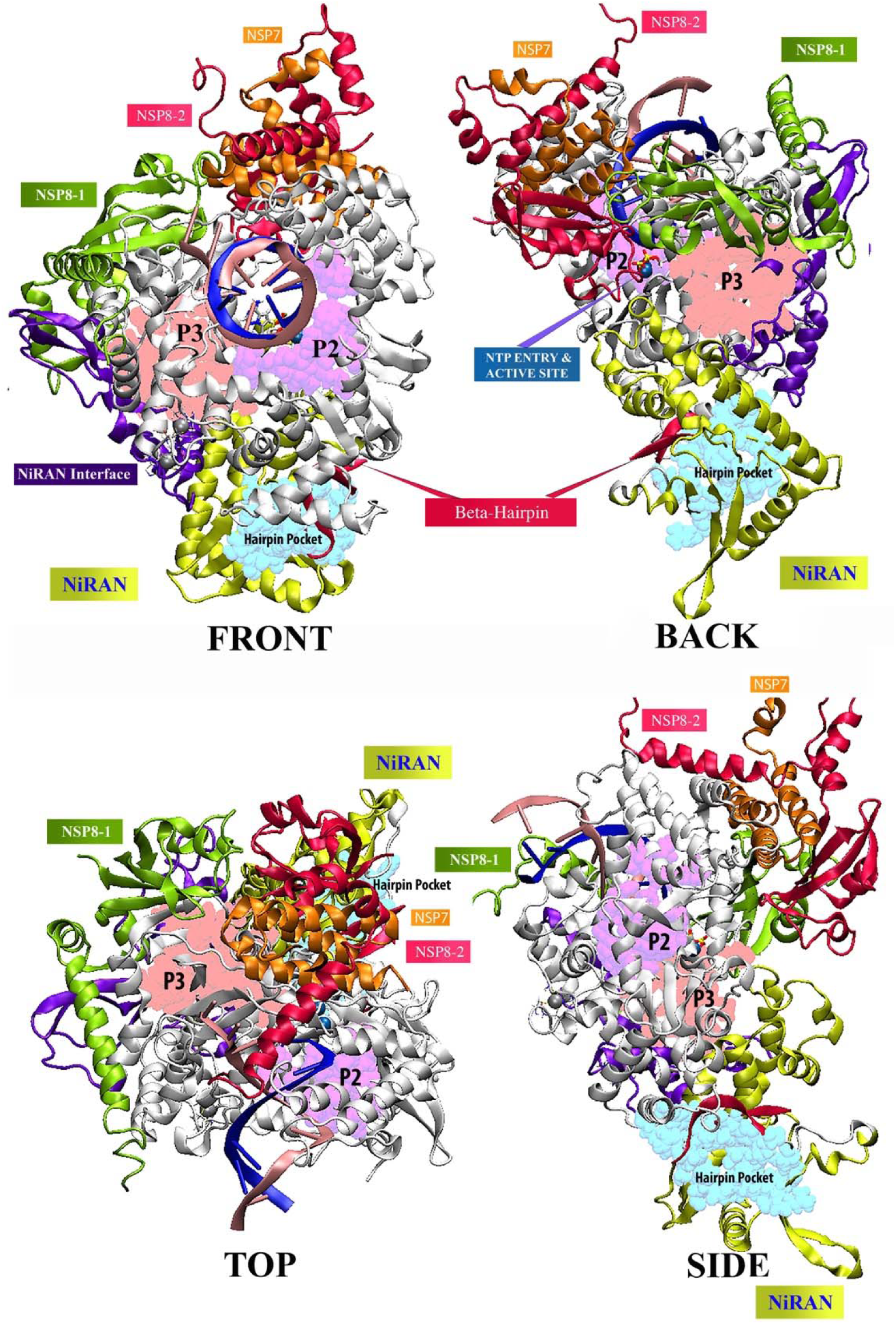
The overall structure of the nsp-7-nsp8-nsp12 polymerase as constructed in this work and the location of the three identified allosteric sites. The nsp12 RdRp domain is bound to nascent and template RNA strands. The β-hairpin pocket, colored in cyan is located in the proximity of the n-terminal β-hairpin loop (D29-K50). Pocket 2, colored in pink is located within the NiRAN interface subdomain and is close to the nsp8 monomer. Pocket 3, colored in mauve, is located close to the catalytic site.

To our knowledge, the present work is the first study to use the recently resolved SARS-CoV-2 cryo-EM structures to explore the CoV-2 polymerase complex. With the absence of a crystal structure for CoV-2, earlier studies used the CoV-1 polymerase as a template to construct homology models for the SARS-CoV-2 polymerase^8-9, 13-14^. Despite its similarity with CoV-2, the available Cov-1 cryo-EM structure lacked the first 116 residues of the nsp12 protein^5^. The importance of these residues will be discussed further below. Furthermore, these earlier studies did not go beyond the active site of the polymerase and most of them did not consider the dynamicity of the polymerase complex and did not study the hydrogen bond networks to the depth provided in the current work^8-9, 13-14^. Additionally, none of these studies investigated the polymerase complex for possible allosteric sites or studied the correlated dynamics within the polymerase structure. Therefore, the objective of the current study is to use the recent SARS-CoV-2 cryo-EM polymerase structures as a tool to fill this gap in knowledge. In this context, the current work is focused on understanding the dynamicity of the various polymerase subdomains, analyzing the hydrogen bond networks at the active site and at the template entry in the presence of water, studying the binding modes of various substrates at the active site, highlighting positions for acceptable nucleotides’ substitutions that can be tolerated at different positions within the nascent RNA strand, identifying possible allosteric sites within the polymerase structure and studying their correlated motion with the catalytic site.

To achieve these goals, we combined classical and accelerated molecular dynamics simulations with principal component analyses, conformational clustering, free energy calculations, binding site identification tools and correlated dynamics. The present work provides an overall view of the nsp7-nsp8-nsp12 polymerase complex, with the resolution of a single atom and with the ability to observe small and large conformational changes in the context of a bound RNA. Our findings analyze the hydrogen bond networks at various parts of the polymerase structure and suggest possible nucleotides’ substitutions that can be tolerated by the polymerase complex. We also report here three “druggable” allosteric sites within the nsp12 RdRp that can be targeted by small molecule inhibitors. Our correlated motion analysis reveals a connection between the dynamics within one of the new identified sites and the polymerase active site, indicating that targeting this site is expected to significantly impact the catalytic activity of the SARS-CoV-2 polymerase.

## Results

### Initial assessment of the polymerase complex

Figure 1 shows the complete structure of SARS-CoV-2 nsp12-nsp8-nsp7 polymerase complex, as constructed in this work, based on the recently resolved Cryo-EM structures^5, 11^ (see methods). In this model the “right hand” shaped RdRp domain of nsp12 is bound to nascent and template RNA strands and clearly reveals the NiRAN subdomain and its interface to the RdRp domain. Furthermore, it highlights the location of the n-terminal β-hairpin loop (D29-K50), which inserts into a groove close to the NiRAN domain. The catalytic active site (*i* site) within the polymerase domain contains two magnesium ions that are well-coordinated by three aspartate residues (D618, D670 and D671) (see below). It also includes two zinc ions that are chelated with a histidine and three cystine residues, each. Similar to the Cryo-EM structure (PDB ID: 6M71)^5^, our model also shows the correct proportions of nsp7, nsp8 and nsp12 as well as their conserved orientations and interfaces within the complex. Our initial structural assessment shows that the missing residues in the original Cryo-EM structures were properly added to the three protein structures. This is evident by the local quality estimates and Ramachandran plots, which shows > 95% of the nsp12 protein structure in the favoured regions for the dihedral angles with only 0.43% Ramachandran outliers. Similar scores were obtained for both the nsp8 and nsp7 proteins (see Figures S1 to S3, supplementary data).

### Stability and dynamics of the polymerase assembly

With all components of the SARS-CoV-2 polymerase complex are properly repaired and assembled, our next step was to study the dynamicity of the whole complex in a functional and physiological state. To do that, we first constructed nine polymerase complexes. This included five complexes in which a nucleotide triphosphate (NTP) was incorporated within the active site. These five NTPs comprised the four normal RNA nucleotides (*i.e.* ATP, CTP, GTP and UTP) in addition to remdesivir triphosphate. Hereafter, these five systems are called the BOUND systems. In addition to the five BOUND systems, we constructed four complexes in which the nucleotides are incorporated in the (i+1) position in the nascent RNA. As the polymerase catalytic site is free from substrates in these complexes, we call these systems the FREE systems. To evaluate the dynamicity of all nine systems, both FREE and BOUND structures were subjected to 100 ns MD simulations. Figure 2 shows the root mean square deviation (RMSD) and atomic fluctuations (beta factors) of the backbone atoms of the nsp12 protein in the BOUND systems (Figure 2A, B) and FREE systems (Figure 2C, D). The RMSD data for both BOUND and FREE systems show that nsp12 behaved relatively similar and was stable in all nine simulations. All nsp12 equilibrated conformations fluctuated from 1.9 Å to 2.5 Å from their initial reference structures, with the UTP system showing the most deviation in the BOUND complexes and remdesivir is showing the least. In the FREE systems the (i+1) incorporated ATP and GTP systems are showing a final similar RMSD behaviour and occupying the upper bound of the RMSD spectrum, while the (i+1) incorporated CTP and UTP systems are showing a lower deviation from the initial structure. The atomic fluctuations for both BOUND and FREE systems (Figure 2C, D) shows several regions with very little flexibility, such as the bound NTPs, the two chelated zinc ions and the two magnesium ions within the catalytic site (see Figure S4). However, this data also shows portions of nsp12 that exhibit large flexibilities (Figure 2C, D), with the most flexible regions comprising the n-terminal β-hairpin (D29-K50), the NiRAN subdomain (D60-R249) and the NiRAN interface subdomain (A250-R365).

**Figure 2.**
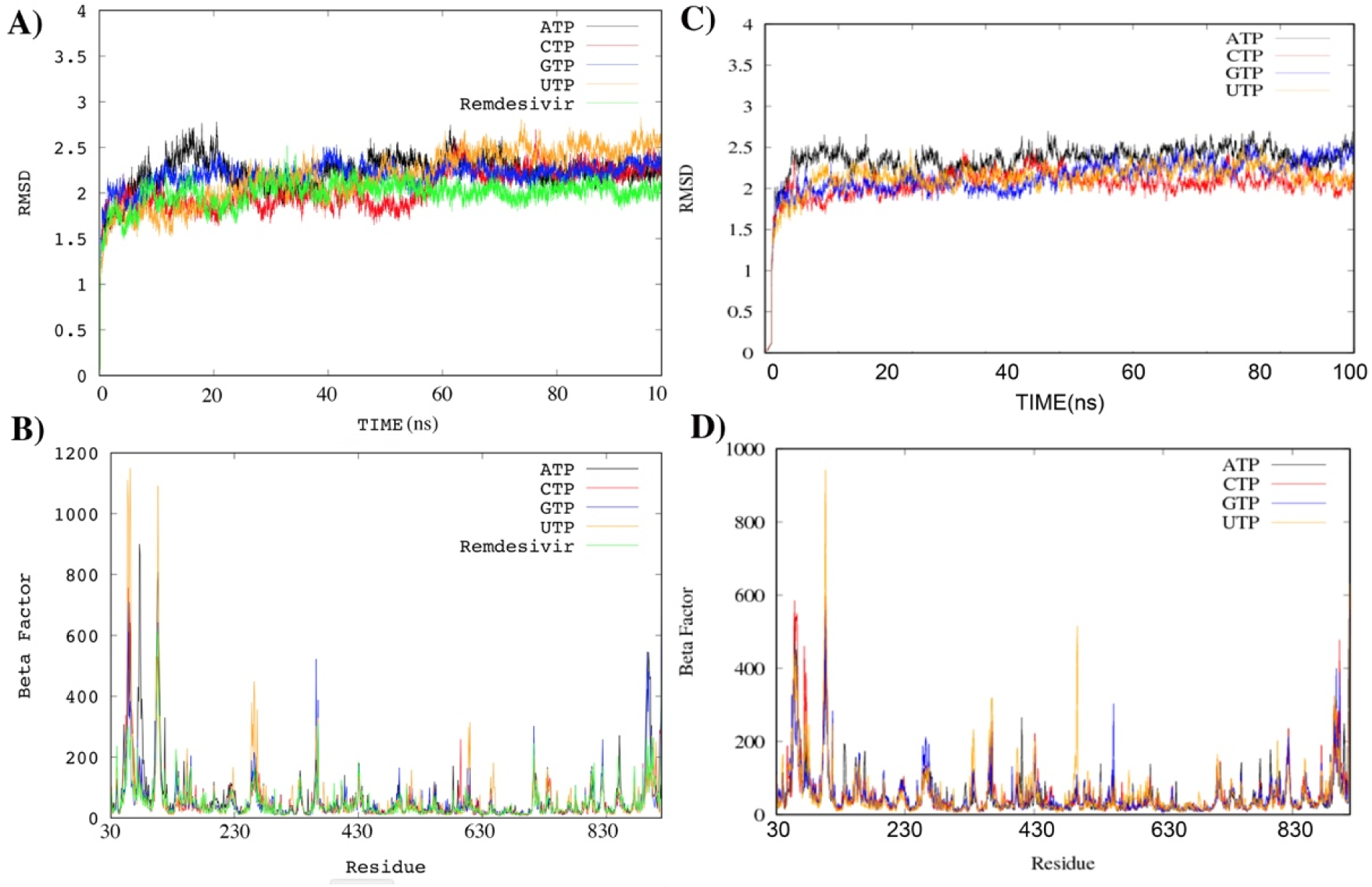
The root mean square deviation (RMSD) and atomic fluctuations (beta factors) of the backbone atoms of the nsp12 protein in the BOUND systems (A, B) and FREE systems (2C, D). The RMSD data for both BOUND and FREE systems show that nsp12 behaved relatively similar. The atomic fluctuations for both BOUND and FREE systems show several regions with very little flexibility, such as the bound NTPs, the two chelated zinc ions. The most flexible regions in nsp12 include the n-terminal β-hairpin (D29-K50), the NiRAN subdomain (D60-R249) and the NiRAN interface subdomain (A250-R365).

The RMSD and atomic fluctuations for the nsp7 protein and the two nsp8 monomers are shown in Figures S5 to S7 (see supplementary data). With the exception of the FREE CTP system, all nsp7 structures did not deviate significantly from the initial structure. NSP7 in all bound systems deviated from 2 Å to 3 Å from its initial structures (Figure S5A). While the same trend was shown in three of the FREE systems, nsp7 in the fourth CTP free system, has deviated almost 5 Å from its initial structure (Figure S5C). This is also reflected in the atomic fluctuations for the nsp7 residues (Figure S5B, D). While for both systems the nsp7 residues S1 to L60 show a clear rigidity during the nine MD simulations, the last ten residues, representing the c-terminal of nsp7 show much larger flexibility. While the nsp7 rigid residues are sandwiched between nsp12 and nsp8, the flexible nsp7 c-terminal are more liberated and interact only with two alpha helices from the nsp7-bound nsp8 monomer (see Figure S8). The RMSD and atomic fluctuations for this nsp8 monomer are shown in Figure S4. Both NTP BOND and FREE systems deviated from 4 Å to 6 Å from their initial structures (see Figure S6A, C), with the FREE systems are showing the greatest fluctuations (see Figure S4C). The most flexible regions in this nsp7-bound nsp8 monomer are the n-terminal residues M67 to R80, where the CTP BOND and FREE systems are showing the largest fluctuations at this region. This part of nsp8 is free to move in our model and has been recently shown to interact with the product RNA (PDB ID: 6YYT, to be published by Hellin et al.). Additionally, residues T107 to I120 in this nsp8 structure form a short alpha helix that is connected to a turning loop and seem to be flexible in both BOUND and FREE systems (see Figure S8). Interestingly, a bound GTP at the catalytic site seems to induce significantly large atomic fluctuations in this region, compared to all other bound and incorporated NTPs.

The second nsp8 monomer did not have a large deviation from the initial structure, compared to the nsp7-bound nsp8 monomer (Figure S7). It is worth mentioning that this monomer was clearly characterized in both Cryo-EM structures that were used to construct this model, indicating its less flexibility compared to the other nsp8 monomer. However, similar to the nsp7-bound nsp8 monomer, the n-terminal of this nsp8 monomer shows the largest flexibility in this structure as this region is also free to move in our model and has been recently shown recently to interact with the product RNA (PDB ID: 6YYT, to be published by Hellin et al.). A few other regions of this protein are showing some flexibility, including residues T107 to I120, described above.

Figure S9 shows the RMSD and atomic fluctuations of the nascent RNA strand. All NTP BOUND systems had and average RMSD around 3 Å at their final equilibrated states (Figure S9A). On the other hand, the nascent strand RMSD data for the FREE systems (Figure S9C) showed an interesting trend, where one can see a clear separation in RMSD averages, ranging from 2 to 5 Å, with the ATP system showing the most deviation from the initial structure. It is worth emphasizing that these systems represent the elongated state of the polymerase, where the bound nucleotides have now been incorporated within the nascent RNA strand and translocated from the (i) site to the (i+1) position, and the (i) site is now empty, waiting for a new incoming nucleotide to be incorporated. The atomic fluctuations for all five BOUND systems exhibit a similar trend (Figure S9B), where the first five nucleotides (i+1) to (i+5) are very stable within the RNA channel. Only nucleotides beyond the (i+5) position are showing large flexibility. These nucleotides are no longer surrounded by the nsp12 residues as they are now extending past the product RNA hybrid exit channel, and are only attached through their base-pairing to the template nucleotides. Similarly, the five nucleotides at positions (i+1) to (i+5) are also stable in the FREE systems (Figure S9D) and nucleotides beyond the fifth position are still flexible; however, they are less flexible, compared to their corresponding nucleotides in the BOUND systems. This RMSD and atomic fluctuations data show a clear difference between the BOUND and FREE states for the stability and dynamicity of the nascent RNA strand.

Figure S10 shows the RMSD and atomic fluctuations of the template RNA strand. With the exception of the ATP BOUND case, the template strand deviated from 2 to 4 Å from their initial structures (Figure S10A). There was a significant fluctuation in the RMSD for the ATP case. These fluctuations seem to emerge from the flexibility of nucleotides beyond the (i+5) position, where similar to the nascent RNA strand, these nucleotides are no longer bound by the protein residues and are extending beyond the hybrid exit channel and are only bound through base-paring with their complementary nucleotides in the nascent RNA strand. The RMSDs for the template strand in the FREE systems exhibit a similar trend to those of the nascent RNA strand, where a clear separation in the RMSD averages can be observed, with the GTP system showing the most deviation from the initial structure (Figure S10C) and has the most fluctuating nucleotides (Figure S10D).

### Principal component analysis (PCA)

Figure 3 shows the magnitudes of the eigenvalues for the first 10 eigenvectors for the BOUND systems. It also shows the projections of their trajectories on the first and second eigenvectors for each system. It is important to emphasize that this PCA was focused on the catalytic site, represented by the bound NTP at the (i) site, the two magnesium ions as well as all the protein and RNA residues within 5 Å from the bound NTP. Our main objective was to study all possible modes of binding of the bound NTPs and to understand their interactions with the surrounding environment. The resulting eigenvectors constitute the essential vectors of the motion, where the larger an eigenvalue, the more important its corresponding eigenvector in the collective motion. As shown in Figure 3, for all BOUND systems the magnitudes of the eigenvalues for the eigenvectors are following an exponential decay, a feature that is associated with reaching an acceptable sampling of the trajectories. A detailed representation of the conformational dynamics can be obtained by projecting the trajectories onto the planes spanned by the most dominant eigenvectors, and these projections provide a visual tool to quantify the size of the conformational space sampled by these trajectories. The higher the occupancy of a conformational state in this projection, the lower the free energy of that state ^46,47^. Therefore, by observing the regions at which many conformations cluster, one can predict the minimal energy conformations visited by these MD trajectories and estimate the number of dominant modes of binding for each bound NTP. As shown in Figure 3, all bound NTPs visited more than one conformation, with the bound CTP seem to have a major mode of binding within the catalytic site. On the other hand, the bound UTP seem to have multiple binding modes. The projections for the bound ATP show that ATP is restricted between two major conformations with a possibility to adopt around 2 minor conformations. Figure 3 also shows that both GTP and remdesivir has visited two flexible dominant conformations as their projected trajectories highlight two main clusters spanning a wide area in their projections. For the FREE systems, all projections of the incorporated NTPs follow an almost identical trend (Figure S11), where all NTPs along with their surrounding environment seem to have visited two major conformations, well-separated in their projections. These two main conformations are accompanied with very minor conformations as shown in Figure S11.

**Figure 3.**
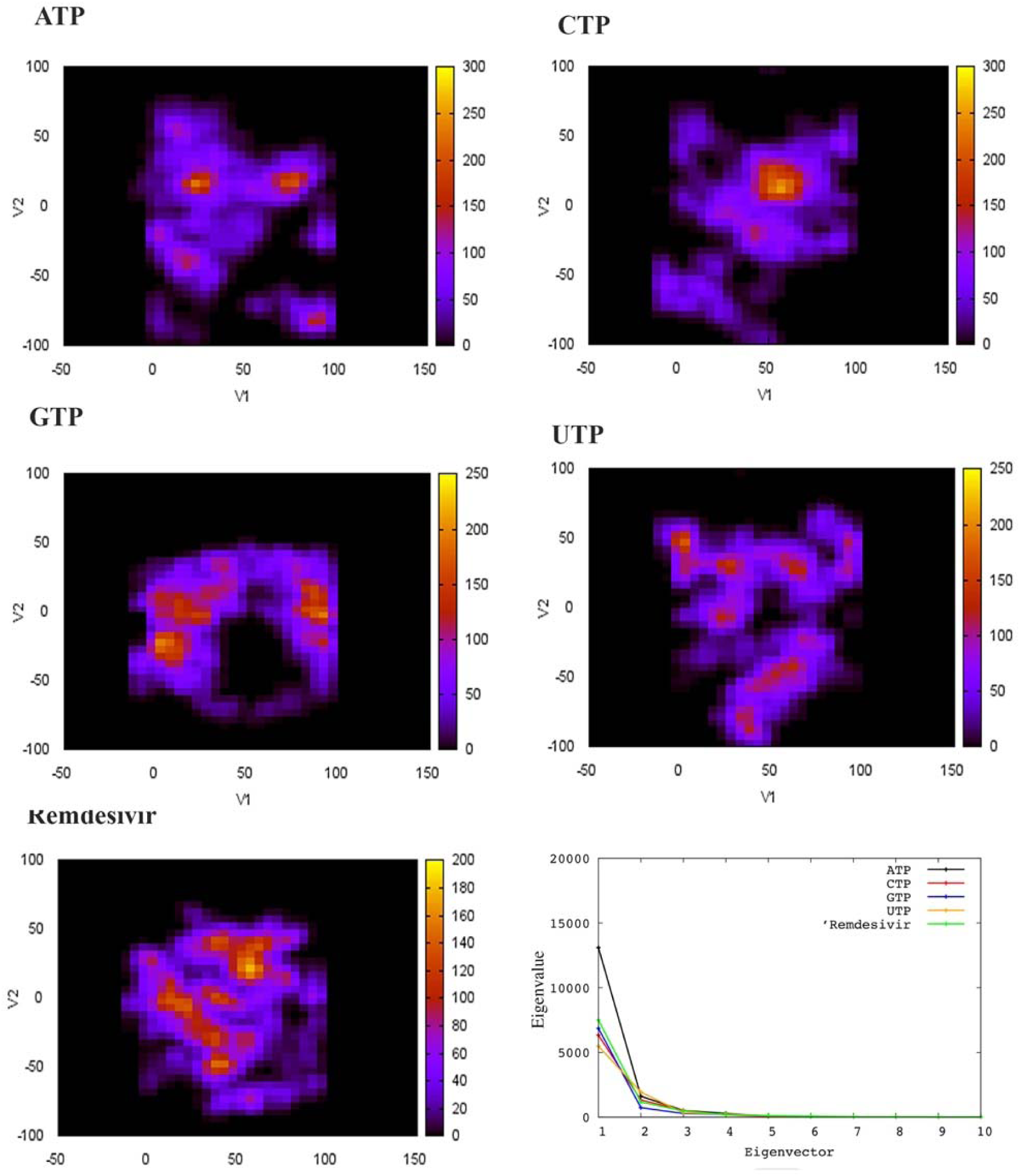
Principal component analysis (PCA) on the catalytic site. The magnitudes of the eigenvalues for the first 10 eigenvectors for the BOUND systems are following an exponential decay, a feature that is associated with reaching an acceptable sampling of the trajectories. Projections of the trajectories of the BOUND systems on the first and second eigenvectors are also shown.

### Clustering analysis

Our next step was to identify and to study the dominant conformations highlighted by the PCA projections described above. To do that we performed RMSD conformational clustering of all conformations visited by the bound NTPs. Figure 4 shows the clustering metrics observed for different cluster counts as well as the clusters’ sizes at the predicted optimal number of clusters for both remdesivir (Figure 4A, B) and ATP (Figure 4C, D). Similar analyses for the other three bound NTPs are shown in Figures S12 to S14. For remdesivir a local minimum in the DBI parameter is observed at a cluster count of 10, which coincides with a plateau in the percentage of variance explained by the data (SSR/SST), indicative of 10 clusters as the optimal number of clusters for remdesivir conformations. The largest cluster of those includes only around 30% of the whole trajectory, and the sizes of the second to the fifth clusters are comparable to the largest cluster, indicating the presence of more than a single dominant conformation for remdesivir. On the other hand, although 14 clusters are predicted to be the optimal number of clustering for the ATP trajectory, the dominant cluster in this case includes more than 70% of the whole ATP trajectory, indicating the existence of a main dominant conformation for ATP within the catalytic binding site. Figure 5 shows a superposition of the centroids of each cluster for ATP (Figure 5A) and remdesivir (Figure 5B). The superimposed centroids are coloured by the ranking of their corresponding clusters, where a red colour represents a higher ranking. For ATP, the representatives of the largest clusters are all grouped close to the dominant conformation. On the other hand, remdesivir seem to have at least two additional major conformations to its main mode of binding.

**Figure 4.**
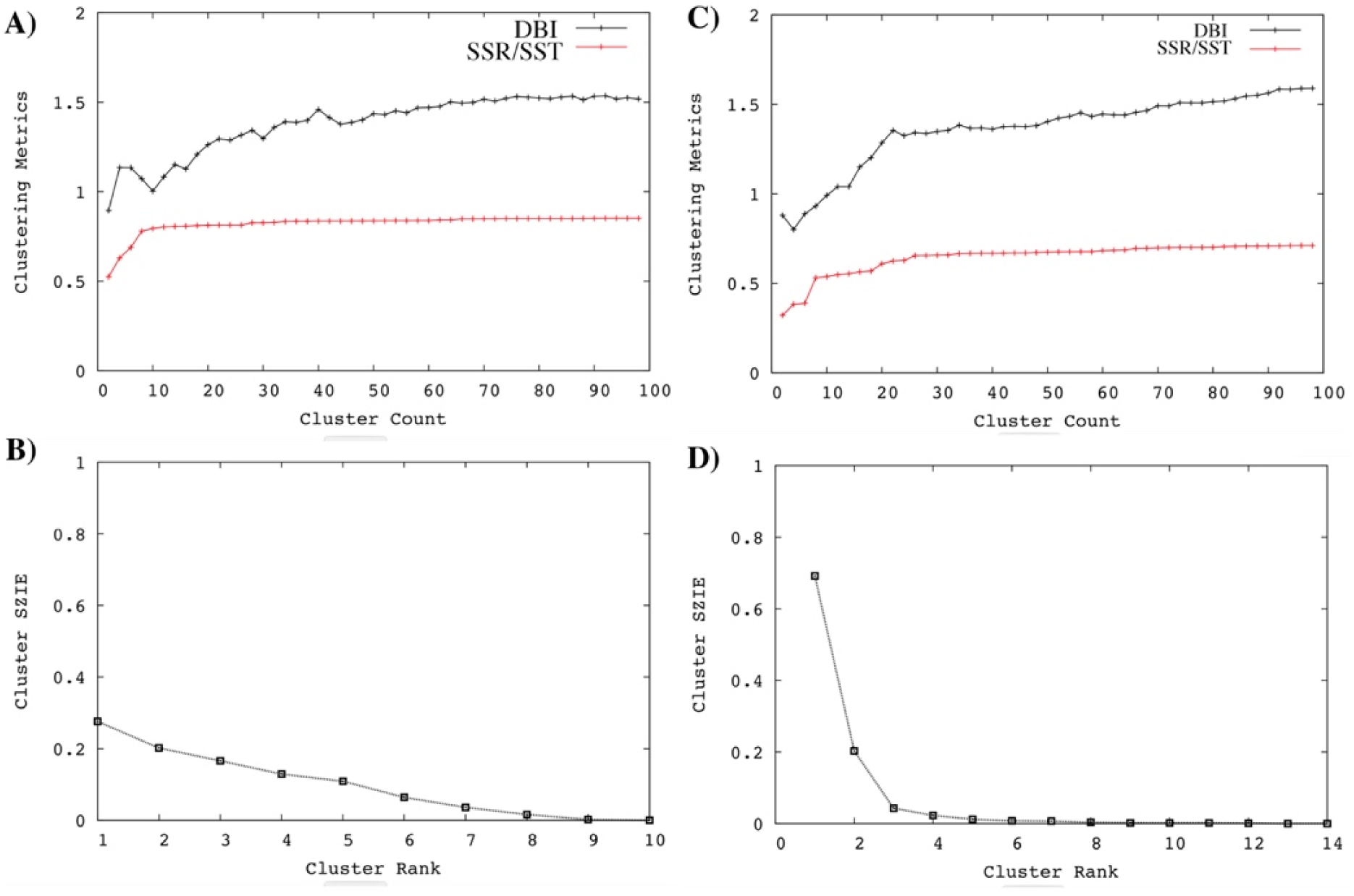
Conformational clustering of the bound nucleotides’ trajectories. Clustering metrics observed for different cluster counts are shown as well as the clusters’ sizes at the predicted optimal number of clusters for both remdesivir **(A, B)** and ATP **(C, D)**.

**Figure 5.**
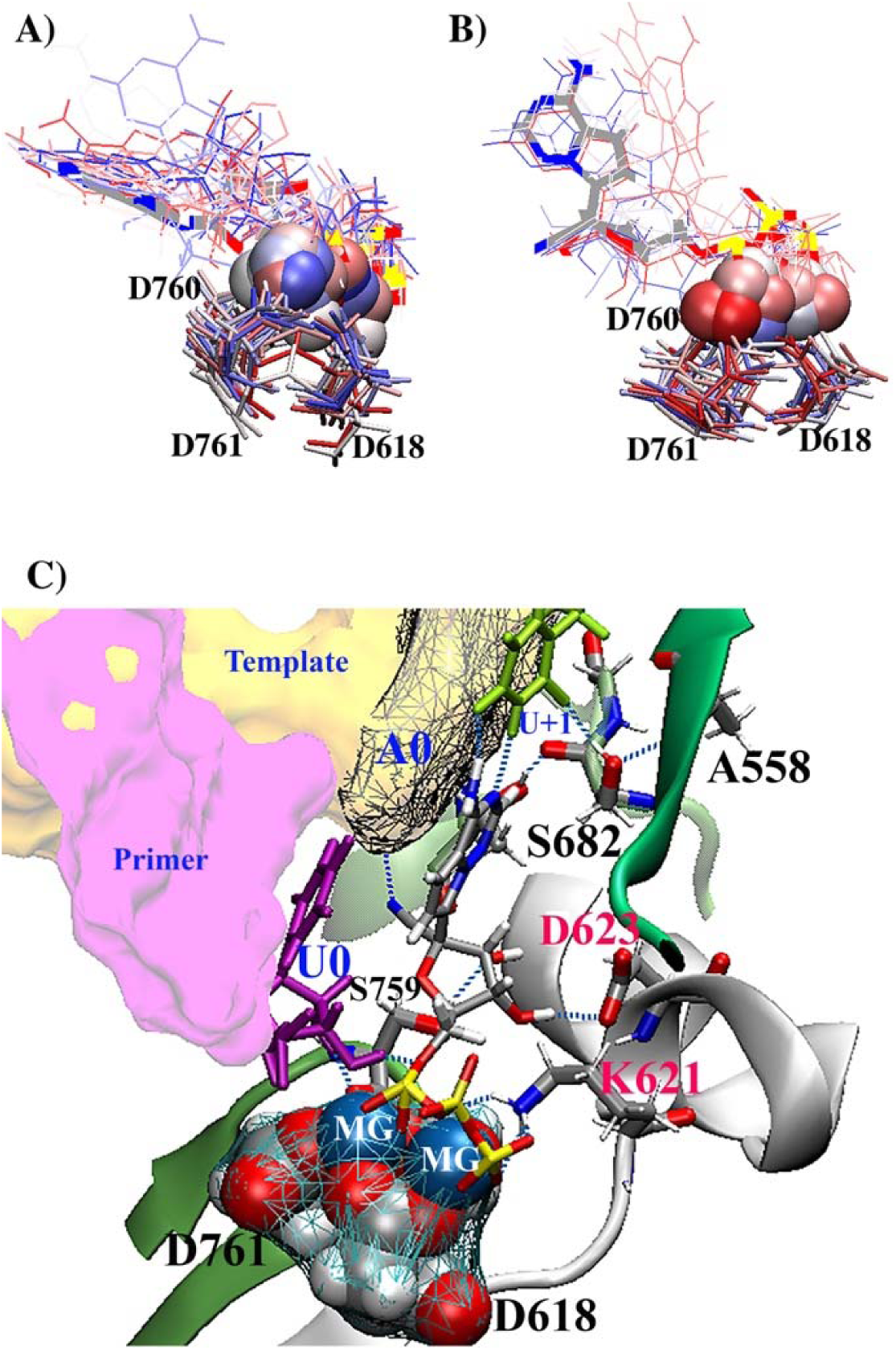
Binding modes of ATP and remdesivir. Superposition of the centroids of each cluster for ATP **(A)** and remdesivir **(B)**. The superimposed centroids are coloured by the ranking of their corresponding clusters, where a red colour represents a higher ranking. **C)** The main mode of binding from the largest cluster for remdesivir reveals the intricate hydrogen bonding network formed within the catalytic site. The triphosphate group from remdesivir are coordinating the two divalent magnesium ions, which are are also coordinated by the carbocyclic groups from the three deprotonated catalytic aspartates (D616, D670 and D671). The phosphate groups from remdesivir are further aligned in their position through at least two hydrogen bonds with the protonated K621. There is also an additional hydrogen bond between the remdesivir base and S682. This interaction is further strengthened through a hydrogen bond between the backbones of A558 and S682.

### NTPs dominant modes and hydrogen bonds analysis

The main mode of binding from the largest cluster for remdesivir is shown in Figure 5C. This binding mode reveals the intricate hydrogen bonding network formed within the catalytic site, which is needed to align the coming nucleotide in its correct confirmation to be incorporated within the nascent RNA. In this binding mode, the triphosphate group from remdesivir are coordinating the two divalent magnesium ions through strong ionic bonds. The magnesium ions are also coordinated by the carbocyclic groups from the three deprotonated catalytic aspartates (D616, D670 and D671). The phosphate groups from remdesivir are further aligned in their position through at least two hydrogen bonds with the protonated K621. The modified adenosine base from remdesivir is forming two hydrogen bonds with its complementary uracil base from the template. There is also an additional hydrogen bond between the remdesivir base and S682. This interaction is further strengthened through a hydrogen bond between the backbones of A558 and S682. The sugar moiety of remdesivir is properly oriented at the catalytic site through two hydrogen bonds. The first is between the sugar 3’ hydroxyl group and the deprotonated carbocyclic group of D623. The second hydrogen bond is between the oxygen atom of the sugar 2’ hydroxyl group and S759. Interestingly, the 1’-ribose cyano substitution in remdesivir forms an extra hydrogen bond with the template nucleotide at the (i+1) position. Finally, the 3’-hydroxyl group from the nascent RNA nucleotide at the (i+1) position is forming a strong hydrogen bond with the oxygen from the first phosphate group in remdesivir. This interaction is essential for nucleotide incorporation as a covalent bond is expected to be formed at this location.

The dominant modes of binding for the four types of RNA nucleotides are shown in Figure 6, where the positions for the 1’-ribose and 1’-ribose hydrogens are represented by vdW mauve spheres, to highlight the available space in these locations for possible substitutions. The ATP dominant mode of binding (Figure 6A) is very similar to that of remdesivir, where K621, D623 and S682 play the same roles described above. Despite this similarity there are still a few, but not significant, differences, such as the involvement of R625, which forms a hydrogen bond with the carbocyclic group of D623. This interaction helps D623 in aligning the ATP sugar moiety in its proper position. An additional difference is the loss of the hydrogen bond formed between the oxygen atom of the sugar 2’ hydroxyl group and S759.

**Figure 6.**
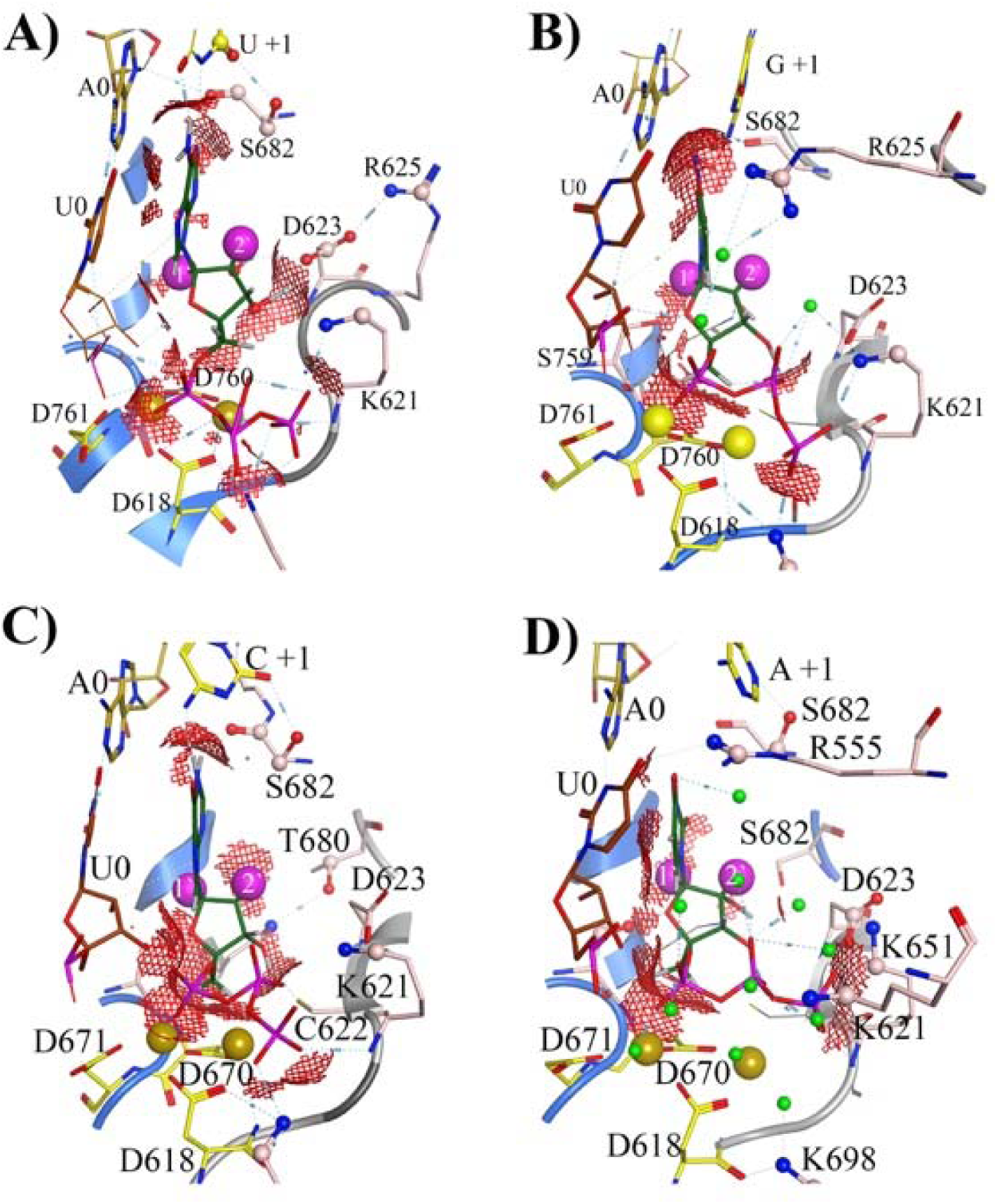
The dominant modes of binding for the four types of RNA nucleotides. The positions for the 1’-ribose and 1’-ribose hydrogens are represented by vdW mauve spheres, to highlight the available space in their locations for possible substitutions. **A)** The ATP dominant mode of binding is very similar to that of remdesivir. **B)** The dominant mode reveals an interesting role for water molecules in coordinating the hydrogen bond network. **C)** The GTP mode of binding retains the preserved coordination between the its phosphate groups and the two divalent magnesium ions. the lost hydrogen bond with D623 is compensated for by a hydrogen bond with T680. **D)** The UTP mode of binding shows a more intricate water network.

The mode of binding for CTP (Figure 6B) reveals an interesting role for water molecules in coordinating the hydrogen bond network for various interactions. It also emphasizes the importance of studying the polymerase complex in an explicitly solvated environment, otherwise these types of water mediated interactions would not be recognized. Similar to the case for ATP and remdesivir, K621, D623 and S682 continue to play the same roles described above in positioning the bound nucleotide for the chemistry to take place. The hydrogen bond between K621 and the phosphate groups are further strengthened via an extra water mediated hydrogen bond. Moreover, R625 is connected to the oxygen of the phosphate group in the nascent RNA nucleotide at position (i+1) through two water mediated hydrogen bonds. The mode of binding also shows the return of S759 to form a hydrogen bond with CTP sugar moiety.

The bound GTP (Figure 6C) seem to have a different set of interactions with the catalytic binding site residues. The GTP mode of binding retains the preserved coordination between the its phosphate groups and the two divalent magnesium ions as well as its strong hydrogen bond with the 3’-hydroxyl group from the nascent RNA nucleotide at the (i+1) position. However, GTP lost its hydrogen bond with D623, although this lost interaction is compensated for by a hydrogen bond with T680, which seems to play the same role of D623 for other nucleotides. In this mode of binding, D623 is forming a hydrogen bond with K621, which still interact with the phosphate groups, as described above.

The UTP mode of binding is shown in Figure 6D. Although in this binding mode UTP seems to lost its base-pairing hydrogen bonds with the template, the UTP bound nucleotide is still aligned properly through a more intricate water network, compared to the CTP case. For example, the UTP 3’ hydroxyl group is forming two water-mediated hydrogen bonds with the deprotonated carbocyclic group of D623, which is further strengthened by an additional hydrogen bond with K651. R555 is forming a hydrogen bond with the nascent RNA nucleotide base at the (i+1) and is keeping the base-pairing template nucleotide at position (i) in close proximity to the bound GTP through a hydrogen bond, which is supported by an additional hydrogen bond with S682.

The persistence of the important hydrogen bonds described above for both the bound NTP and template alignment are shown in Figures S15 and S16 for the BOUND and FREE systems, respectively. All bound NTPs, including remdesivir, show stable hydrogen bonding with the base-paring template nucleotide at the (i) position as well as with S759 and D623. Remdesivir has the most persistent and dominant hydrogen bond interactions with K621 and in the presence of remdesivir seem to strengthen the hydrogen bond interactions between A558 and S682. A bound UTP seem to have the dominant hydrogen bond interactions with D623, while a bound ATP sustains a strong interaction with S759. GTP forms the strongest interactions with the template nucleotide at the (i+1) position.

Aside from the interactions with the base-pairing nucleotide at the (i) position, most of the hydrogen bond interactions described above are weekend once the bound nucleotide is translocated from the (i) to the (i+1) position (see Figure S16). For example, all incorporated nucleotides lost their hydrogen bond interactions with the template nucleotide at position (i+1) and only UTP and ATP show very weak interactions with S759. On the other hand, D623 and S682 seem to still strongly interact with the incorporated nucleotide at position (i+1), possibly due to the lack of a bound nucleotide at the (i) site.

### The template at the entry

Figure S17 [now figure 7] shows the hydrogen bond network associated with the RNA template strand at the entrance to the catalytic site for the remdesivir case as an example. This network of hydrogen bonds reveals the complex interactions between the RNA template backbone and the nsp12 residues at the template entrance. In these interactions the protonated R569 seem to play a major role in orchestrating this network. It forms at least 3 strong hydrogen bonds with the backbones of the two template nucleotides at positions (i) and (i+1). It also forms a hydrogen bond with K500, which in turn forms a hydrogen bond with the backbone of the complementary nucleotide to remdesivir. The backbone of S501 forms two hydrogen bonds with the backbone of the two nucleotides preceding the (i) site. An additional hydrogen bond is formed between N507 with the last nucleotide in the template strand.

**Figure 7.**
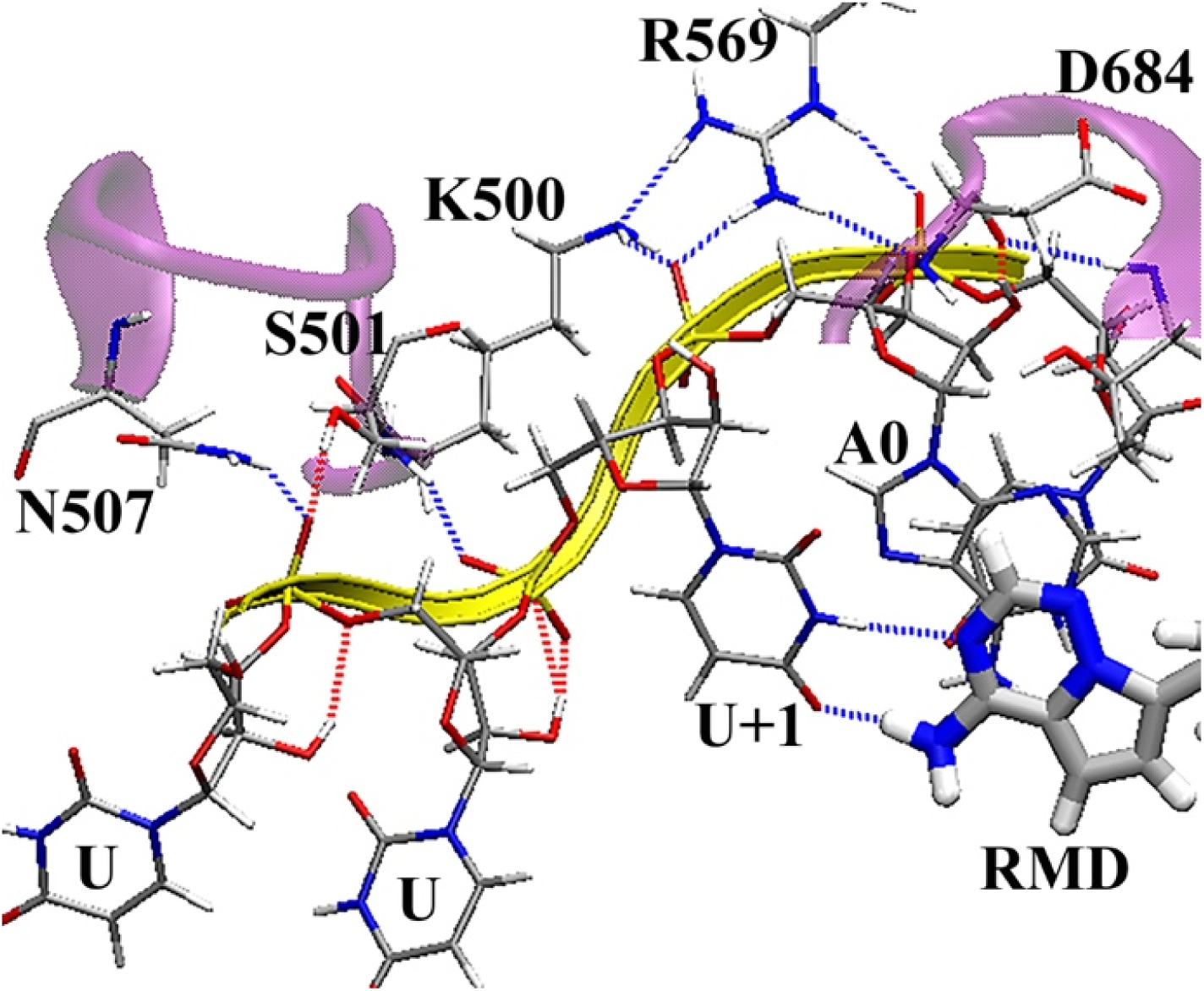
The hydrogen bond network associated with the RNA template strand at the entrance to the catalytic site for the remdesivir. The protonated R569 plays a major role in orchestrating the hydrogen bond network network. The backbone of S501 forms two hydrogen bonds with the backbone of the two nucleotides preceding the (i) site. An additional hydrogen bond is formed between N507 with the last nucleotide in the template strand.

Figures S17 and S18 show the persistence of hydrogen bonds between nsp12 protein and the RNA template at the template entrance gate, for the BOUND and FREE systems, respectively. R569 is clearly playing a key role in aligning the template at this position. Its hydrogen bond interaction with the variable bas-pairing nucleotide at position (i) (denoted as V(i) in Figures S17 and S19) is very strong, in both BOUND and FREE systems. R569 also forms a strong hydrogen bond with K500. This interaction seems not to be largely affected with the type of bound nucleotide at the active site. On the other hand, the hydrogen bond between S501 and the template nucleotide preceding the base-paring nucleotide (*i.e.* U(i-1)) has been predominantly lost in the FREE systems, except for the UTP system.

### Dihedral angles

Figures S19 and S20 shows the dihedral angles for the catalytic binding site amino acids, for the BOUND and FREE systems, respectively. The dihedral angles for the four conserved residues D618, S759, D760 and D761 are restricted throughout the simulations for all systems. This is mainly due to their key role in coordinating the two magnesium ions. While both dihedral angles for K621are constrained to ∼ - 40°to ∼80° when an NTP is present at the (i) site, both angles can move freely when the bound NTP is translocated to the (i+1) site. Similarly, the dihedral angels for D623 and S682 span a larger area in the FREE systems compared to their positions when a nucleotide is present at the catalytic site.

### Electrostatic Calculations

Figure 8 shows the electrostatic potential mapped over the whole polymerase complex. The negative electrostatic potential of the RNA strands is surrounded by a predominantly positive potential from the surrounding basic proteins’ residues. Figure 8B shows a detailed electrostatic potential view of the catalytic site with the bound remdesivir. The presence of the two divalent magnesium ions as well as the basic lysine residue, K621, provided a favourable positive electrostatic environment for the negatively charged phosphate groups of remdesivir. The side chains for K545, V557, K621, D623, S682, T687, A688 and N691, provide a positive potential cage for the bound remdesivir nucleotide and its complementary template base, UMP, where the negative electrostatic potential of the 1’-ribose cyano substitution is completely buried between S682, T687, A688, N691from one side, and K545, V557 from the other side.

**Figure 8.**
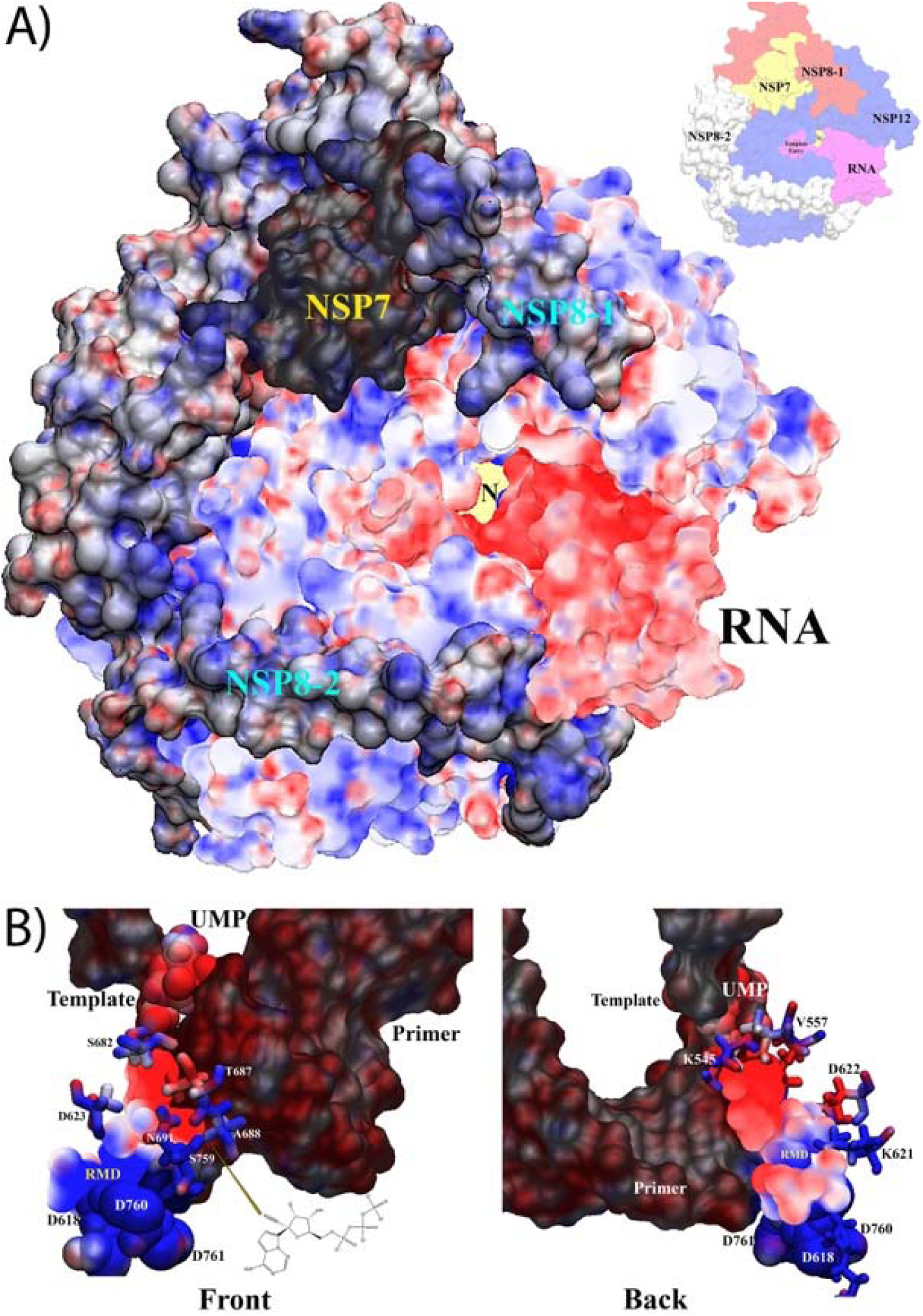
Electrostatic potential map. **A)** The electrostatic potential mapped over the whole polymerase complex. The negative electrostatic potential of the RNA strands is surrounded by a predominantly positive potential from the surrounding basic proteins’ residues. **B)** The electrostatic potential view of the catalytic site with the bound remdesivir. The side chains for K545, V557, K621, D623, S682, T687, A688 and N691, form a positive potential cage for the bound remdesivir nucleotide and its complementary template base, UMP, where the negative electrostatic potential of the 1’-ribose cyano substitution is completely buried between S682, T687, A688, N691from one side, and K545, V557 from the other side.

Figure S21 shows the electrostatic potential for the environment surrounding the two chelated ZINC ions. Each of the two positively charged ZINC ions are chelated with three deprotonated cystine residues and a histidine residue, protonated at their epsilon positions. The four residues on each site provide the necessary electrostatic environment to coordinate and confine the two ZINC ions. The nearby positively charged nsp12 amino acids also provide a favourable environment for the bound RNA in the background.

### Interaction energies at the active site

Table S1 shows the total vdW, electrostatic energy and total non-bonded interactions for remdesivir and the three nucleotides, UTP, CTP and GTP, relative to the ATP energies. Although remdesivir show very similar vdW energy compared to ATP, its electrostatic energy is significantly larger. This is mainly due to the 1’-cyano group substitution, which is buried in the electrostatic cage described above. While UTP seem to have the largest electrostatic interactions within the active site (∼-135 kcal/mole), it also shows the most unfavorable vdW interactions among all nucleotides (∼+35kcal/mole). GTP shows the same total energy as ATP. Its slightly unfavorable vdW interactions (∼+6 kcal/mole) is compensated for by (−6kcal/mole) from its electrostatic interactions. A bound CTP shows the same trend as for remdesivir. Although all bound NTPs as well as remdesivir show the same energetic trend in their interactions with the two magnesium ions and three aspartate residues (see Figure S22), the unfavourable UTP vdW energies seem to stem from its interaction with site. These unfavourable interactions are compensated for by electrostatics interactions with the magnesium ions and their attached aspartate residues (D618, D760 and D761) (see Figure S22) as well as favourable interactions with D623 (see Figure S23). Furthermore, all bound NTPS tend to interact similarly with S795 and all show better interactions with this residue compared to ATP, with an average total relative energy of ∼-3 kcal/mole for all NTPs (see Figure S24).

A bound remdesivir shows the most favourable energetics for the overall bound RNA, with around 90 kcal/mole lower energy compared to a bound ATP (see Figure S25 and Table S2). The second favoured NTP for the bound RNA is CTP, which chows around 50 kcal/mole overall lower energy compared to the bound ATP case. On the other hand, a bound ATP seems to be more favourable to the RNA compared to bound UTP and GTP, which lead to around ∼+13 kcal/mole and ∼+86 kcal/mole higher energy for the RNA, compared to ATP.

### Water accessible surface area for NTPs

Figure S26 shows the water accessible surface area (SASA) around the different NTPs in the BOUND (Figure S26A) and FREE (Figure S26B) systems. As expected, remdesivir shows the largest SASA, due mainly to the extended 1’-ribose cyano substitution. The difference in SASA between remdesivir and all other NTPs is clear since the beginning of the simulations. The second largest SASA is associated with the bound GTP. The SASA for all NTPs while they are binding at the catalytic (i) site is bigger than their SASA once they are incorporated within the nascent RNA. The smallest available SASA is for UTP in its (i+1) position, which is significantly smaller than other nucleotides. The SASA for all NTPs ranges from 250 Å^2^ for the uracil nucleotide at position (i+1) to 360 Å^2^ for remdesivir at position (i).

### Translocation of bound NTPs and implications for possible substitutions

Figure 9 shows the interactions of the RNA nascent nucleotides at position i+1 (Figure 9A) and i+2 (Figure 9B). It also shows possible substitutions positions (highlighted in green) that would be tolerable by the surrounding environment at these locations. Substitutions at these locations will allow the bound nucleotides at the active site (site i) to be translocated to sites i+1 and i+2. This can protect these incorporated nucleotides from the nsp14 exonuclease (ExoN). Among all possible substitutions, it seems that only modifications at the 1’-ribose and 3’-ribose as well as possible substations in the base would be acceptable at position i+1. Similarly, modifications at the 1’-ribose and 3’-ribose are still acceptable at the i+2 site. This is mainly because these two locations are not interacting with the surrounding environment (see Figure 9) and small modifications at these locations will not affect either the base-paring of the RNA nascent nucleotides with the template strand.

**Figure 9.**
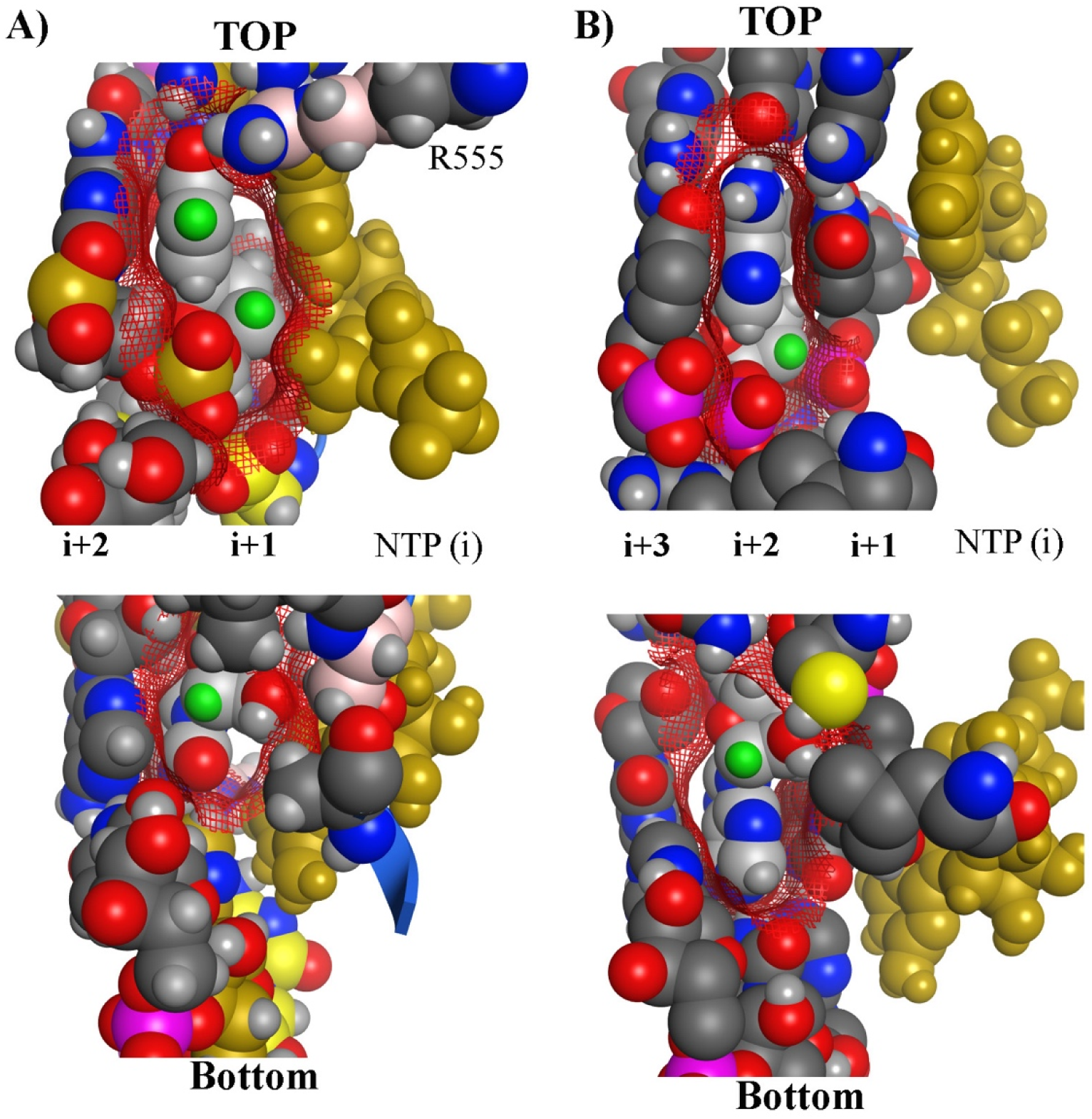
Possible nucleotides’ substitutions positions at the i+1 and i+2 sites. The interactions of the RNA nascent nucleotides at position i+1 (**A**) and i+2 (B). Possible nucleotide substitutions positions are highlighted in green. Substitutions at these positions are predicted to be tolerable by the surrounding environment.

### Allosteric Binding sites

Our binding site search tools (see methods) identified three potential allosteric pockets within the nsp12 polymerase. The locations for the three pockets are shown in Figure 1 and a close up on these pockets is shown in Figure S27. The first pocket (denoted here as the β-hairpin pocket) is located in the proximity of the n-terminal β-hairpin loop (D29-K50). This pocket is formed by the nsp12 residues F35 to N39, K50, N52, V71, K73, R74, R116, G203, V204, T206, D208, N209, Y217 to F222 and S236. The second binding site, denoted here as pocket 2, is located within the NiRAN interface subdomain and is close to the nsp8 monomer (see Figure 1). This pocket is formed by the nsp12 residues V315 to P323, F326 to P328, Y346 to E350, T394 to F396 to C622 to L630, S664, M666, V675 to P678. The third pocket (pocket 3) is located close to the NRA product exit channel (see Figure 1) and is formed by the nsp12 residues V588 to F594, W598, M601, A688, Y689, L758, F812 to A815, R836, S861to D865. It is important to emphasize that our site prediction tools identified serval pockets within the polymerase complex, however, these three pockets ranked the highest in terms of size, druggability and accessibility to small molecule compounds. To evaluate the accessibility and size of these pockets throughout the MD simulations, we studied the water hydration of these pockets, by counting the number of water molecules within the first and second hydration shells (see methods). Figure S28 shows the number of water molecules within the first two shells for the three pockets in the BOUND state, compared to the catalytic binding site. While the active site allows only up to 30 water molecules to enter in the presence of a bound NTP, the identified pockets allow from 150 to 250 water molecules in the first hydration shell, and up to 400 water molecules in the second hydration shell.

### Accelerated MD simulations

Figure 10A (Figure S29) shows the protein heavy atoms root mean square deviations (RMSD) values during the last 80 ns of the accelerated and classical MD simulations. The average RMSD values increase from 2.6 Å In the classical simulation to reach a value of 3.3 Å for the accelerated simulation. Maximum RMSD values also increased from 3.3 Å during the classical MD simulation to reach a value of 4.2 Å in the accelerated MD simulation. Both systems converge approximately after 10 ns of simulations time. The small increase in the average and maximum RMSD values is attributed to the biasing potential, which only exists in the accelerated simulations.

**Figure 10.**
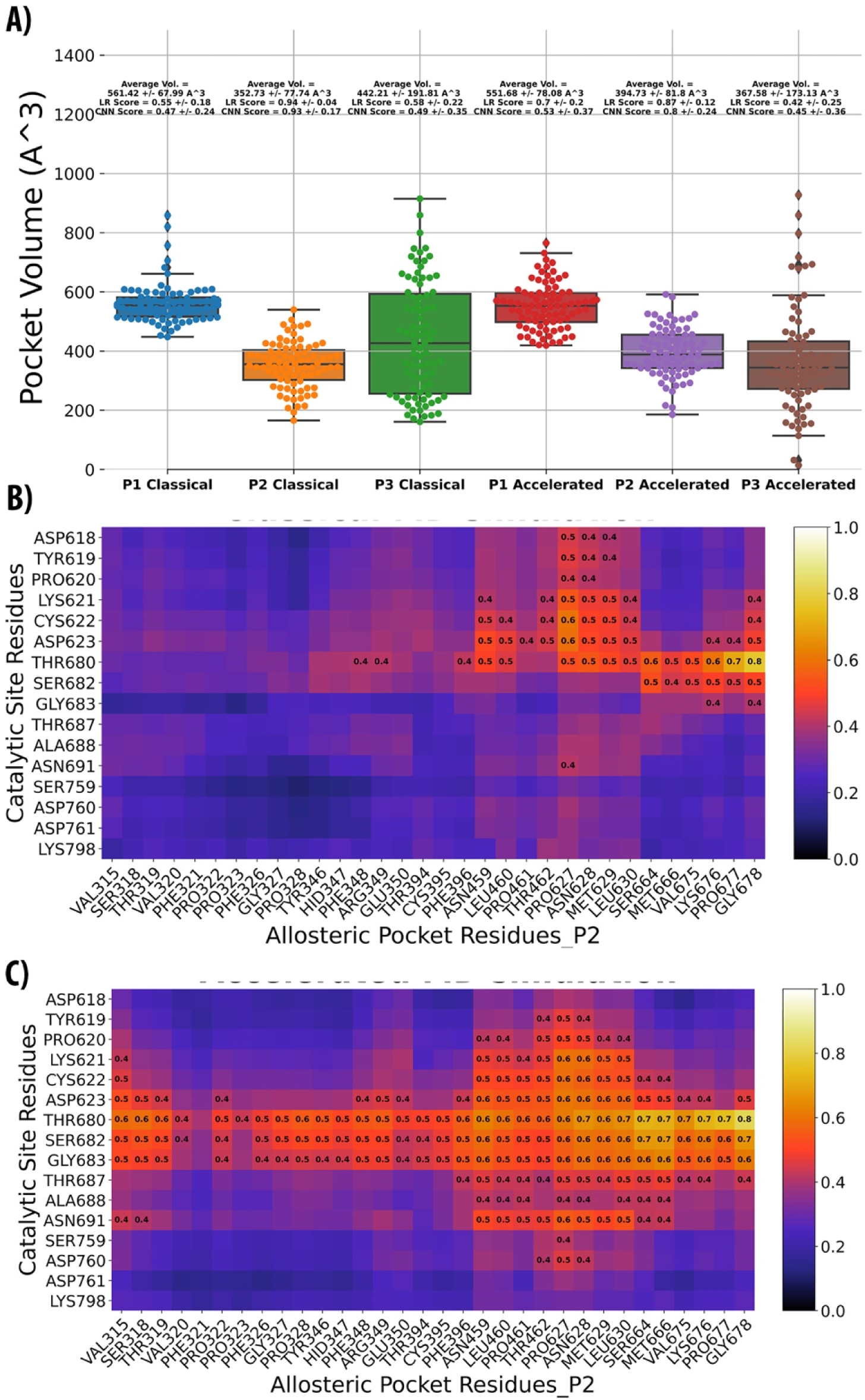
Druggability and correlated analyses. **A)** Boxplots showing the volume and druggability scores distribution for 500 snapshots extracted from the classical and the accelerated trajectories for the three pockets. **B & C)** residue correlation heatmaps focused on the catalytic residues relative to residues forming pocket 2.

### Pockets Average Volumes

The three identified pockets (see above) have been analyzed for their volume and druggability scores using TRAPP (see methods) throughout the classical and accelerated MDS. Figure 10 A shows the volumes and druggability scores data (*i.e.* logistic regression scores and CNN scores) depicted as a boxplot. The average pocket size for the three pockets ranged from 352.73 Å^3^ (for pocket 2) to 561.42 Å^3^ (for the β-hairpin pocket). Also, while the β-hairpin pocket and pocket 2 exhibit limited flexibilities, as evident by their reduced distributions of the standard deviation values of pocket sizes during the MD simulations (both classical and accelerated), pocket 3 showed a wider distribution of the standard deviation values (> 170.0 Å^3^). This indicates that this pocket has undergone a significant geometrical transformation during the simulations. This is not surprising given the close proximity of pocket 3 to the catalytic site. On the other hand, the maximum pocket size was observed for the β-hairpin pocket, with an average size of ∼552-562 Å^3^.

### Pockets’ Druggability and Residue Correlation Analysis

Figure 10A shows the TRAPP-LR and the TRAPP-CNN druggability scores calculated for the three pockets, employing the same MD frames used for volume calculations. Typical TRAPP-LR and TRAPP-CNN scores range between 0 (poor druggability score) to 1 (optimum druggability scores), with a consensus score of ∼ 0.9 from both models usually highlights acceptable druggable pockets. Figure 10A shows a dramatic difference in the predicted druggability scores of the three pockets, and these did not change between the classical and the accelerated MD simulations. Optimum druggability scores were observed for pocket 2, with an LR score ranging between 0.87±0.12 to 0.94±0.04 and CNN scores ranging between 0.93±0.17 to 0.8±0.24, for the accelerated versus classical simulations, respectively. On the other hand, the β-hairpin pocket and pocket 3 exhibited lower druggability scores, compared to pocket 2. The average LR druggability scores of the β-hairpin pocket is 0.55 in the classical simulation to 0.7 for the accelerated simulation (see Figure 10 A). A lower score (0.42 to 0.58) was observed for pocket 3. The same trend was observed for the CNN scores, ranging from 0.47 to 0.53 for the β-hairpin pocket and from 0.45 to 0.49 for pocket 3.

Given the good druggability scores for pocket 2, we decided to perform a more detailed correlation analysis on the amino acid residues forming this pocket to investigate their coupled motion relative to the catalytic site residues. This can provide more insights on the effect of targeting this pocket on the enzymatic activity of the catalytic site. In this context, the excellent memory utilization of the BIO3D package enabled us to use the full accelerated and classical MD trajectories (20,000 MD frames each) as inputs for the intended correlation analysis. The generated LMI correlation matrices were saved as comma-separated value files (CSV) and were processed with matplotlib to generate correlation heatmaps focused on the coordinated motions between residues forming pocket 2 and residues within the catalytic. The heatmaps (Figures 10B and 10C) are colour-coded by the correlation coefficient values. To facilitate the interpretation, we have also explicitly recorded the correlation values for cells that showed correlation values >= 0.4.

As shown in Figures 10B and 10 C, there is a relatively strong correlated motion between residues in pocket 2 and the catalytic site residues. As expected, this is more apparent in the AMDS trajectory, compared to the classical trajectory. For example, the correlation values of the catalytic site residue THR680 with several P2 residues such as P627, N628, M629, L630, S664, M666, V675 and K676) have been increased from (0.5, 0.5, 0.5, 0.5, 0.6, 0.5, 0.5, 0.6) in the classical MDS to (0.6, 0.7, 0.6, 0.7, 0.7, 0.7, 0.7, 0.7), in the accelerated simulation, respectively. Similar observations can be made for residues C622, K621, D623 and S682.

## Discussion

Replication of SARS-CoV-2 genome is a fundamental step in the virus life cycle and is mediated by a huge replicase protein complex, involving the nonstructural proteins (nsp)7 to nsp16^4^. While the RNA-dependent RNA polymerase (RdRp), nsp12, forms the core of this complex, several auxiliary proteins and cofactors are essential for the coordinated processes needed to produce the final RNA product^15^. Studying how these proteins associate together to form the viral replicase can have a significant impact on understating their structure, function and organization within this complicated machinery. It can also provide deep insights on the hydrogen bond networks formed within the polymerase structure and the important role played by water molecules in mediating these networks. Additionally, studying the stability, dynamicity and correlated motions within this complex can lead to the discovery of important regions and novel druggable hotspots to inhibit the replicase activity. These allosteric binding sites are usually hidden in the inactive/unbound forms of proteins and their discovery requires an overall view of the replicase complex, with the resolution of a single atom and with the ability to observe small and large conformational changes in the context of a bound RNA. The existence of such allosteric sites is supported by our findings (see results) as well as by earlier lessons from similar RNA viruses (*e.g.* the HCV polymerase has four druggable allosteric sites in addition to its active site^16-17^). The need to identify more druggable binding sites within the SARS-CoV-2 replicase machinery is highly desirable as targeting only the active site of the polymerase with nucleoside analogs (NA)s may not prove enough to fully inhibit viral replication. This is mainly due to the presence of an additional CoV-specific mechanism, which impairs NAs potency, represented by an exonuclease (ExoN), residing in the N-terminal of nsp14, with similarity to the active site in nsp12^9, 18-19^. This domain proofreads the generated viral genome and can simply remove the incorporated NA and restore the viral polymerase function^8-9^.

The recent SARS-CoV-2 polymerase cryo-EM structures^5, 11^) provide a unique opportunity to construct an accurate model for this complex. Therefore, the objective of the current work was to use these structures as a starting point and build several comprehensive dynamical models for the SARS-CoV-2 polymerase to study several aspects related to its function. This includes understanding the dynamicity of the various polymerase domains, analyzing the hydrogen bond networks at the active site and at the template entry in the presence of water, studying the binding modes of the nucleotides at the active site, highlighting positions for acceptable nucleotides’ substitutions that can be tolerated at different positions within the nascent RNA strand, identifying possible allosteric sites within the polymerase structure and studying their correlated motion with the catalytic site. Our results provide a detailed view of the SARS-CoV-2 polymerase complex in the context of these research questions. Furthermore, our findings suggest possible nucleotides’ substitutions that can be tolerated by the polymerase structure and reveal three possible allosteric sites within the nsp12 RdRp that can be targeted by small molecule inhibitors.

To our knowledge, this is the first study to use a SARS-CoV-2 structure to build the polymerase complex. A few recent homology modelling studies used the SARS-CoV polymerase as a template to construct a model for SARS-CoV-2^8-9, 13-14^. However, these studies did not go beyond the active site of the polymerase and most of them did not consider the dynamicity of the polymerase complex and did not study the hydrogen bond networks to the depth provided in the current work. Furthermore, none of these studies explored the polymerase complex for possible allosteric sites or studied the correlated dynamics within the polymerase structure. It is also important to emphasize that the CoV-1 nsp12 that was used in all these earlier studies as a template lacked the first 116 residues^5^. These residues include the n-terminal β-hairpin loop (D29-K50) and a major part of the NiRAN domain of nsp12. Aside from the importance of these residues in maintaining the stability and dynamicity of the whole nsp12 protein (see Figure 2), these residues also form one of the three binding sites that we identified in the current study (see Figure 1 and Figure S27).

Our findings revealed the detailed modes of binding at the catalytic site for all four types of RNA nucleotides as well as for remdesivir. Our results also show the importance of water molecules in mediating the interactions of the bound nucleotides – a factor that was never discussed or considered in earlier publications related to the SARS-CoV-2 polymerase. The bound CTP and UTP provide a clear example for the role of water in facilitating their interactions within the active site (See Figure 6). This is not surprising as our water shell analysis also show a presence of at least 20 water molecules within 4 Å from the bound nucleotide in the (i) site. This observation was noted for all bound nucleotides (see Figure 28S A), suggesting a major role for water inside the active site. It is important to also note that all the predicted binding modes have been obtained following a comprehensive PCA and clustering analyses. Although there is no guarantee that a complete equilibrium sampling of the nucleotides’ conformations has been reached, the achieved eigenvalues and the exponential decay of their magnitudes as well as the observed projections of the trajectories on the dominant eigenvectors (see Figure 3) suggest that an adequate conformational sampling for the catalytic site has been obtained for all simulations. RMSD conformational clustering analyses were then used to identify and to study the dominant conformations suggested by our PCA projections. Of the various clustering metrics described in the literature^20^ [refs], we used the elbow criterion and the DBI index as a measure for clustering convergence ^42^. These two metrics have been extensively tested and validated to study MD trajectories^20^ and the visualization of clustering convergence using these metrics is obvious. Here, we have applied this methodology by calculating the percentage of variance found within the data after each attempt to extract a new cluster from the simulated systems. As the number of clusters exceeds the optimal number, the percentage of variance plateaus, and the DBI exhibits a local minimum, indicating a complete extraction of the significant information included in the MD simulations^42^. This is illustrated in Figures 4 and Figures S12 to S14, where the elbow criterion suggested different cluster counts for the different bound nucleotides and the numbers of suggested clusters were in an excellent correlation with the projections obtained from PCA (see Figure 3).

The proposed binding modes not only explain how each nucleotide is interacting within the catalytic site, but can also suggest positions within these nucleotides to modify without affecting their hydrogen bond network. Since any drug designed to occupy the catalytic site has to be a derivative of a natural nucleotide, studying the hydrogen bond patterns and their persistence throughout the simulations for the individual nucleotides is tremendously important. If a loss of a particular hydrogen bond is not significant in altering the alignment of a given nucleotide within the catalytic site, then a modification at this position would be tolerable by the polymerase. Tolerable modifications at the i+1 and i+2 sites are also desirable and can also provide further protection for the incorporated drug from the nsp14 ExoN^9, 18-19^. For example, our modelling explains how the 1’-ribose cyano substitution in remdesivir is not only allowed within the active site, but also improves the electrostatic interactions relative to its parent scaffold, ATP. This cyano group forms an extra hydrogen bond with the template nucleotide at the (i+1) position (see Figure 5C) and its negative electrostatic potential is completely buried within a positive electrostatic potential cage formed by the nsp12 residues S682, T687, A688, N691, K545, V557 (see Figure 8). This provided remdesivir with a significantly favourable electrostatic energy (∼-100 kcal/mole) and with almost no difference in its vdW energy (∼zero kcal/mole), relative to ATP. This data is supported by the recent findings from Gordan et al.^13^, where their competitive inhibition RNA synthesis data suggested that remdesivir can compete with its natural counterpart, ATP and showed that the drug can be indeed incorporated within the polymerase more efficiently compared to ATP, with an IC_50_ of ∼32 nM ^13^. It is important to note that we used ATP as a reference in all our energy calculations as it was previously identified as the natural substrate of the SARS-CoV polymerase^21^.

Among all possible substitutions that can be integrated within the nucleotides, our modelling suggests that only modifications at the 1’-ribose and 3’-ribose as well as possible substations in the bases would be acceptable for a nucleotide to move from position i to positions i+1 and i+2 (see Figures 6 & 9). These possible substitution points in the bound nucleotides’ structures do not participate in any important interactions with the surrounding environment and can allow small modifications. The observed solvent accessible surface area (SASA) for the different nucleotides at positions (i) and (i+1) are ranging from 250 Å^2^ for the uracil nucleotide at position (i+1) to 360 Å^2^ for remdesivir at position (i) (see Figure S26). This SASA data can provide guidance on the allowable sizes for nucleotides’ modifications at the different sites within the polymerase. As discussed above, it is important for a modified nucleotide to be translocated from the (i) position to a further position within the nascent RNA strand to be protected from the nsp14 ExoN^9^. Our findings also show that the first 5 nucleotides (from position i+1 to i+5) in both RNA strands are very stable, indicating their tight binding and optimal interactions with the nsp12 polymerase residues forming the RNA channel (see Figures S9 and S10). A clear example of these strong interactions is the hydrogen bond network formed at the template entrance, where residues such as R569, K500 and N507 play a major role in orchestrating the hydrogen bond network stabilizing the RNA template nucleotides (see Figure S7). These hydrogen bonds are very stable and persistent throughout the MD simulations for both FREE and BOUND systems (see Figures S17 & S18).

A major finding from the current study is the identification of three “druggable” binding sites within the nsp12 RdRp (see Figure 1 and Figure S27). It is important to emphasize that recognizing these pockets directly in the initial Cryo-EM structures would not be possible. This is evident by our water shell analysis, which shows that the number of water molecules entering these pockets maximizes only after a significant time of our MD simulations and is almost ZERO during the first few nano seconds (see Figure S28). Furthermore, identifying the β-hairpin pocket (see results) would not be possible in the earlier SARS-CoV-2 polymerase studies, which used the nsp12 from CoV-1 as a template. As discussed above, the available CoV-1Cryo-EM structure lacks the first 116 residues^20^, which encompass the n-terminal β-hairpin loop (D29-K50) and a major part of the NiRAN domain of nsp12. The electrostatic map for this pocket is shown in Figure S30 and reveals a sizable rectangular pocket, extending along the β-hairpin loop, with a nicely distributed electrostatic potential, showing a negative electrostatic middle core in between two electrostatic positive regions at the north and south borders of the binding site. This β-hairpin pocket can accommodate at least ∼100 water molecules as indicated by our water shell analysis (see Figure S28). Our second pocket (pocket 2) is located within the NiRAN interface domain of nsp12 (see Figures 1 and S27). This is a smaller pocket, relative to the β-hairpin pocket and is formed by the nsp12 residues V315 to P323, F326 to P328, Y346 to E350, T394 to F396 to C622 to L630, S664, M666, V675 to P678. This pocket is relatively rigid, compared to other residues within the NiRAN interface domain, as indicated by the atomic fluctuations data shown in Figure S31. The third pocket (pocket 3) is located close to the active site in a cavity between the primer and template strands (see Figures 1 and S27). Comparing these binding sites to the four know HCV allosteric sites^16, 22^ reveals interesting observation (see Figure S32). While pocket 2 and the hairpin pocket identified in the current study seem to be unique to nsp12, due to its distinctive organization, the identified pocket 3 overlaps with the two palm pockets in the HCV polymerase. This comparison also illustrates the clear differences in the architecture between the SARS-CoV-2 and HCV polymerase structures. For example, while the two polymerases share the same right-hand fold with clear fingers, thump and palm subdomains as well as a conserved catalytic site, the nsp12 includes two additional subdomains, namely the NiRAN and its interface subdomain. Furthermore, while the β-hairpin loop is located within the thump domain of the HCV polymerase (see Figure S32), in nsp12, the β-hairpin is located within the n-terminal and in a close proximity to the nsp12 NiRAN domain and very far from the nsp12 thump domain. Our accelerated MD simulations and druggability analysis suggest that pocket 2 is the most druggable among the three identified pockets. Its average volume was around 352.73 Å^3^ with an average LR druggability score ranging between 0.87±0.12 to 0.94±0.04 and CNN scores ranging between 0.93±0.17 to 0.8±0.24, for the accelerated versus classical simulations. These druggability scores are significantly higher than what were observed for the β-hairpin pocket and pocket 3. Residues in pocket 2 also showed very strong correlated dynamics with the catalytic site residues (see Figure 10). Particularly residues P627, N628, M629, L630, S664, M666, V675 and K676 from pocket 2 seem to be highly correlated with the catalytic site resides T680, K621, C622, D623 and S682. This suggests that targeting this pocket can have a significant impact on the catalytic activity of the SARS-CoV-2 polymerase.

As the burden of the novel corona virus, SARS-CoV-2, in the world is immense, there is an urgent and rapid need for innovative and effective treatment options against COVID-19. Furthermore, the short life cycle and virulence of SARS-CoV-2 provide a very short window for effective therapeutic intervention. This implies that focusing on a single viral target/binding site may not be enough to overcome the virus before developing severe symptoms or death, and raises an urgent need to add more druggable options (*e.g.* allosteric sites) within the viral proteins. In this context, we hope the abovementioned findings help expedite the rational discovery of potent and more effective treatments against SARS-CoV-2 infection. We also hope the newly reported allosteric binding sites in this work provide more drug intervention possibilities against the SARS-CoV-2 polymerase beyond conventional nucleoside analogs.

## Conclusion

SARS-CoV-2 is an enveloped, positive-sense RNA virus and has been identified as the cause of COVID-19^1^. Replication of the SARS-CoV-2 genome is a fundamental step in the virus life cycle and inhibiting the SARS-CoV2 replicase machinery has been proven recently as a promising approach in combating the virus^2^. A successful proof of concept is remdesivir, a nucleoside analog (NA) drug that has been repurposed recently against COVID-19^*3*^. Studying the structure and dynamics of the CoV-2 viral replicase can lead to the discovery of important regions and novel druggable hotspots to inhibit the replicase activity. It can also provide deep insights on the hydrogen bond networks formed within the polymerase structure and the important role played by water molecules in mediating these networks.

Earlier efforts to study the CoV-2 polymerase used CoV-1 structures as templates^8-9, 13-14^. These earlier studies were focused mainly on the active site of the polymerase and on understanding the mode of binding of a substrate within this site. In doing so, most of these studies did not consider the dynamicity of the polymerase complex and did not study the hydrogen bond networks to the depth provided in the current work. Additionally, none of these studies investigated the CoV-2 polymerase complex for possible allosteric sites or studied the correlated dynamics within the polymerase structure^8-9, 13-14^.

The recent SARS-CoV-2 polymerase cryo-EM structures^5, 11^ provide a unique opportunity to fill this gap in knowledge using a more accurate structure. Therefore, the focus of the current study was to use these structures to analyze the hydrogen bond networks at the active site and at the template entry in the presence of water, study the binding modes of the nucleotides at the active site, highlight positions for acceptable nucleotides’ substitutions that can be tolerated at different positions within the nascent RNA strand, identify possible allosteric sites within the polymerase structure and study their correlated motion with the catalytic site.

To achieve these goals, we combined classical and accelerated molecular dynamics simulations with principal component analyses, conformational clustering, free energy calculations, binding site identification tools and correlated dynamics. The present work provides an overall view of the nsp7-nsp8-nsp12 replicase complex, with the resolution of a single atom and with the ability to observe small and large conformational changes in the context of a bound RNA. Our findings analyze the hydrogen bond networks at various parts of the polymerase structure and suggest possible nucleotides’ substitutions that can be tolerated by the polymerase complex. We also report here three “druggable” allosteric sites within the nsp12 RdRp that can be targeted by small molecule inhibitors. Our correlated motion analysis reveals a connection between the dynamics within these new identified sites and the polymerase active site, indicating that targeting these sites can impact the catalytic activity of the SARS-CoV-2 polymerase.

## Methods

### Preparation of the nsp12-nsp8-nsp7-RNA complex

The recent SARS-CoV-2 polymerase cryo-electron microscopy structures (PDB IDs: 6M71^5^ and 7BV2^11^) were used to construct the nsp7-nsp8-nsp12-RNA assembly. In this context, the 7BV2 structure was used to build the initial complex, as it included a bound RNA and an incorporated remdesivir drug within the RNA primer chain. While this structure provided a good starting point to model the nsp12-RNA complex, it suffered from various anomalies. For example, the trio-aspartate residues D618, D670 and D671 as well as the two magnesium ions were not properly positioned at their coordination conformation. Furthermore, this structure lacked a complete nsp8 monomer that has been reported to interact with the nsp7 protein on the surface of nsp12 [ref]. This structure also lacked various residues in nsp7, nsp8 and snp12 (highlighted in blue in Figure S33) (See supplementary data). To incorporate the missing nsp8 monomer, we used the SARS-CoV-2 polymerase structure (PDB ID: 6M71), which included a nsp8 dimer, with one of the nsp8 monomers was correctly positioned to interact with nsp7. Missing residues in all three proteins were added using homology modelling tools. The full sequences for the three proteins were obtained from the 7BV2 related files as deposited in the Protein Data Bank. For selecting the best templates to incorporate the missing residues, sequence alignments were performed in BLASTP suite using BLOSUM62 algorithm. Through this process BLOSUM62 identified chain A of the SARS-CoV-2 polymerase complex (PDB ID: 6M71) as the best template for nsp12 and chain C of the same structure for nsp8. On the other hand, it identified the bound nsp7 in the nsp7-nsp8 complex (PDB ID: 6WIQ) as the best template for nsp7. Several homology models were built using I-TASSER, Modeller Suite and SWISS-MODEL. The highest quality model based on QMEAN scoring function, Ramachandran plots and ERRAT error values was ultimately chosen (from SWISS-MODEL). It is worth mentioning that our use of homology modelling was not meant to construct the whole structures of nsp7, nsp8 and nsp12 from scratch, as these structures were properly determined at a 2.5Å resolution and, overall, they were correctly folded, particularly at the proteins’ interfaces (*i.e.* 6M71) and at the interaction sites of nsp12 and the bound RNA (*i.e.* 7BV2). Our use of homology modelling was mainly focused on incorporating missing residues that formed critical discontinued regions within the protein structures. Therefore, our final constructed nsp12-nsp7-nsp8-RNA complex included the continues residues’ sequences V30 to E919 for nsp12; S1 to E73 for nsp7; and M67 to N192 for NSP8. Our final structure also included the RNA primer sequence (GCUAUGUGAGAUUAAGUUAU) and the RNA template sequence (UUUUUUUUUUAUAA CU UAAUCUCACAUAGC).

### Preparation of Bound and Incorporated Substrates

The active site of the SARS-CoV-2 polymerase is formed by the conserved polymerase motifs A-G in its palm domain, similar to other RNA polymerases^23^. Given this conserved nature of the polymerase among RNA viruses, we used the structure of the hepatitis C virus (HCV) polymerase, NS5B, in complex with pp-sofosbuvir (PDB ID: 4WTG)^24^ to guide our modelling of the metal coordination within the SARS-CoV-2 nsp12 active site. To do that, we used the software MOE^25^ to superimpose the SARS-CoV-2 and HCV polymerase structures and to manually orient the three aspartate residues D618, D670 and D871 (corresponding to D220, D318 and D319 in HCV) to be properly positioned to coordinate the two magnesium ions. It is worth mentioning that the nsp12 SARS-CoV-2 structure (PDB ID: 7BV2) contained three magnesium ions. Only two of these ions were kept in our model and were positioned at their correct coordination sites, with guidance from the HCV NS5B structure. To accurately position the incoming nucleotide triphosphate (NTP) within the nsp12 active site, we first removed the incorporated remdesivir from the primer chain in the nsp12 complex (PDB ID: 7BV2) and used the bound sofosbuvir diphosphate in the HCV NS5B structure (PDB ID: 4WTG) as a guidance to position remdesivir triphosphate. We then used this model to create four additional complexes for the four possible types of RNA nucleotides (i.e. ATP, CTP, GTP and UTP). Therefore, we created five different active site-bound systems in total. To construct models for the incorporated nucleotides within the primer chain, we used the incorporated remdesivir in the 7BV2 structure as a starting point and modified its bound chemical structure to other nucleotides’ structures (*i.e.* A, C, G and U) using the MOE software^25^. We also modified the template complementary nucleotide (at position i) to the correct base-paring matching structure for each nucleotide. Therefore, in addition to the five bound systems, we have also constructed four polymerase structures representing the incorporation of the four different types of RNA nucleotides in the RNA chain.

### Protonation States Adjustment

Protonation states of all ionizable residues for the nine systems described above were calculated using the program PDB2PQR at pH 7^26^. Protein residues were then visualized and carefully inspected to ensure that their correct protonation states were predicted by PDB2PQR. For example, the three catalytic aspartate residues (D220, D318 and D319) were deprotonated to provide a negatively charged environment for the two positively charged magnesium ions. The nearby aspartate, D623 was also deprotonated. Lysine residues K798, K551, K621as well as the serine residue S759 were all protonated. All ZINC chelating cysteine residues (*i.e.* C301-C306 and C487-C645) were deprotonated and their corresponding histidine residues (H295 and H642) were protonated at their epsilon positions.

### Classical MD Simulations

The nine protein-RNA complexes, described above, were subjected to classical MD simulations at a temperature of 300 K, employing the software NAMD^27^ at physiological pH (pH 7) and using the all-hydrogen AMBER99SB force field. Charges for the bound remdesivir and four nucleotides were calculated at the semi-empirical AM1-BCC procedures in Antechamber ^28^ and their force field parameters were prepared according to the generalized amber force field (GAFF) ^29^. All systems were immersed in the center of a TIP3P water cube after adjusting the protonation states of proteins, RNA and bound ligands. The cube dimensions were chosen to provide at least a 14 Å buffer of water molecules around each complex. The total initial charge for the five nucleotide-bound protein-RNA complexes were −42 and were −38 for the four nucleotide-free complexes. To neutralize and prepare the protein-RNA complexes with a physiological ionic concentration (*i.e.* 150 mM), chloride and sodium ions were introduced by replacing water molecules having the highest electrostatic energies on their oxygen atoms. In total we added 181 sodium and 140 chloride ions to the nucleotide-bound systems and added 178 sodium and 140 chloride ions to the nucleotide-free systems. Following an MD protocol similar to our earlier published work [ref], the fully solvated-ionized systems were then minimized and subsequently heated gradually to the simulation temperature with heavy restraints placed on all backbone atoms of the proteins and RNA as well as on the bound ligands, metal ions and their coordinating amino acids. Following heating, the systems were equilibrated using periodic boundary conditions for 200 ps and energy restraints were reduced to zero in successive steps of the MD simulations. The simulations were then continued for ∼100 ns and atomic coordinates were collected at intervals of 0.1 ps for subsequent analyses.

### Principal component analysis

The active site residues for all nine systems was subjected to principal component analysis (PCA). This analysis transforms the original space of correlated variables from the MD trajectories into a reduced space of independent variables comprising the essential dynamics of each system^39^. We focused in this analysis on the bound/incorporated nucleotide, the base paring RNA nucleotide (+1), the two magnesium ions, and all heavy atoms of 18 nsp12 residues forming the catalytic site (*i.e.* K551, R555, D618, P620, K621, C622, D623, T680, S682, G683, T687, A688, N691, S759, D760, D761 and K798). To perform PCA on these atoms, we followed our earlier work^30-31^, by first fitting the entire MD trajectory for each system to a reference structure (*i.e.* the minimized complex). This was important in order to remove all rotations and translations. The covariance matrix was then calculated from the Cartesian atomic coordinates of the residues mentioned above as follows: 

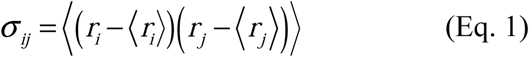

The eigenvectors of the covariance matrix constitute the essential vectors of the motion. It is generally accepted that the larger an eigenvalue, the more important its corresponding eigenvector in the collective motion. To ensure completeness of conformational sampling of the active site residues and the bound nucleotides, we projected the whole trajectory for these residues, for each system, on the first and second eigenvectors as well as on the second and third eigenvectors.

### Clustering Analysis Protocol

To analyze the conformational dynamics of the bound nucleotides and to extract their dominant mode(s) of binding within the active site, we carried out RMSD conformational clustering on the whole MD trajectories. We used the average-linkage algorithm as implemented in the CPPTRAJ utility of AMBER18 using cluster counts ranging from 2 to 100 clusters. The application of the average-linkage algorithm, among other clustering algorithms, to MD trajectories has been validated in recent studies.^32^. To perform the clustering analysis, we focused on all atoms of the bound/incorporated nucleotide by first performing an RMSD fitting of these atoms to their minimized initial structure conformation in order to remove overall rotation and translation. These atoms were clustered into groups of similar conformations and the optimal numbers of clusters were predicted after evaluation of the Davies-Bouldin index (DBI)^33^ and the “elbow criterion”^32^. A high-quality clustering scheme is expected when a local minimum in the DBI values coincides with a plateau in the percentage of variance explained by the data (SSR/SST), while varying the number of clusters.^32^ The centroid of each cluster, the structure having the smallest RMSD to all members of the cluster, was then selected as the cluster representative structure and the dominant structures, representing the dominant clusters, were then used for further analysis.

### Binding Site Identification Analysis

Trajectories from the MD simulations were used to identify potential allosteric binding sites within the nsp12-nsp7-nsp8-RNA polymerase assembly. We used three different pocket identification tools to search for these sites. These tools included fPocket^34^ COACH^35^ and Site Finder as implemented in MOE^25^. Consensus between the three tools, size and druggability of the identified sites were the main criterion for the final selection sites to focus on.

### Hydrogen Bond Analysis

Hydrogen bond analyses were performed on the whole trajectory of each of the nine MD systems using the CPPTRAJ tool as implemented in AMBER 18 and visualization of the hydrogen bonds was performed using the visualization software VMD^36^. A hydrogen bond was defined by a cut-off distance of 3.5Å between a donor and acceptor atom and an absolute angular deviation below 60° from linearity.

### Water Hydration Shell Analysis

To determine the available space around the bound/incorporated nucleotides within the catalytic site, we counted the number of water molecules representing the first and second solvation shells in their vicinity. To do that, we used the CPPTRAJ of AMBER 18 to count the number of water molecules within 2Å and 4Å of their structures. We used the same analysis to predict the size of the identified allosteric sites throughout the whole MD simulations.

### Electrostatics Potential Calculations

Electrostatic potentials for nsp12-nsp8-nsp7-RNA complexes were calculated using the APBS program^37^ and mapped onto a reduced molecular surface with the VMD visualization program^38^. The solute (proteins, RNA and bound/incorporated nucleotides) were treated as a low dielectric medium (*ε*_*in*_ = 1) surrounded by a high dielectric solvent (*ε*_*in*_ = 78.54 for water). The ionic strength was set to 0.1 M. The low-dielectric region of the protein was defined as the region inaccessible to contact by a 1.4 Å sphere rolling over the molecular surface, defined by atomic co-ordinates of the MD structure and vdW radii taken from the all-hydrogen AMBER99SB force field. The electrostatic potential calculations employed a 125 × 125 × 125 grid with a spacing of 1 Å.

### Nonbonded energy analysis

All energy calculations for the nonbonded interaction energy analysis were performed using the NAMD Energy Plugin, Version 1.4 as implemented in the VMD visualization software. All energy calculations were performed using the all-hydrogen AMBER99SB force field and using the same parameters and periodic boundary conditions that were used to run the classical MD simulations described above.

### Accelerated MD Simulations

To study the polymerase system in a time and length scale beyond the limits of classical MD simulations. To do that, we used accelerated MD simulations (AMDS) module as implemented in AMBER, which applied a boosting potential with a positive value (ΔV(r)) when the potential energy falls below a certain energy threshold^39^. We have applied the dual boosting simulation scheme (iamd = 3), where the boosting potential is applied to the whole system, and an additional boosting is applied specifically to the torsion angles. The method has been usefully applied in a number of studies, where a brute force classical simulation is not sufficient to study the conformational dynamics of a bio-molecular system^40-42^. For more discussion about the method, readers are encouraged to consult our recently published work, where the method was used to study the conformational dynamics of the programmed death ligand 1 protein (PD-L1)^43^. Initial system preparation for AMDS was performed using a FREE CTP system, similar to what has been descibed above. That is, the solvated system was minimized in four steps: First, a harmonic potential of 100 kcal/mol/Å^2^ was applied to all protein/RNA heavy atoms. In the following minimization stages, the restraining potential was reduced to 50 kcal/mol/Å^2^, 5 kcal/mol/Å^2^ and 0 kcal/mol/Å^2^. At each step, minimization was performed by the steepest descent method for the first 1000 steps and the conjugated gradient method for the rest of the 5000 total minimization steps. Each system was then gradually heated in the NVT ensemble from 0 to 300 K in 100 ps using a Langevin thermostat with a coupling coefficient of 1.0/ps, and a force constant of 5.0 kcal/mol/Å^2^ on the protein/RNA heavy atoms, using a 1 fs integration time step. And then additional two rounds of MD (50 ps each at 300 K) were performed with decreasing heavy atoms restraint weights reduced from 1, 0.5 to 0.1 kcal/mol/Å^2^. Finally, each system was again equilibrated for 10 ns by releasing all restrains. This was followed by 100 ns accelerated MD simulation, and in parallel, an additional 100 ns classical MD simulation using the AMBER MD simulations package was used as a reference for the AMDS. For each simulation, the first 20 ns of the simulation was used as an equilibration phase and was discarded from the analysis. All equilibration and production simulations were performed at 300 K with Berendsen temperature coupling ^44^ and 1 atm with isotropic molecule-based scaling and a 2fs integration time step. Long-range Coulombic interactions were handled using the particle mesh Ewald (PME) method ^45^. The cut-off distance for the electrostatic and van der Waals (vdW) energy term was set at 9.0 Å. To avoid edge effects in all calculations, periodic boundary conditions were applied. To allow for an integration time step of 2 fs, the SHAKE algorithm ^46^was applied for all bonds involving hydrogen atoms. Coordinate trajectories were recorded every 4 ps throughout all production runs.

### Druggability Assessment of the Identified Pockets

500 protein conformations from the last 80 ns of the accelerated MD simulation were samples and used to confirm the existence of the identified pockets and to evaluate their druggability. Sampled protein conformations were used as inputs for TRAnsient Pockets in Proteins (TRAPP) – a powerful machine-learning tool for binding pocket detection and druggability assessment^47-48^. In its most recent implementation, TRAPP offers the possibility of assessing the druggability of the detected pockets using either a logistic regression model (TRAPP-LR) and/or a convolutional neural network model (TRAPP-CNN), deep learning model^49^. These models were trained using publicly available and authors-collected datasets. In their assessment, the models achieved superior performance in identifying correct druggable pockets for the test dataset, even in challenging protein targets. Both TRAPP-LR and TRAPP-CNN use the geometrical as well as the physicochemical properties (*e.g.* overall hydrophobicity, hydrogen bond donors and acceptors) gathered over the sampled protein conformation to calculate druggability scores for the corresponding sampled protein conformers. In performing this analysis, all sampled protein conformations were fitted on the heavy atoms of the starting protein coordinates considered for our analysis (the first frame of the last 80 ns of the MD simulation). We have focused on the three potential pockets identified from classical MD simulations (see results), namely the β-hairpin pocket, pocket 2 at the NiRAN interface and pocket 3, close to the active site. The residues center of mass for each pocket were used a geometric center for the corresponding pocket search. Pocket volumes and druggability scores were collected and analyzed (See results).

### Correlated Dynamics

A residues correlation analysis between the residues constituting each pocket and the catalytic site was performed using the LMI algorithm implemented in the BIO3d R library ^50^. Correlation analysis was performed for 20,000 snapshots extracted from the last 80 ns production simulation and was performed for the classical as well as the accelerated MD simulations.

## Contributions

KHB conceived the study. KB and MA developed and implemented the computational studies with contributions from YT and MH. KHB drafted the manuscript. MA, YT and MH edited the manuscript and analyzed the data.

## Acknowledgements

KB acknowledges NSERC Discovery grant and funding from Alberta Cancer Foundation. We acknowledge the HPC computing facilities provided by Compute Canada.

## Conflict of Interests

None

## Notes

### Competing Interest Statement

The authors have declared no competing interest.

