## Supporting Information for "A “Deep Dive” into the SARS-Cov-2 Polymerase Assembly: Identifying Novel Allosteric Sites and Analyzing the Hydrogen Bond Networks and Correlated Dynamics"

**Supplementary Data**

| **NTP(i)** | **vdW (kcal/mole)** | **ELE (kcal/mole)** | **Total (kcal/mole)** |
| --- | --- | --- | --- |
| Remdesivir | -0.3 | -100.7 | -101.0 |
| CTP | -0.2 | -98.2 | -98.4 |
| GTP | 5.8 | -6.0 | -0.2 |
| UTP | 37.1 | -136.0 | -98.9 |

**Table S1.** The total vdW, electrostatic energy and total non-bonded interactions for remdesivir and the three nucleotides, UTP, CTP and GTP, relative to the ATP energies.

| **RNA** | **vdW (kcal/mole)** | **ELE (kcal/mole)** | **Total (kcal/mole)** |
| --- | --- | --- | --- |
| Remdesivir | 11.0 | -92.2 | -81.2 |
| CTP | 20.4 | -54.0 | -33.5 |
| GTP | 5.8 | +5.7 | +85.8 |
| UTP | 37.1 | +17.0 | +12.9 |

**Table S2.** The total vdW, electrostatic energy and total non-bonded interactions for the bound RNA strands for remdesivir and the three nucleotides, UTP, CTP and GTP, relative to the ATP energies.

**Figure S1.** Structural assessment for the nsp12 initial structure.

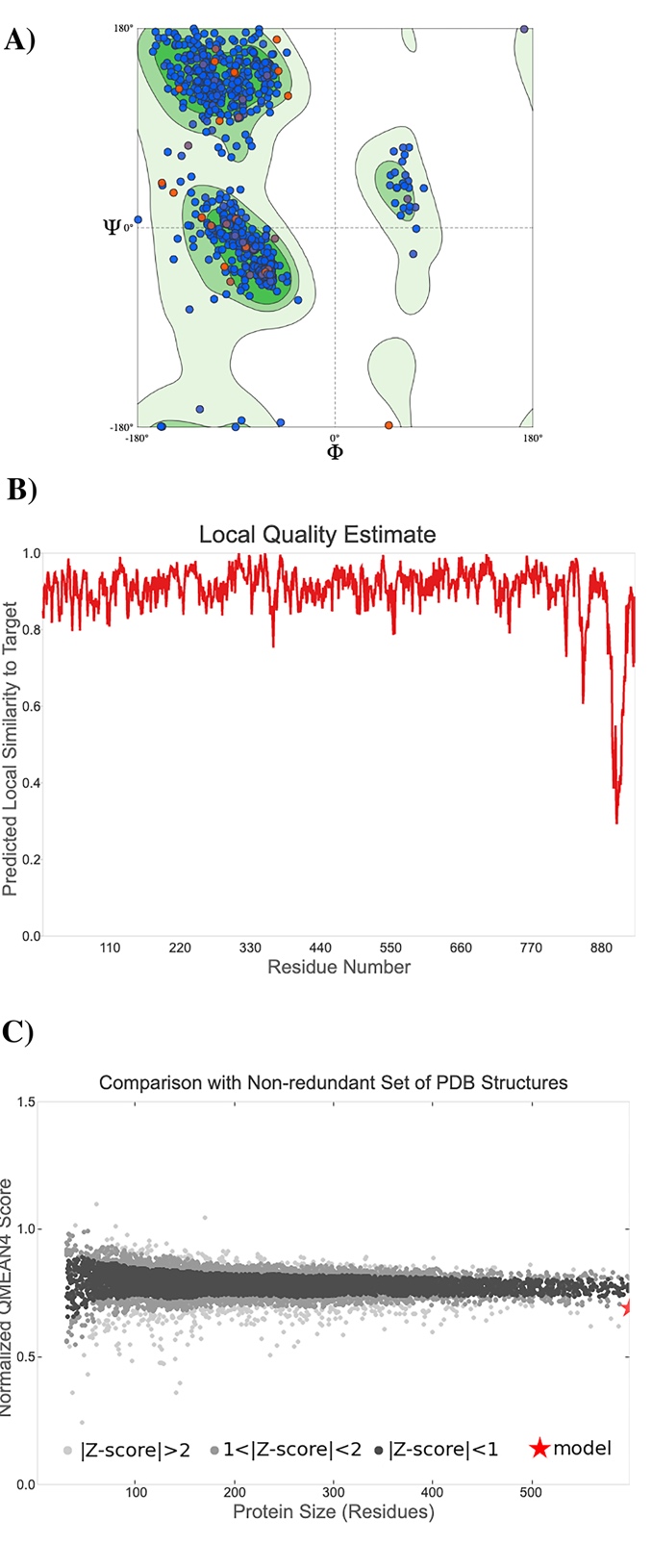

**Figure S2.** Structural assessment for the nsp8 initial structure.

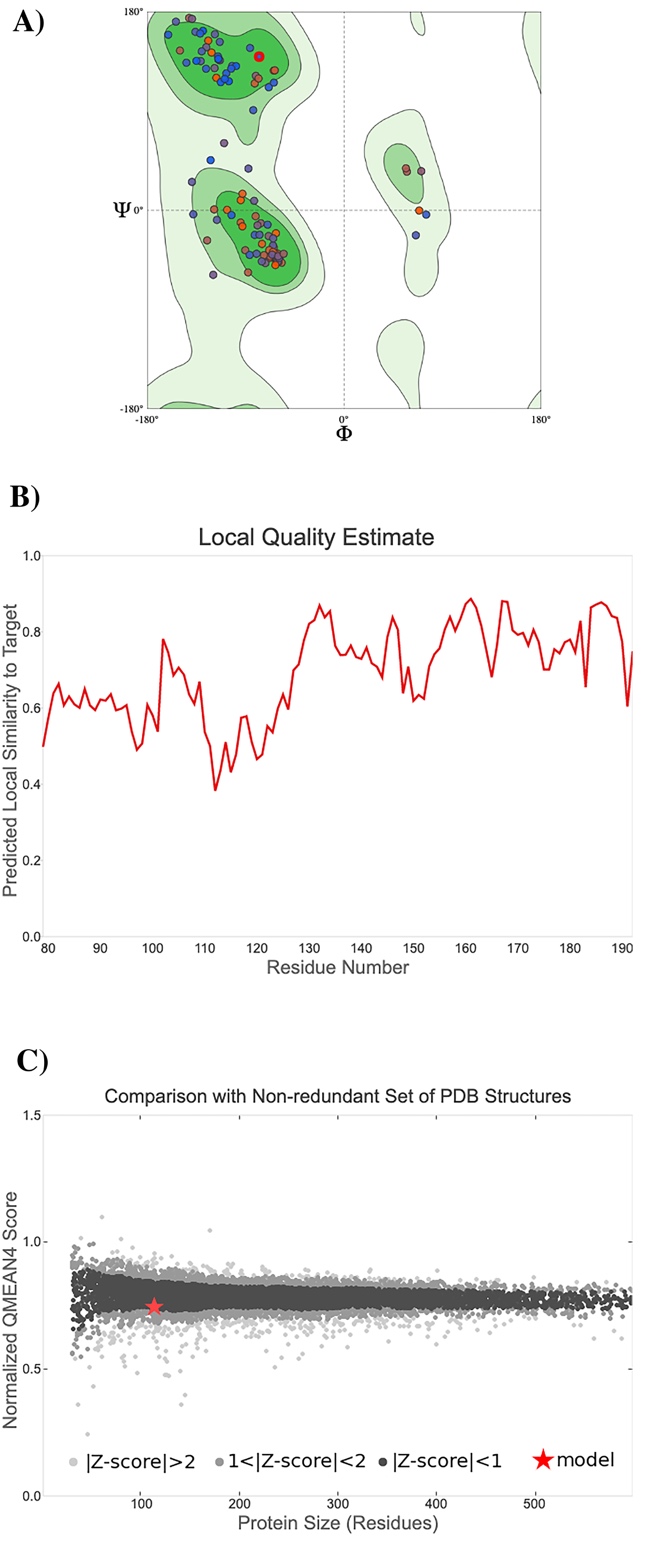

**Figure S3.** Structural assessment for the nsp7 initial structure.

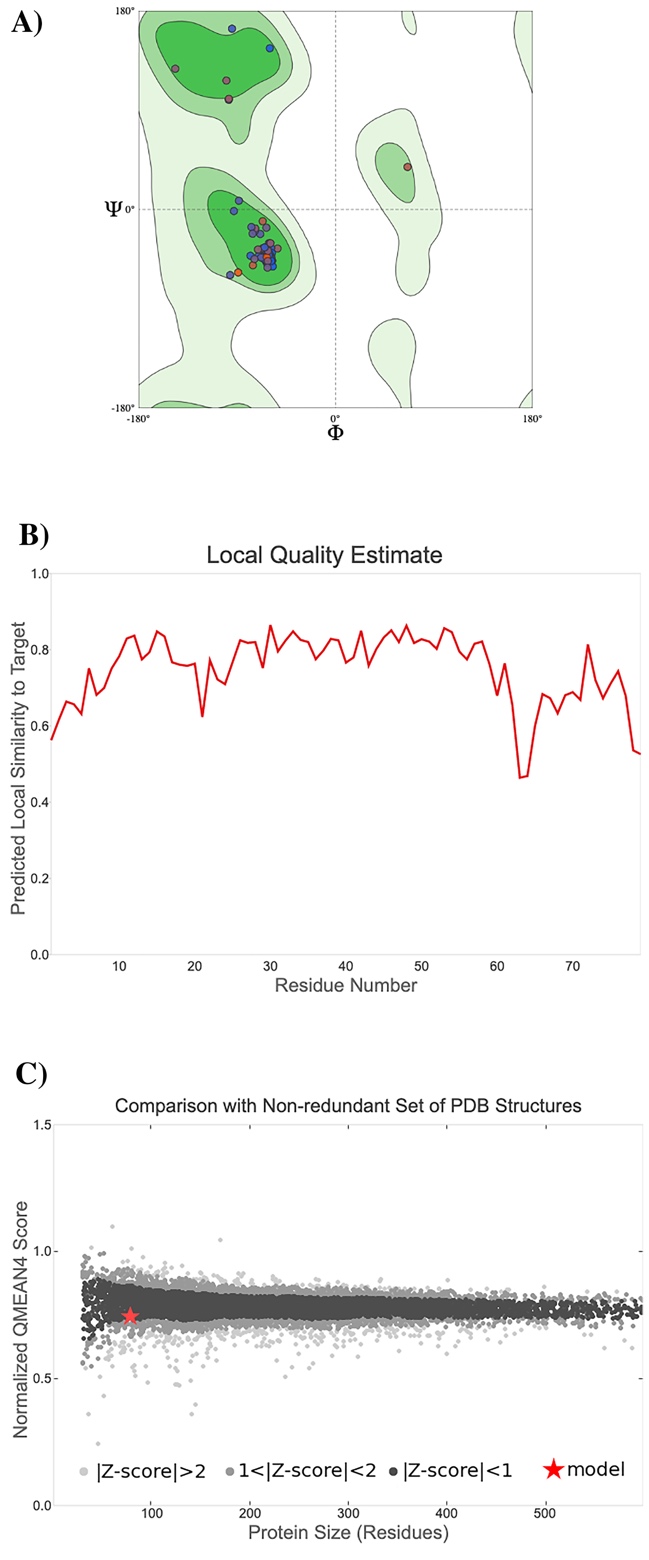

**Figure S4.** Atomic fluctuations for the bound NTPs, the two chelated zinc ions and the two magnesium ions for the BOUND systems.

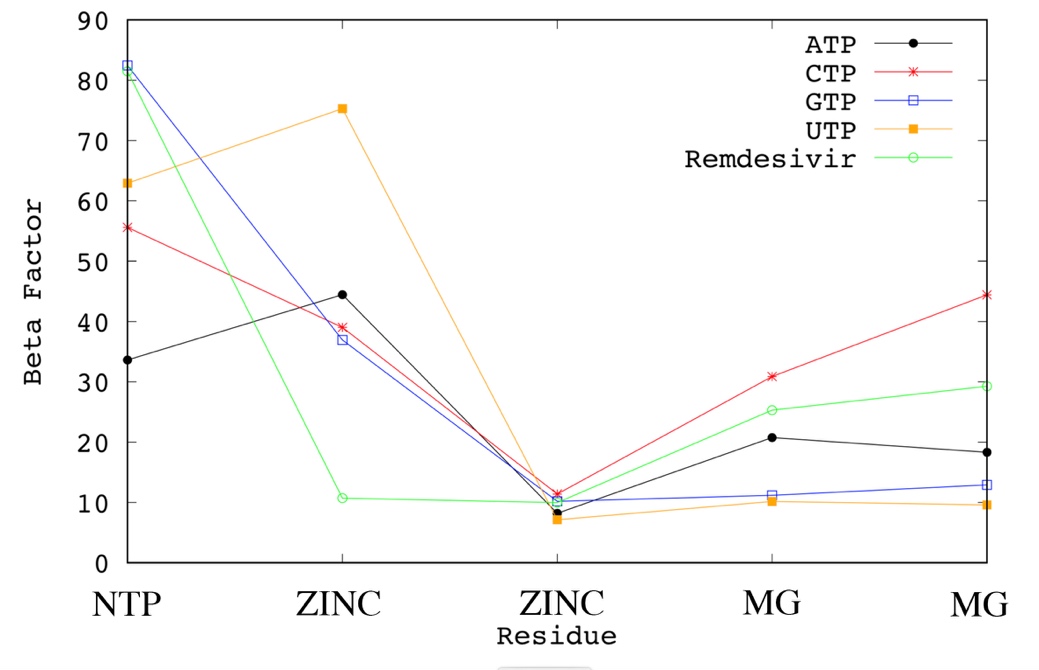

**Figure S5.** RMSD and Atomic fluctuations for nsp7 for the BOUND (A, B) and FREE (C, D).

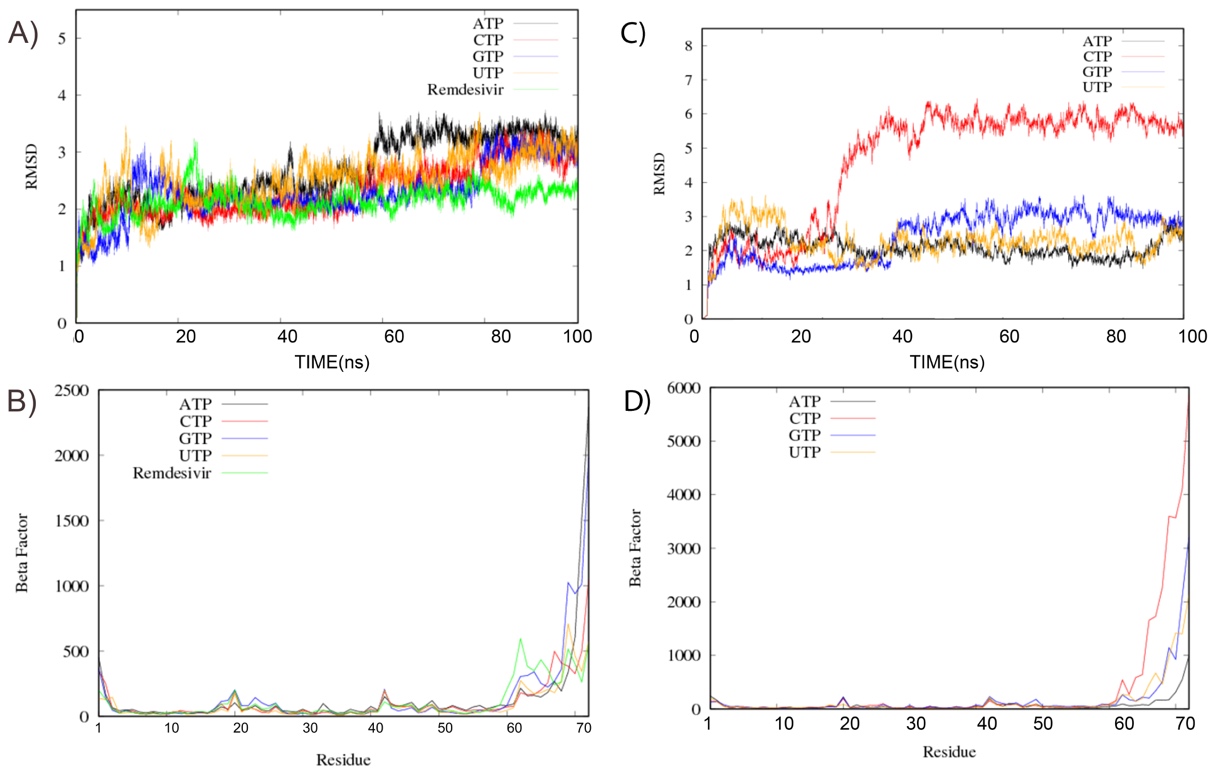

**Figure S6.** RMSD and Atomic fluctuations for nsp8-1 for the BOUND (A, B) and FREE (C, D).

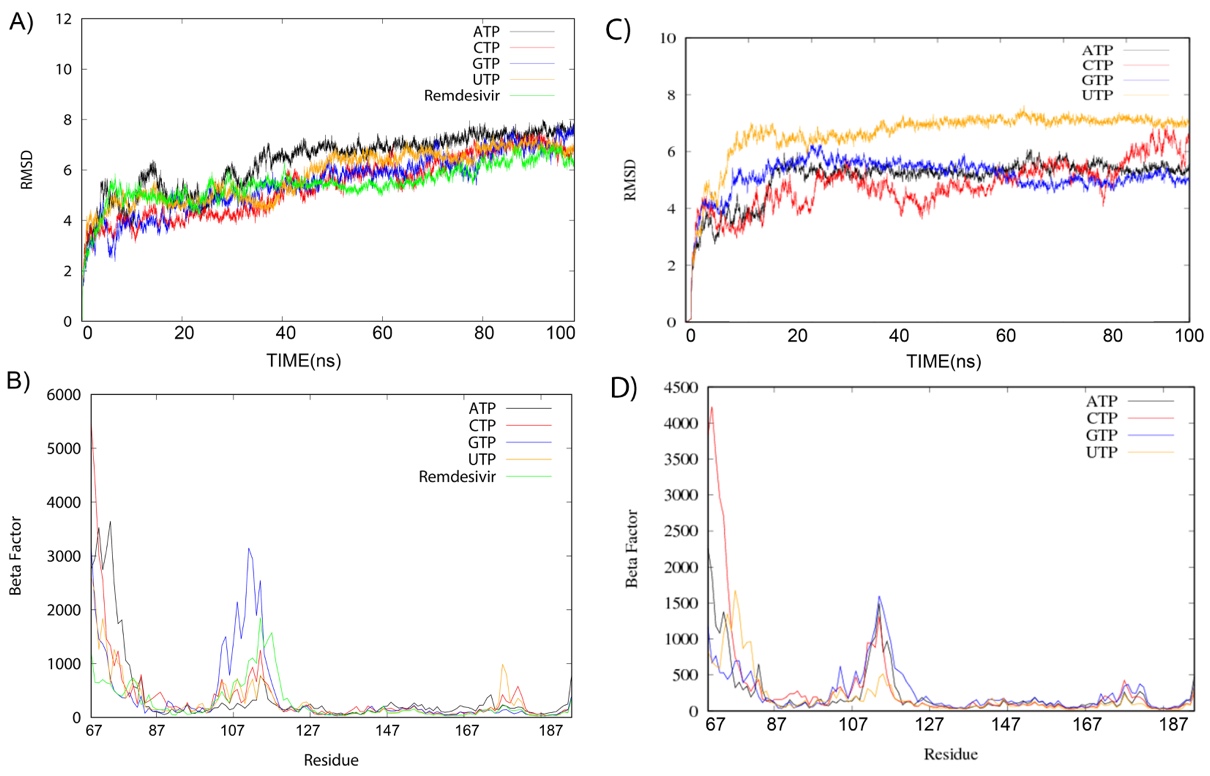

**Figure S7.** RMSD and Atomic fluctuations for nsp8-2 for the BOUND (A, B) and FREE (C, D).

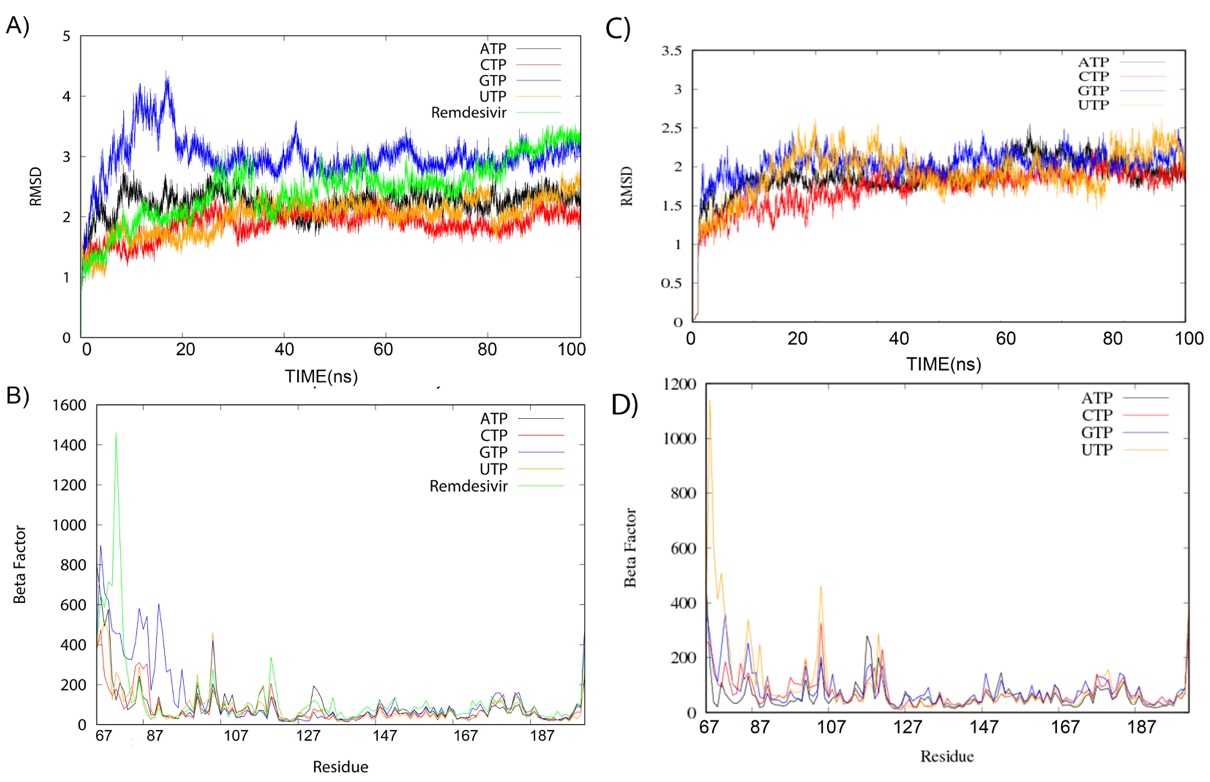

**Figure S8.** Flexible regions within nsp7 (highlighted in yellow), nsp8-1 & nsp8-2 (highlighted in green).

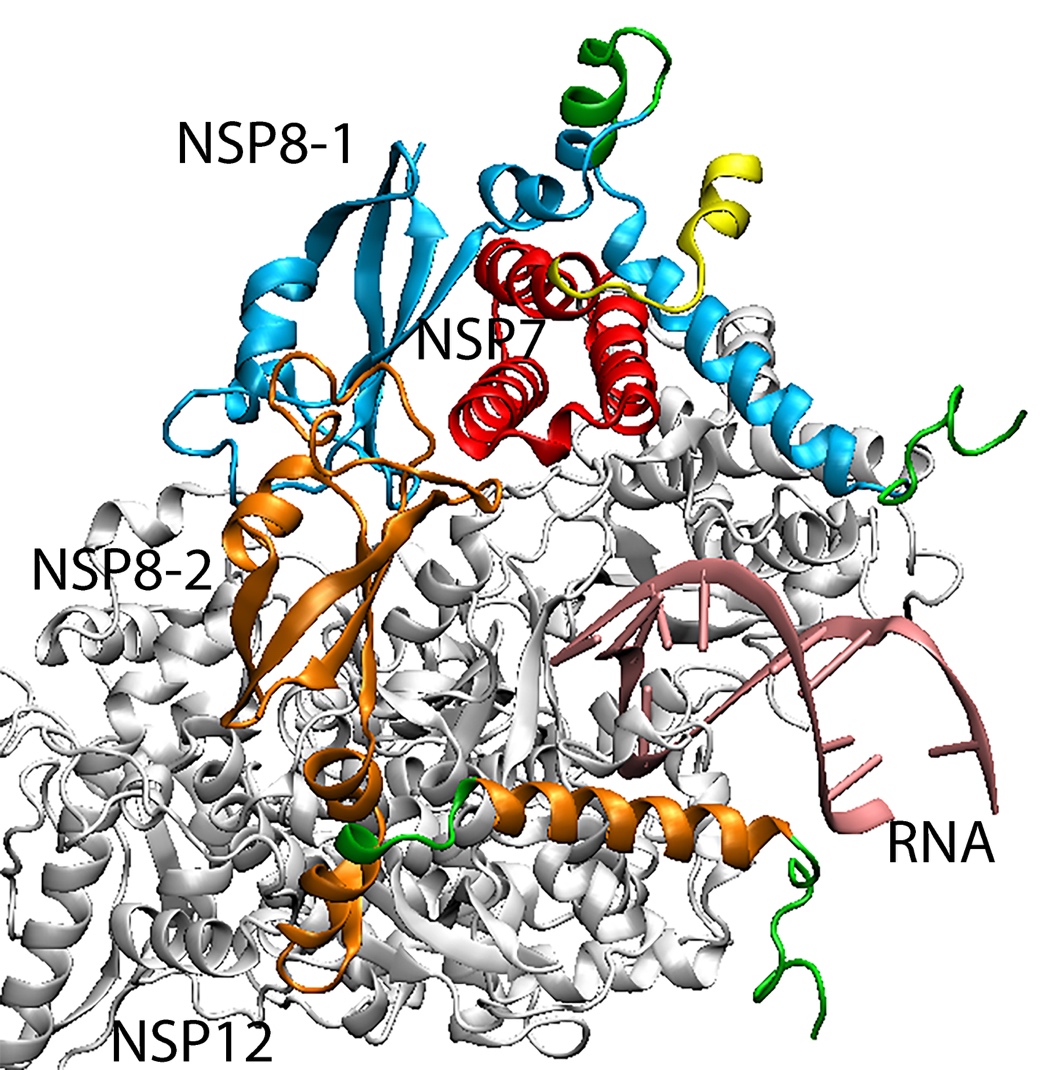

**Figure S9.** RMSD and Atomic fluctuations for the nascent RNA strand for the BOUND (A, B) and FREE (C, D).

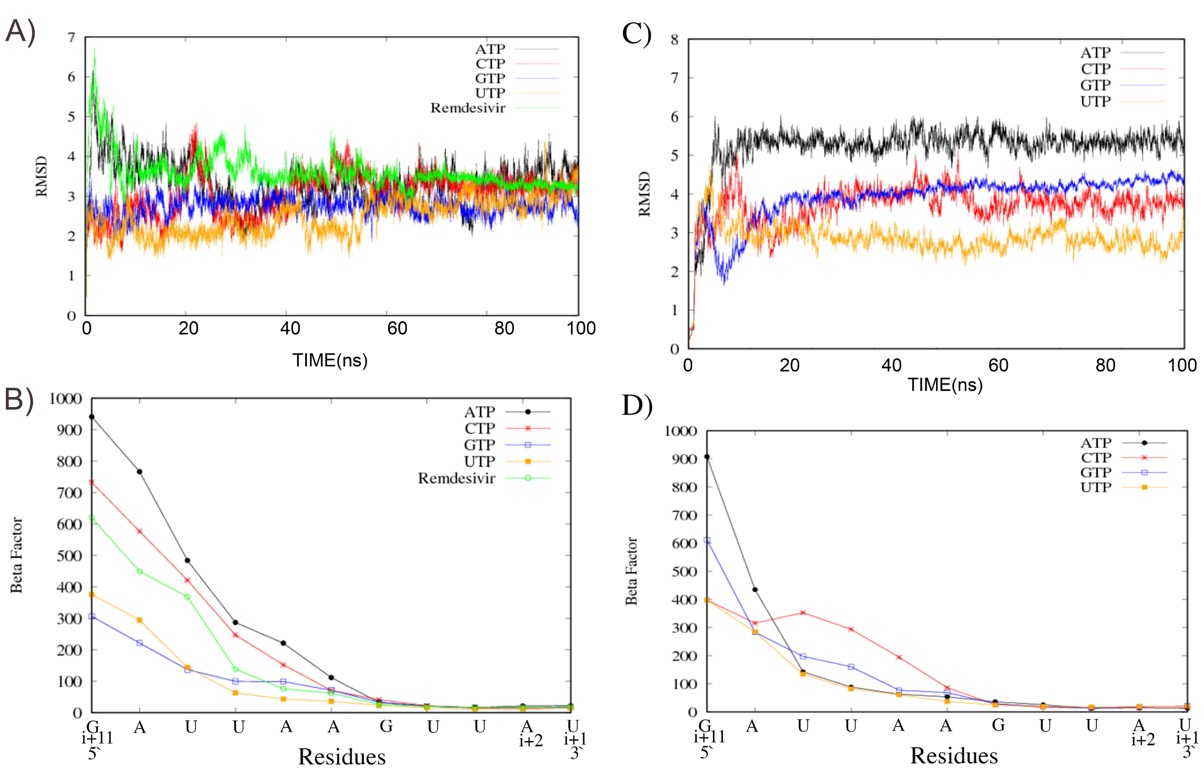

**Figure S10.** RMSD and Atomic fluctuations for the template RNA strand for the BOUND (A, B) and FREE (C, D).

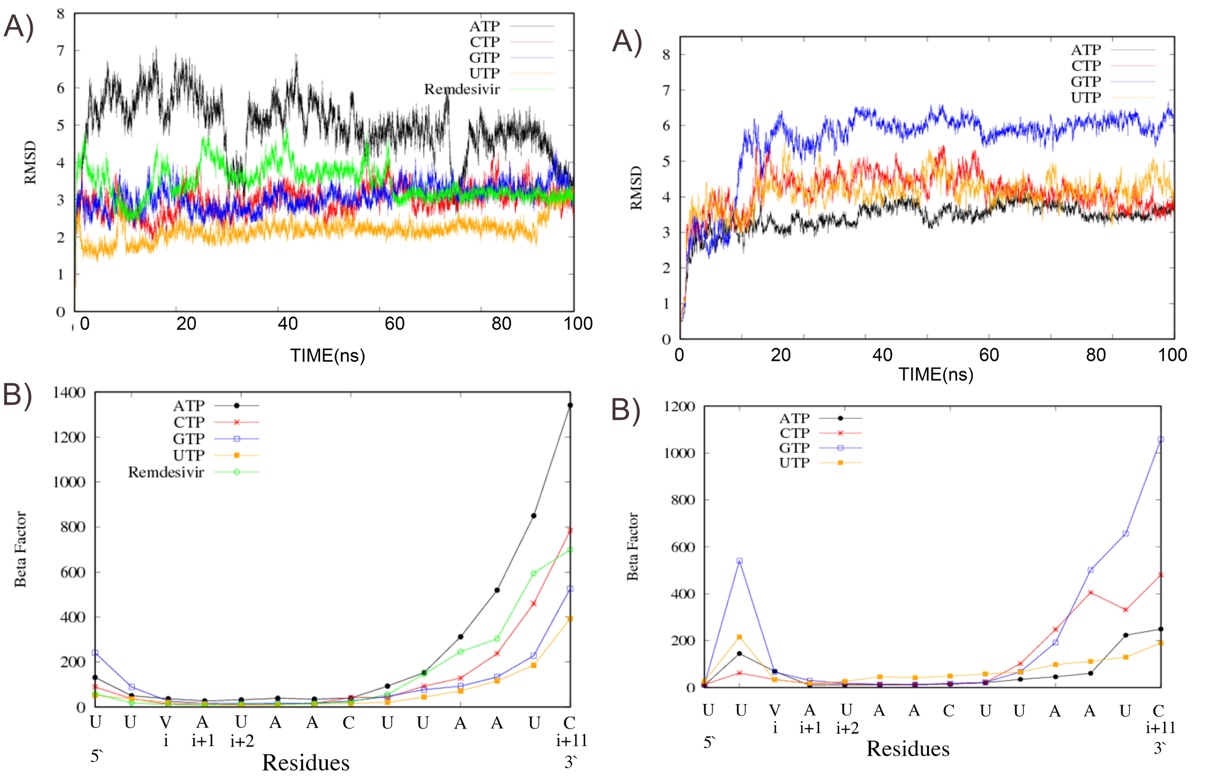

**Figure S11.** Principal component analysis (PCA) on the catalytic site for FREE (C, D) systems.

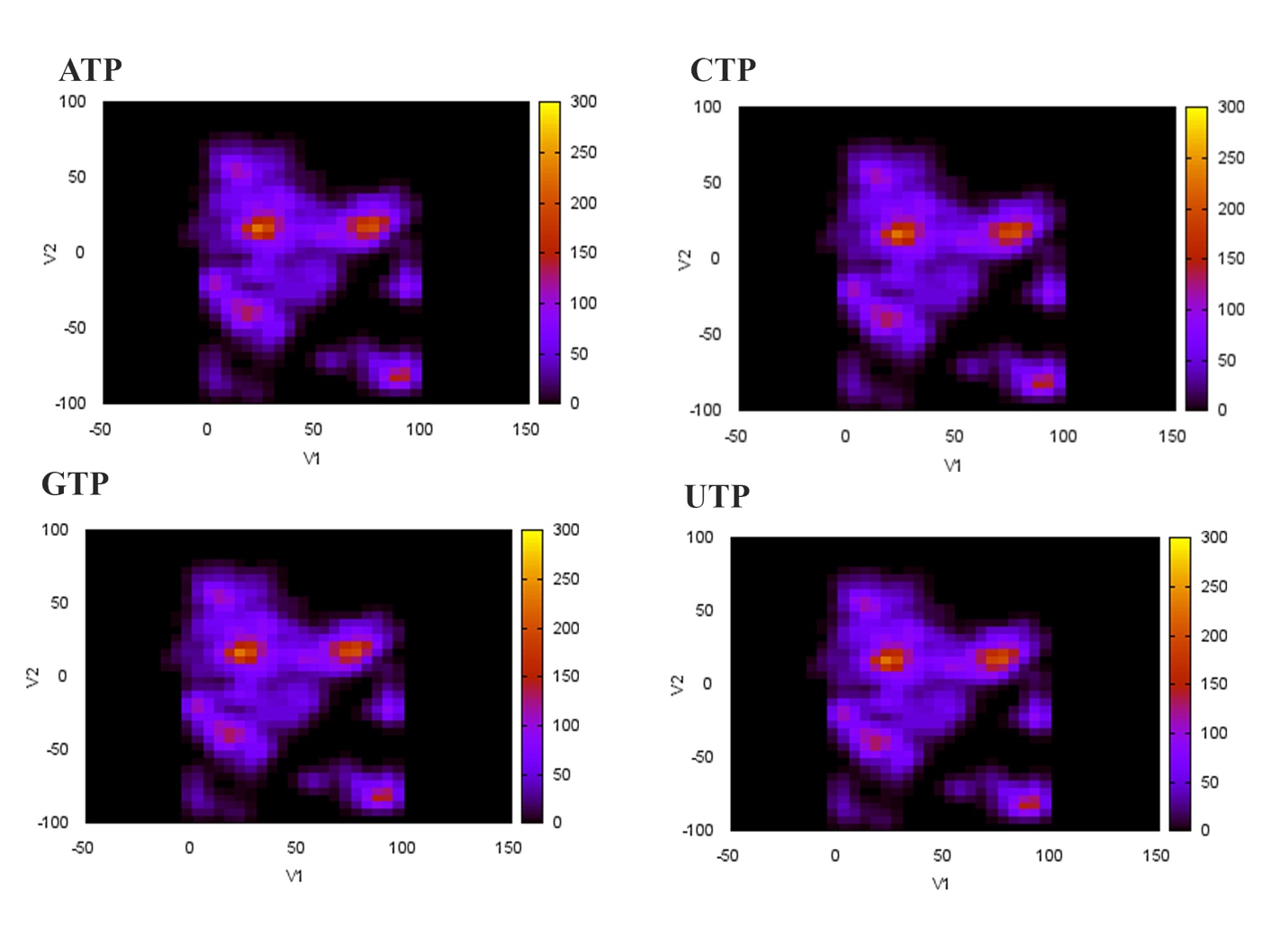

**Figure S12** Conformational clustering of the bound nucleotides’ trajectories for CTP.

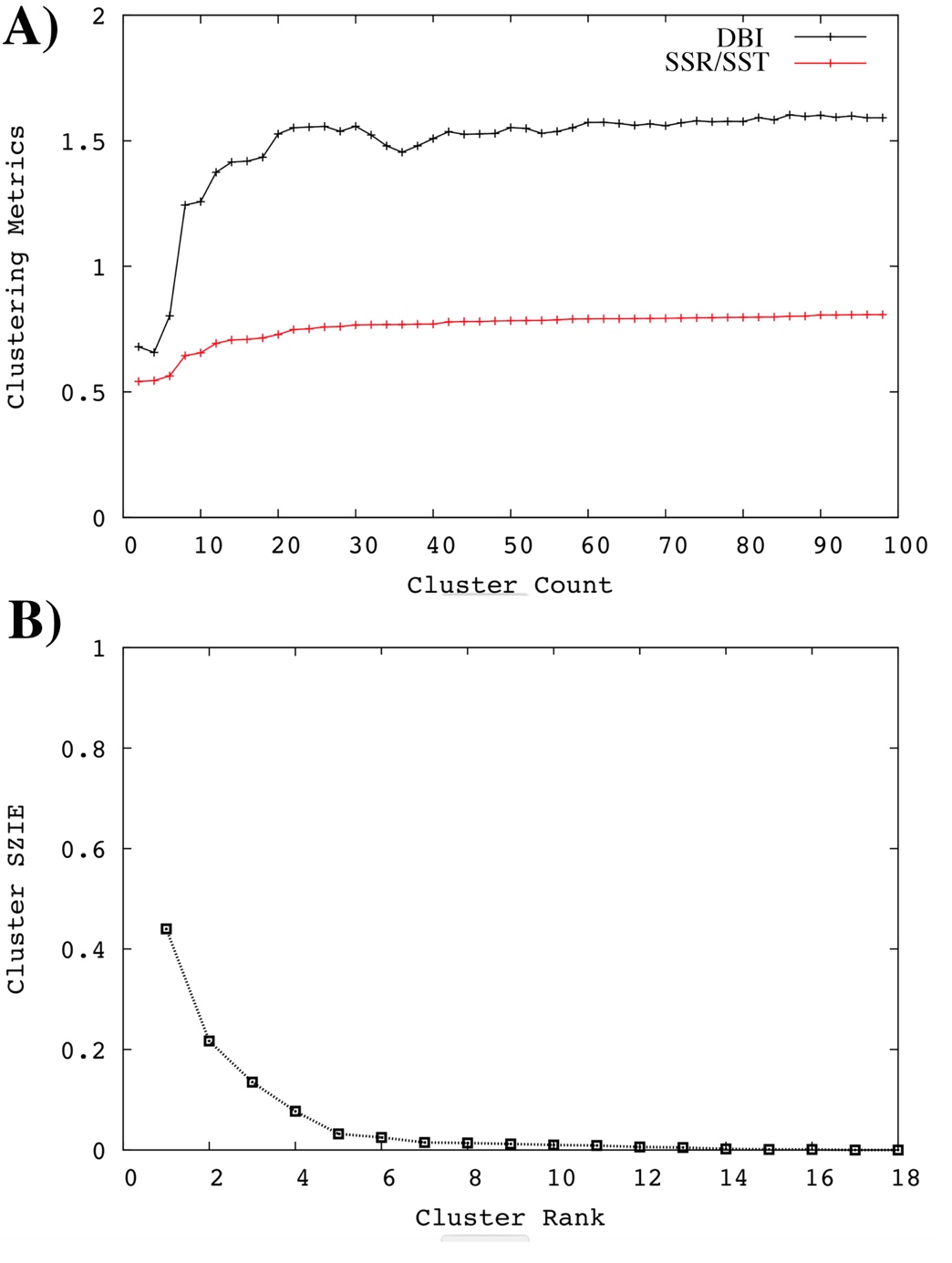

**Figure S13** Conformational clustering of the bound nucleotides’ trajectories for GTP.

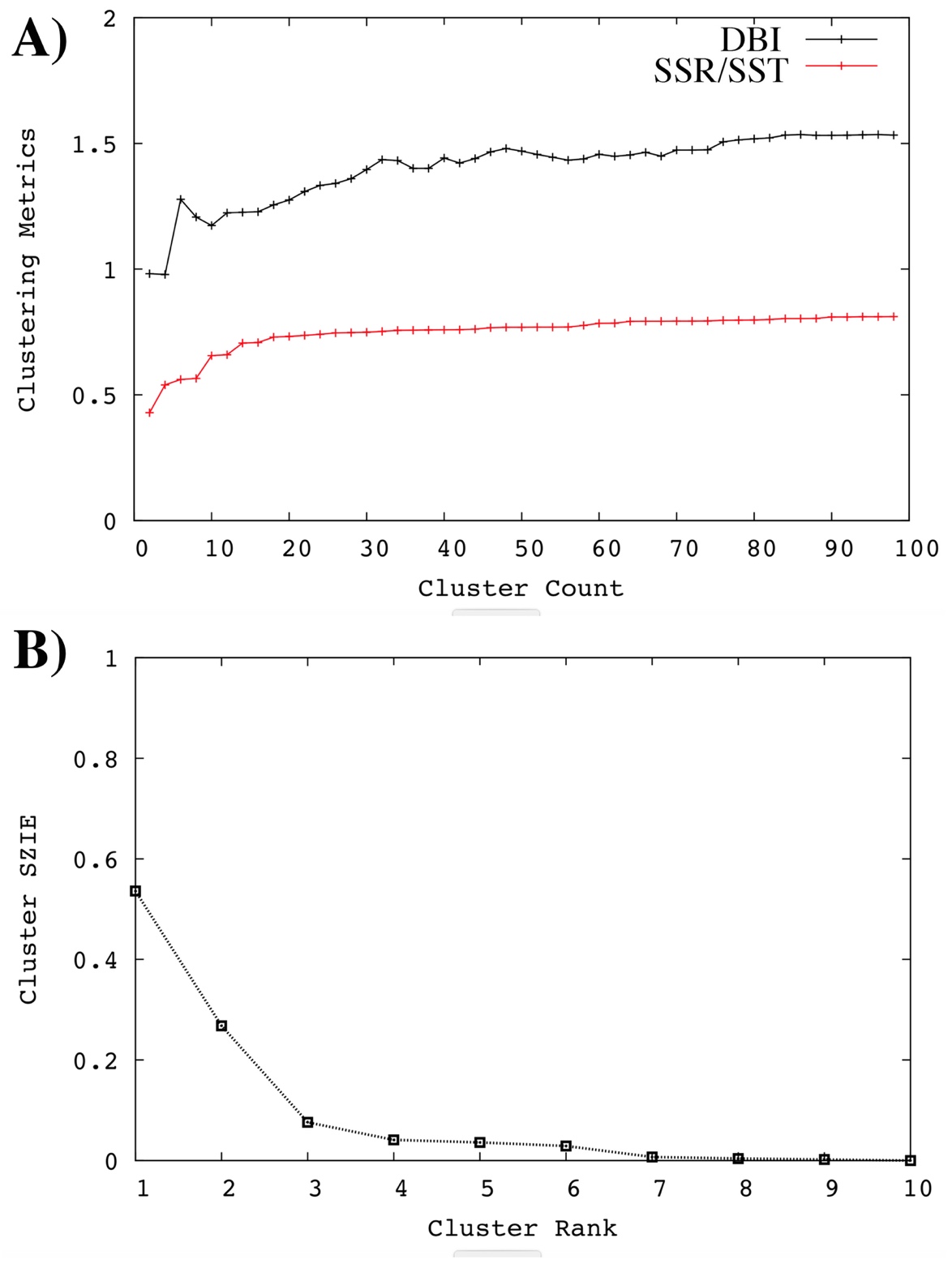

**Figure S14** Conformational clustering of the bound nucleotides’ trajectories for UTP.

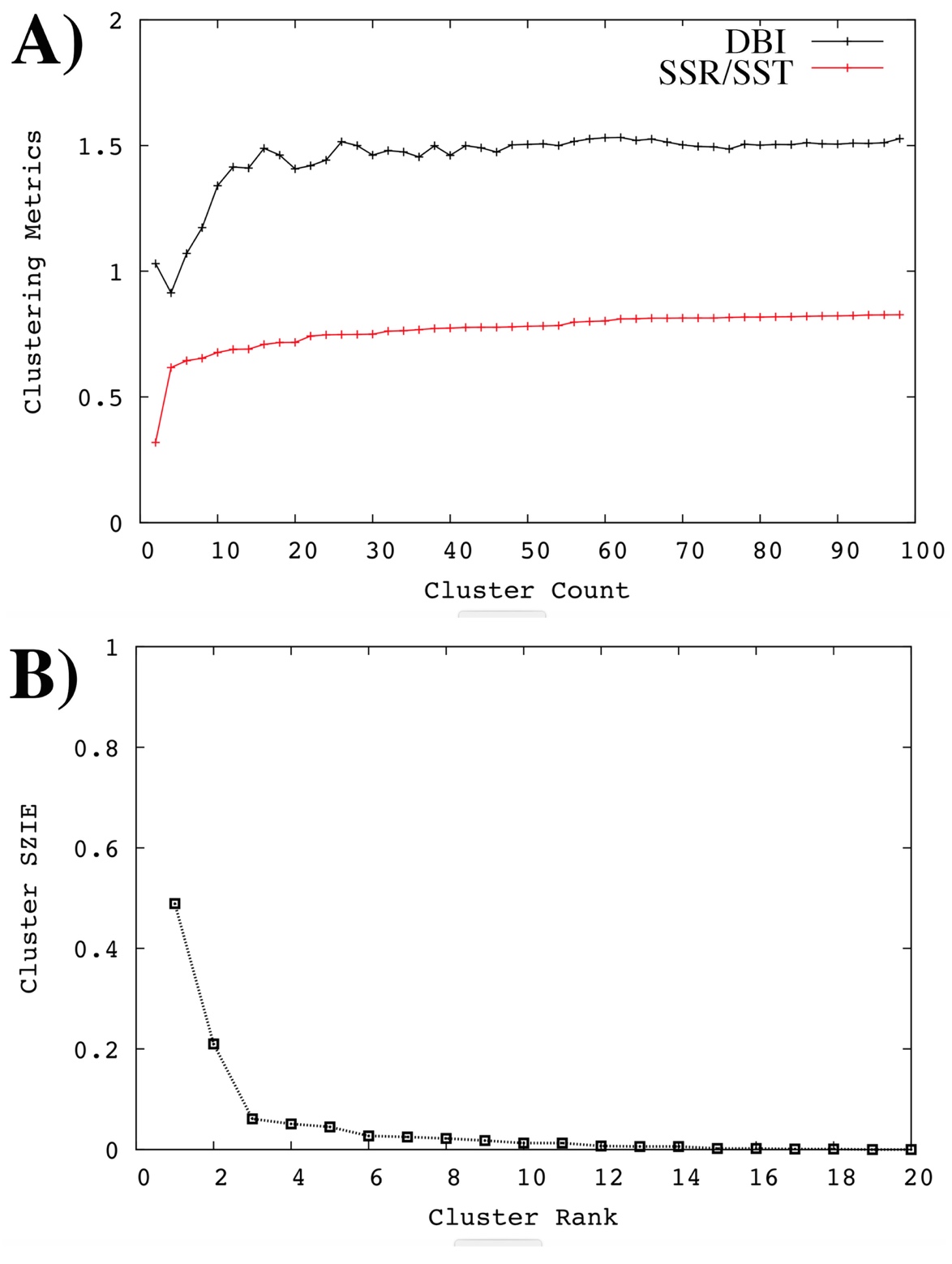

**Figure S15.** Persistence of the important hydrogen bonds for the BOUND systems.

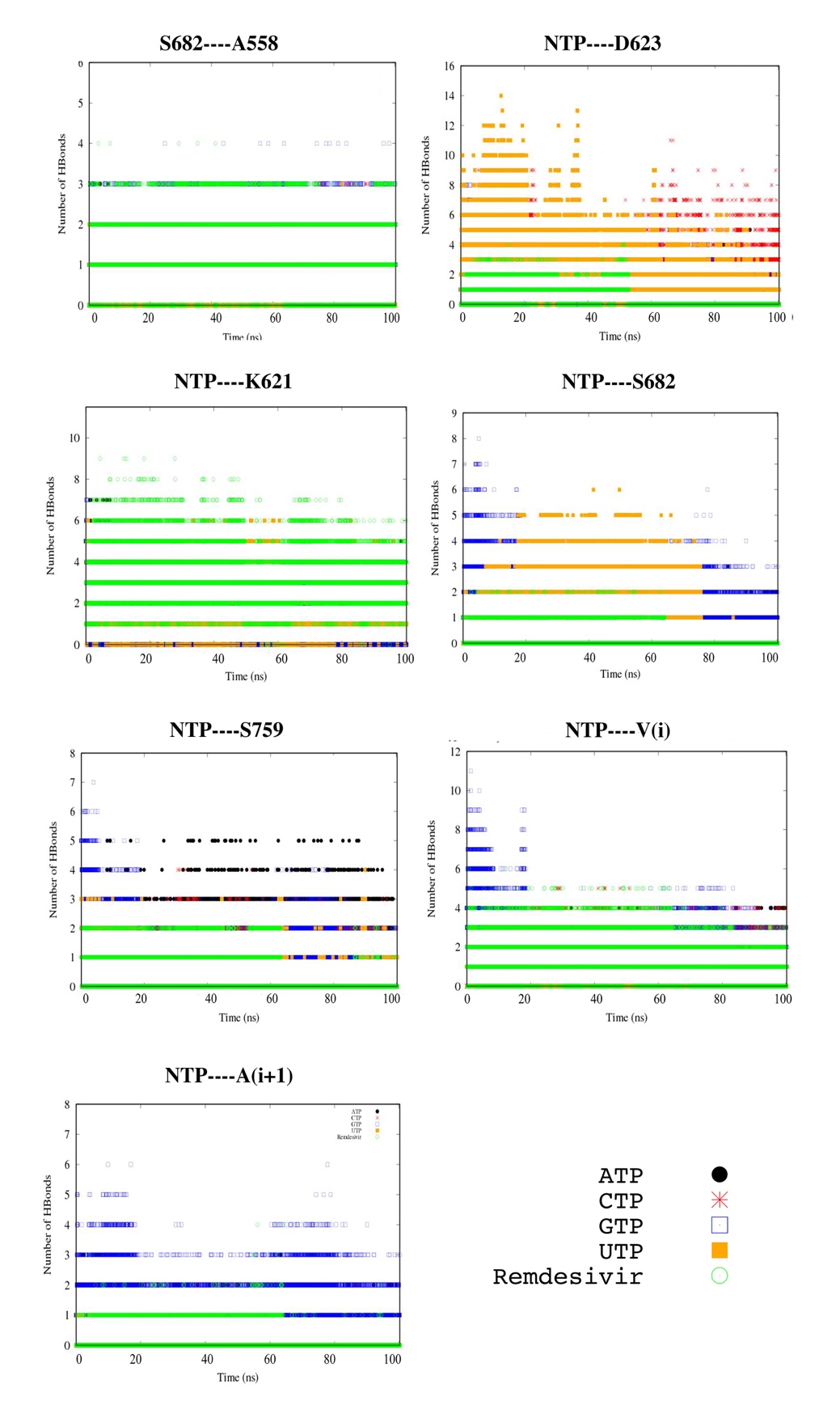

**Figure S16.** Persistence of the important hydrogen bonds for the FREE systems.

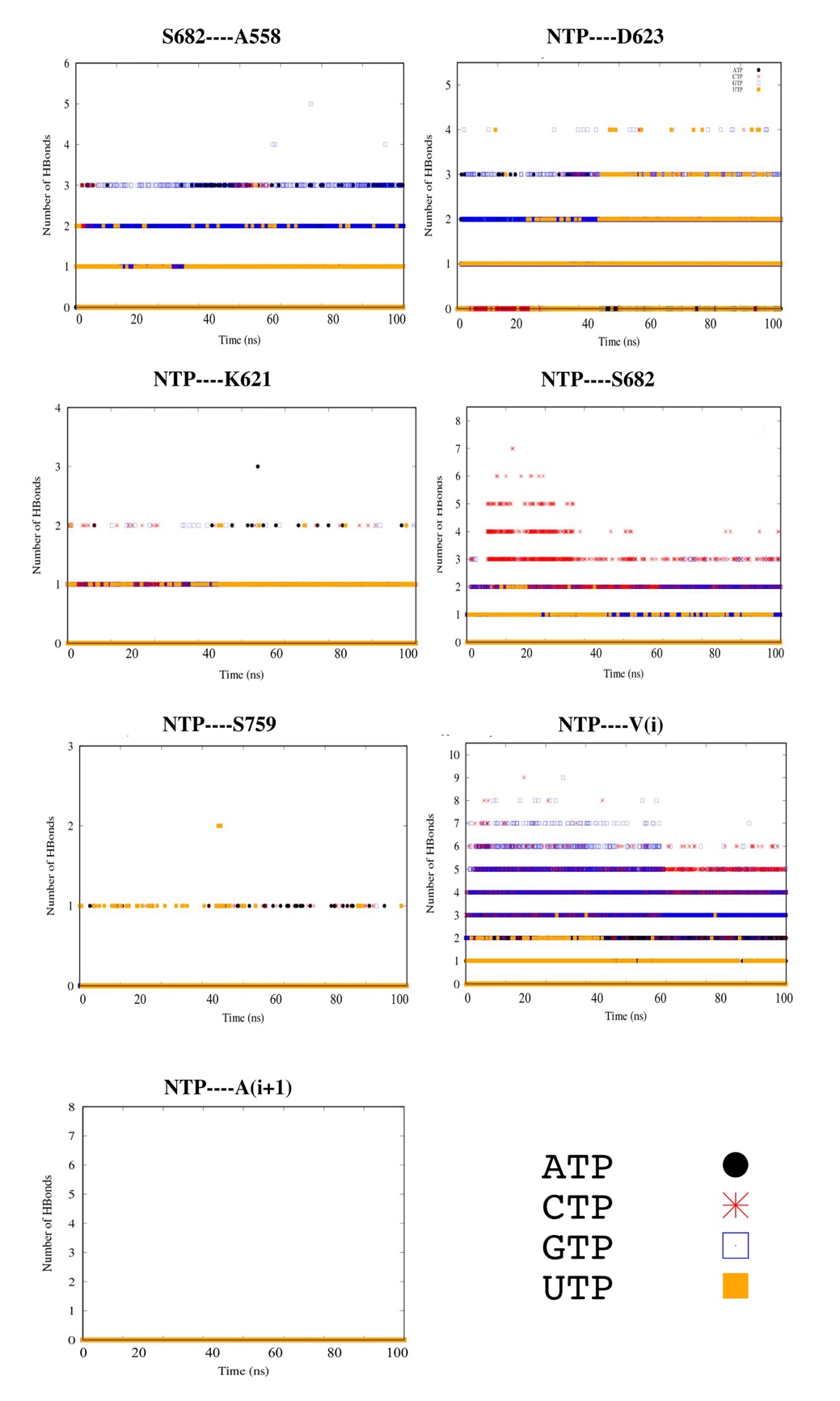

**Figure S17.** Persistence of hydrogen bonds between nsp12 protein and the RNA template at the template entrance gate for the BOUND systems.

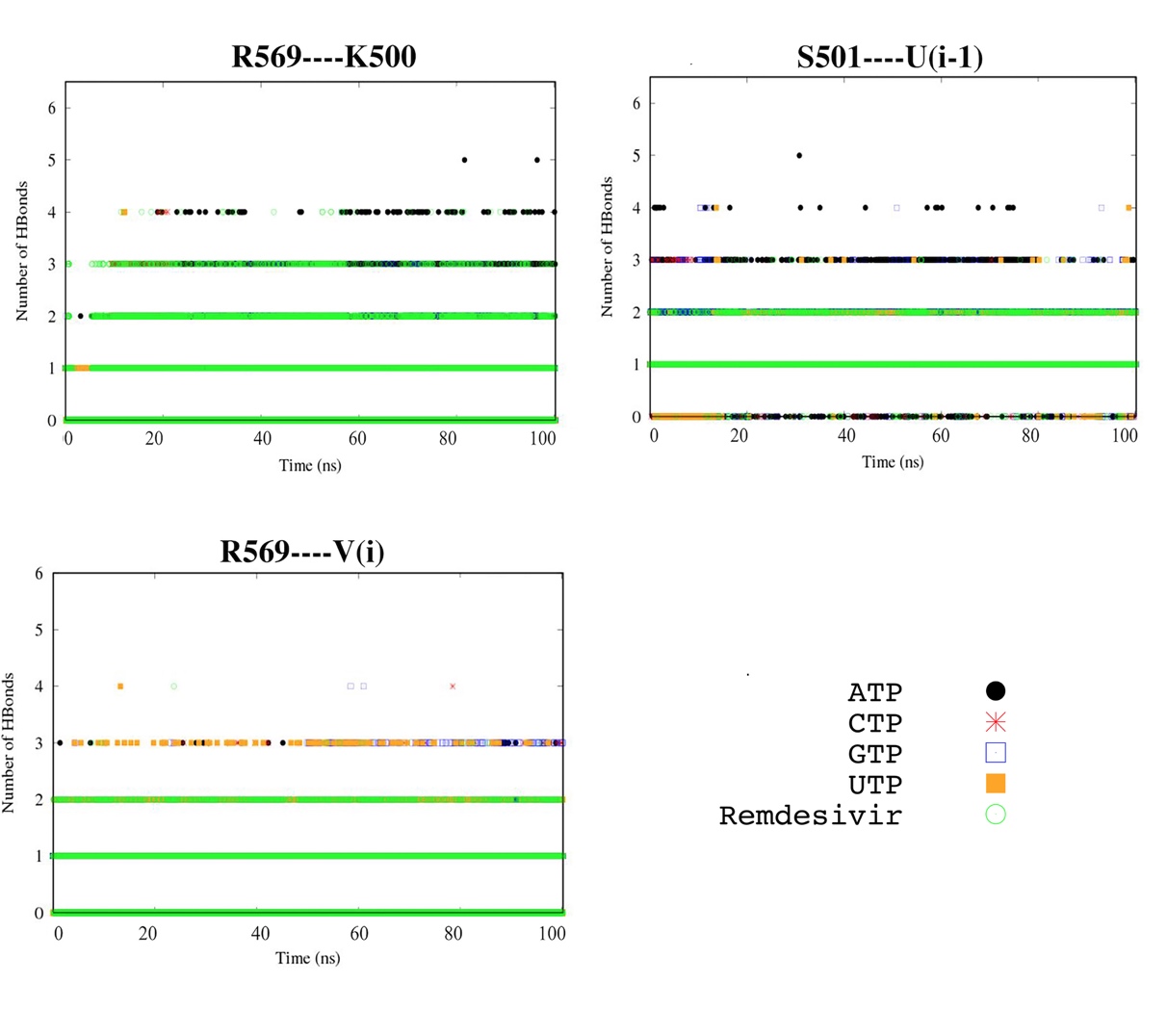

**Figure S18.** Persistence of hydrogen bonds between nsp12 protein and the RNA template at the template entrance gate for the FREE systems.

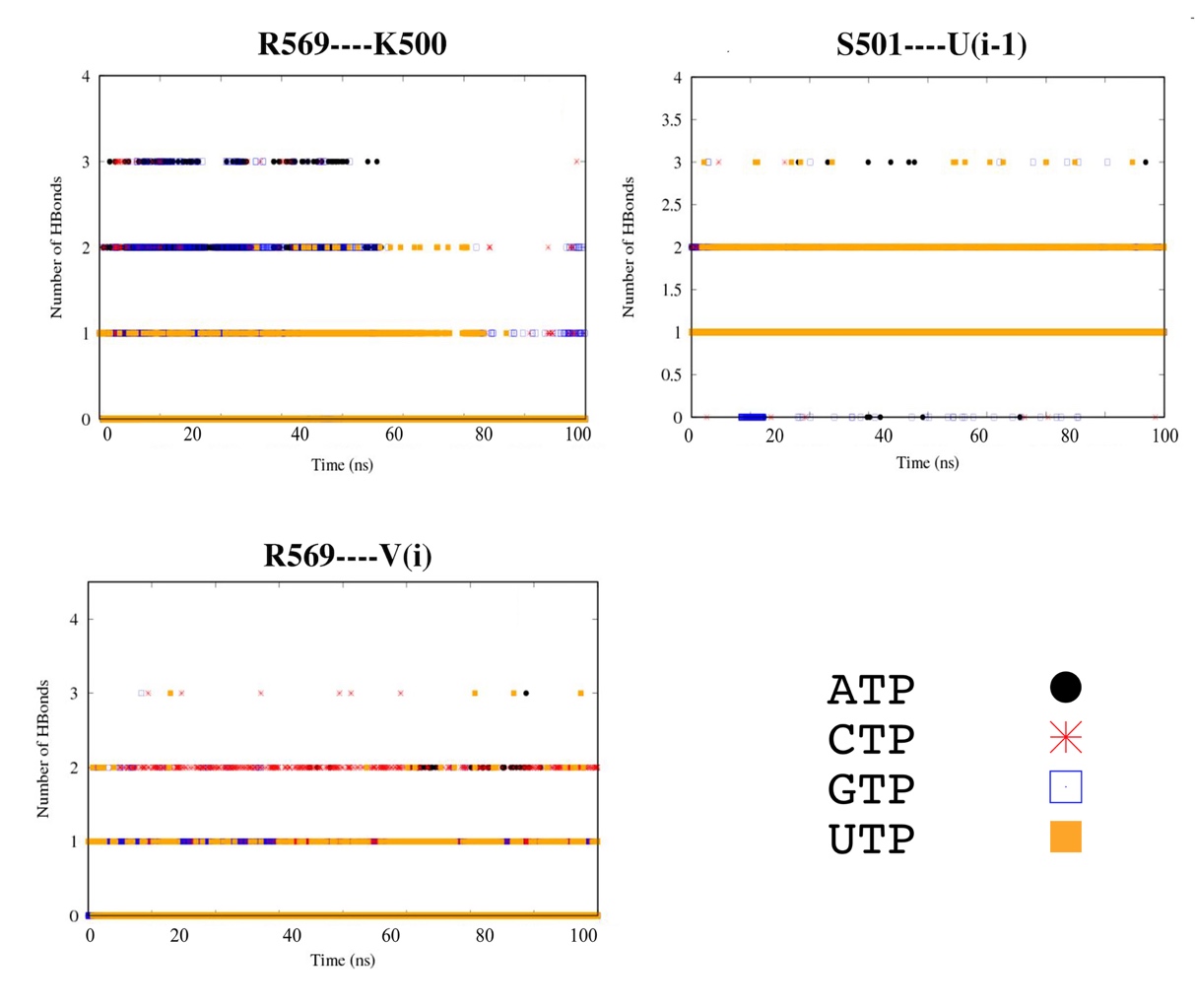

**Figure S19.** Dihedral angles for the key residues for the BOUND systems.

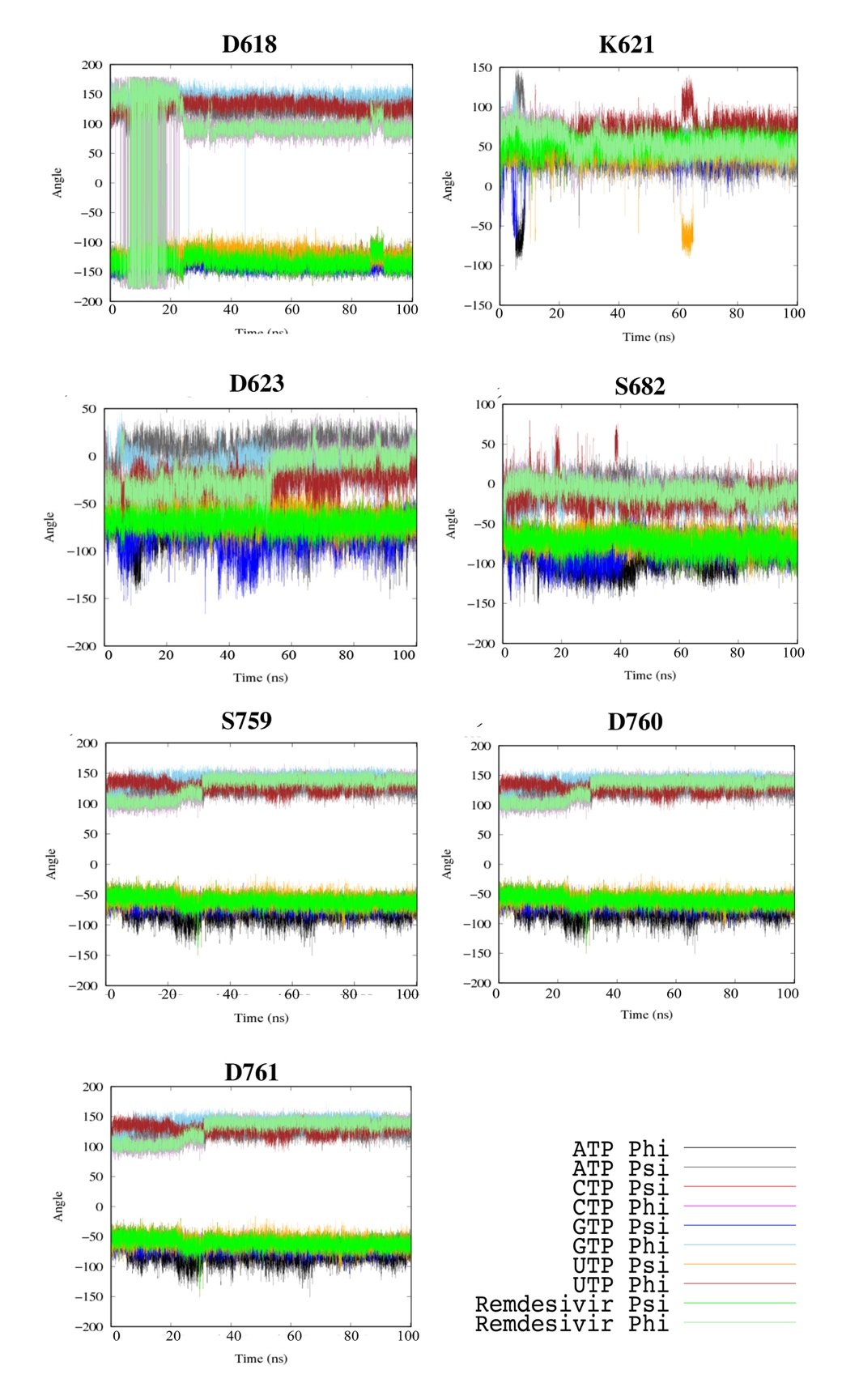

**Figure S20.** Dihedral angles for the key residues for the FREE systems.

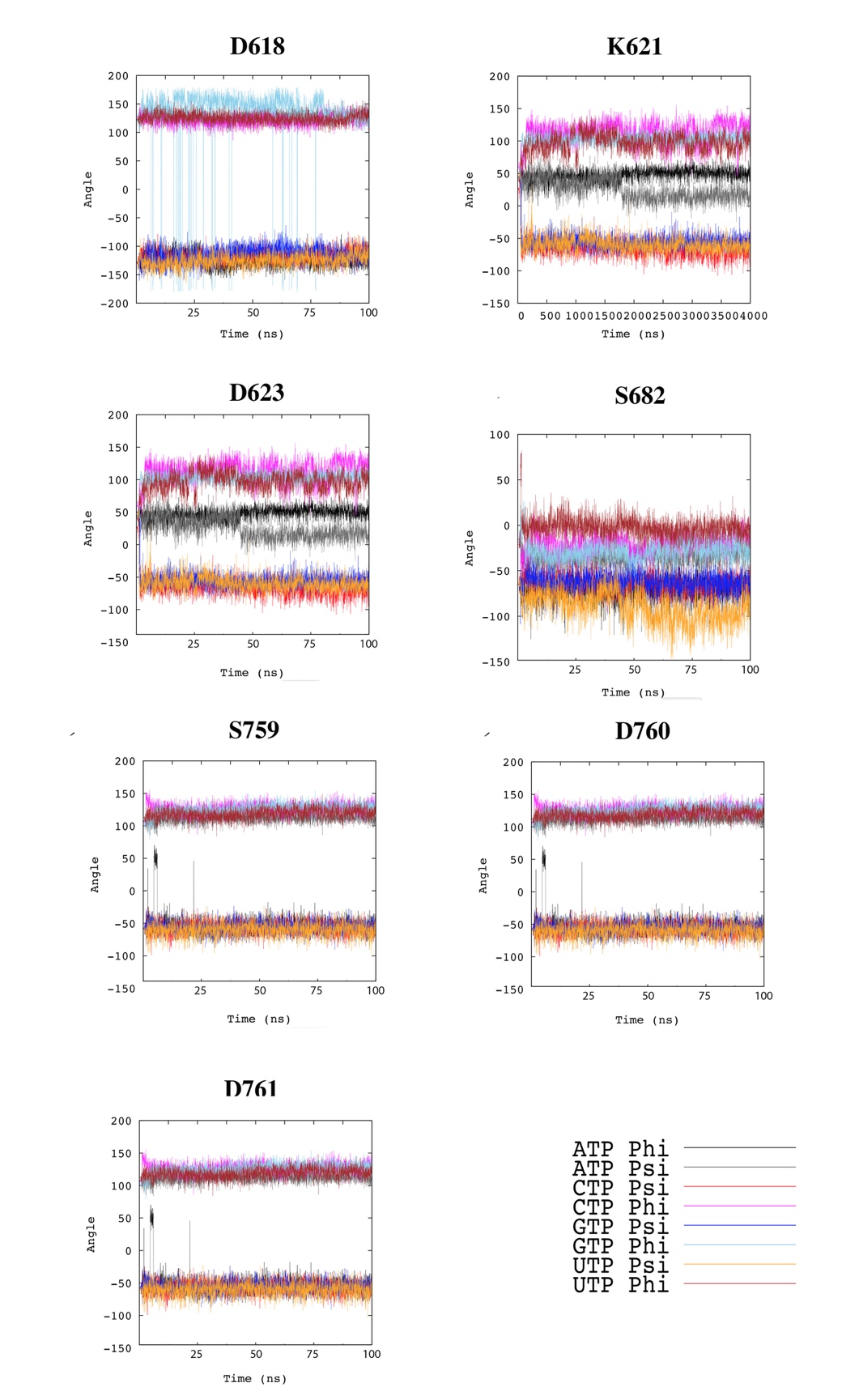

**Figure S21.** The electrostatic potential for the environment surrounding the two chelated ZINC ions.

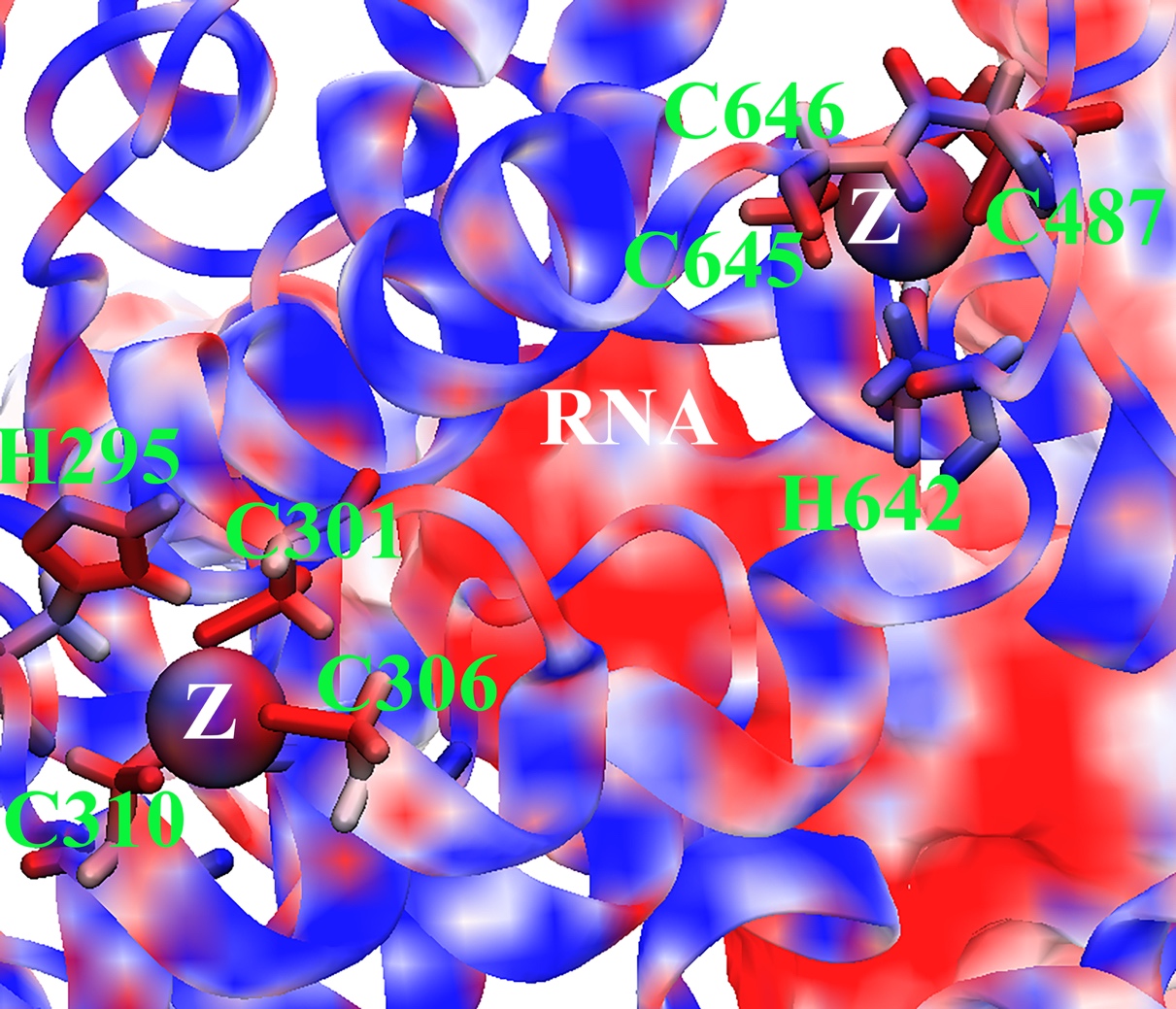

**Figure S22.** Non-bonded interactions with the two magnesium ions and three aspartate residues for the five BOUND nucleotides.

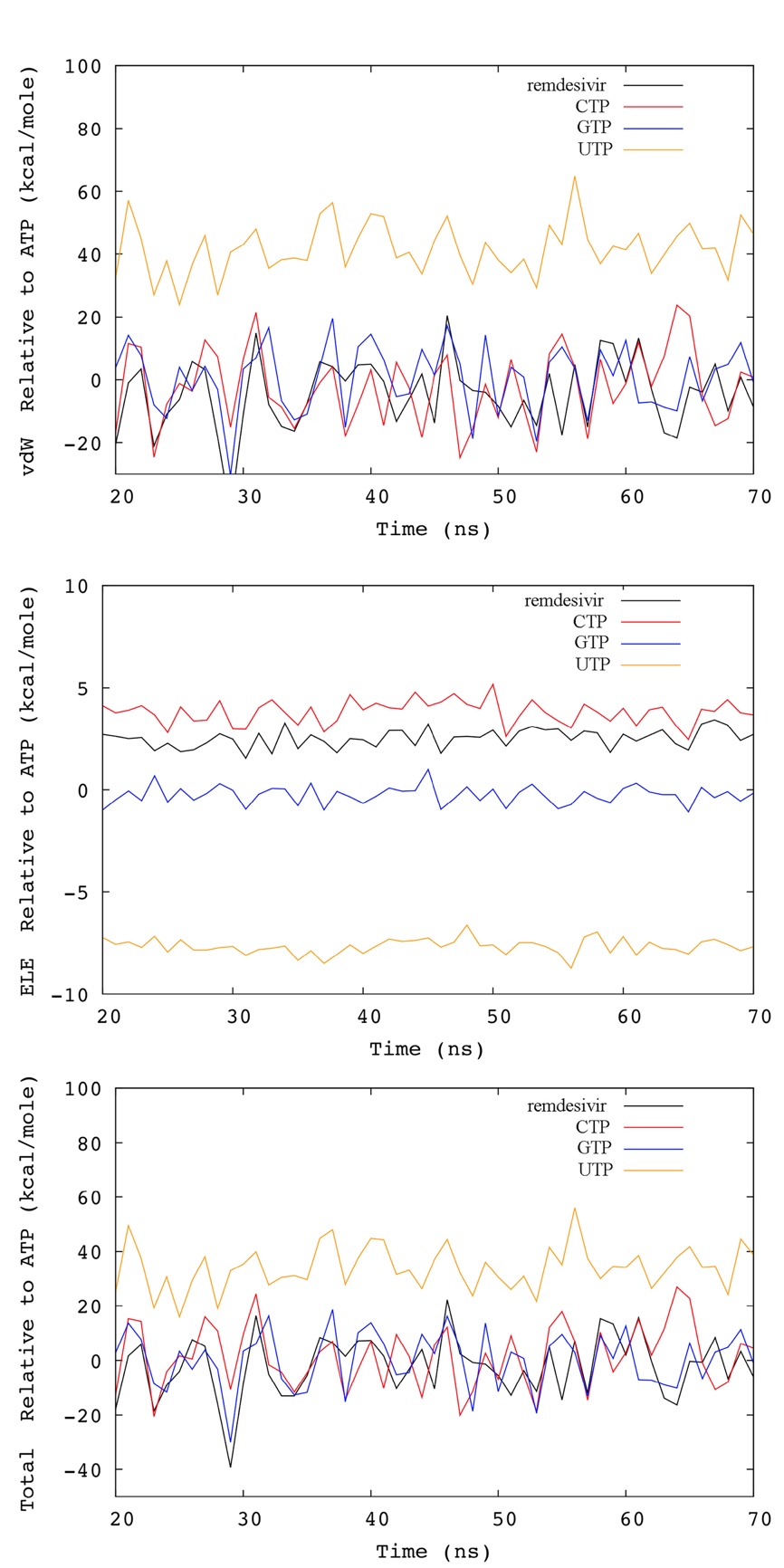

**Figure S23.** Non-bonded interactions with D623 for the five BOUND nucleotides.

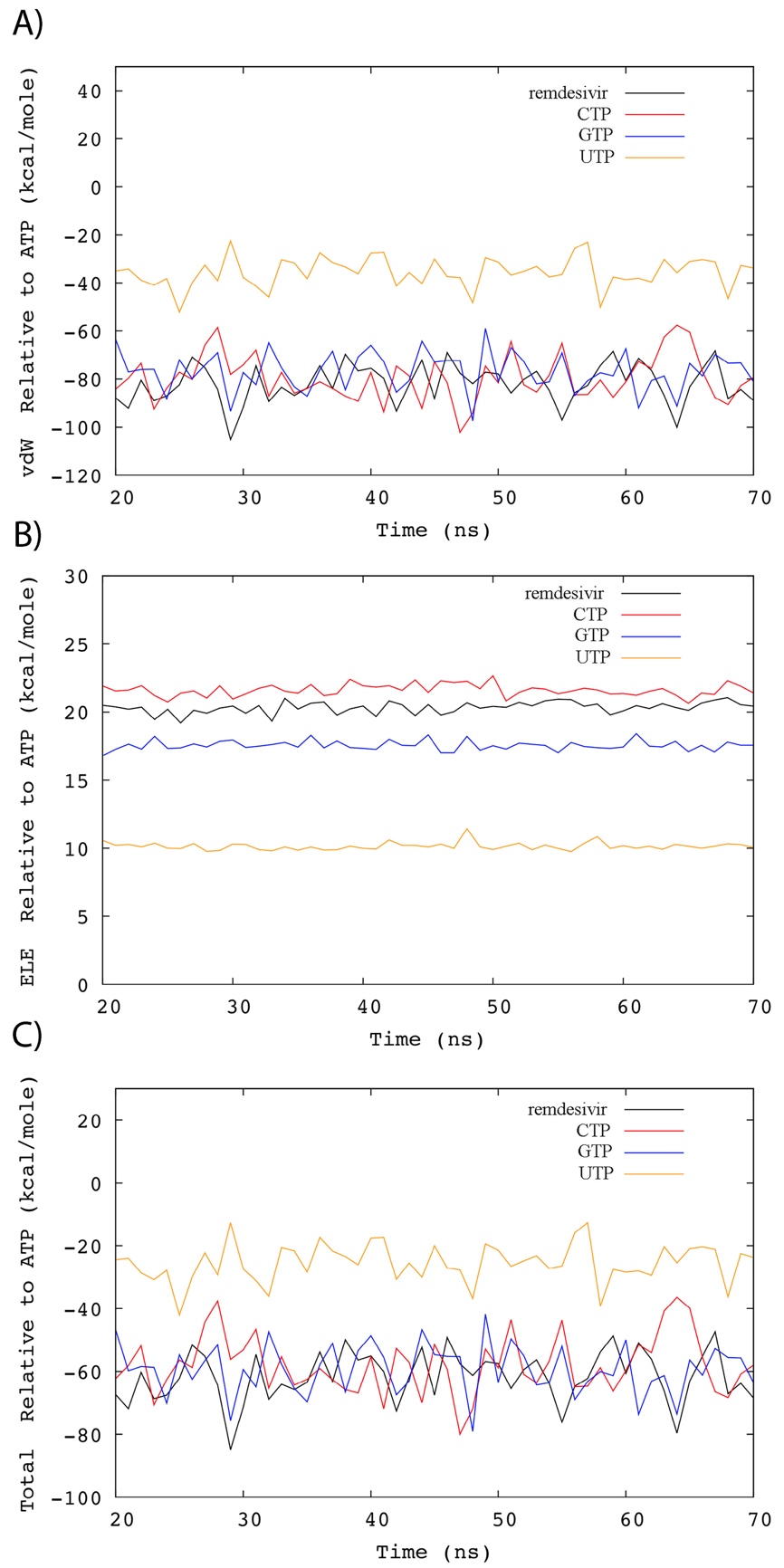

**Figure S24.** Non-bonded interactions with S795 for the five BOUND nucleotides.

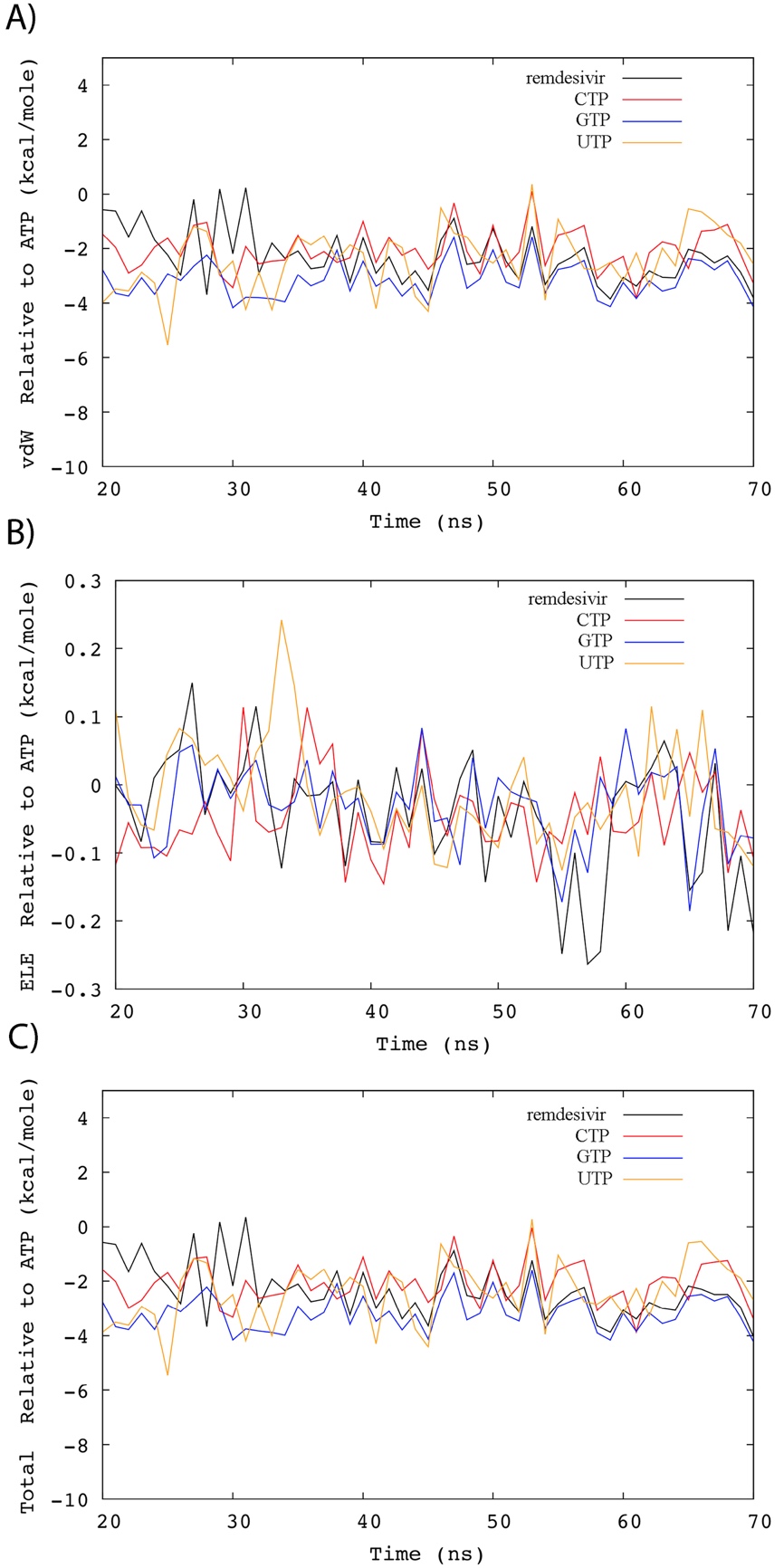

**Figure S25.** Non-bonded interactions for the whole RNA for the five BOUND nucleotides.

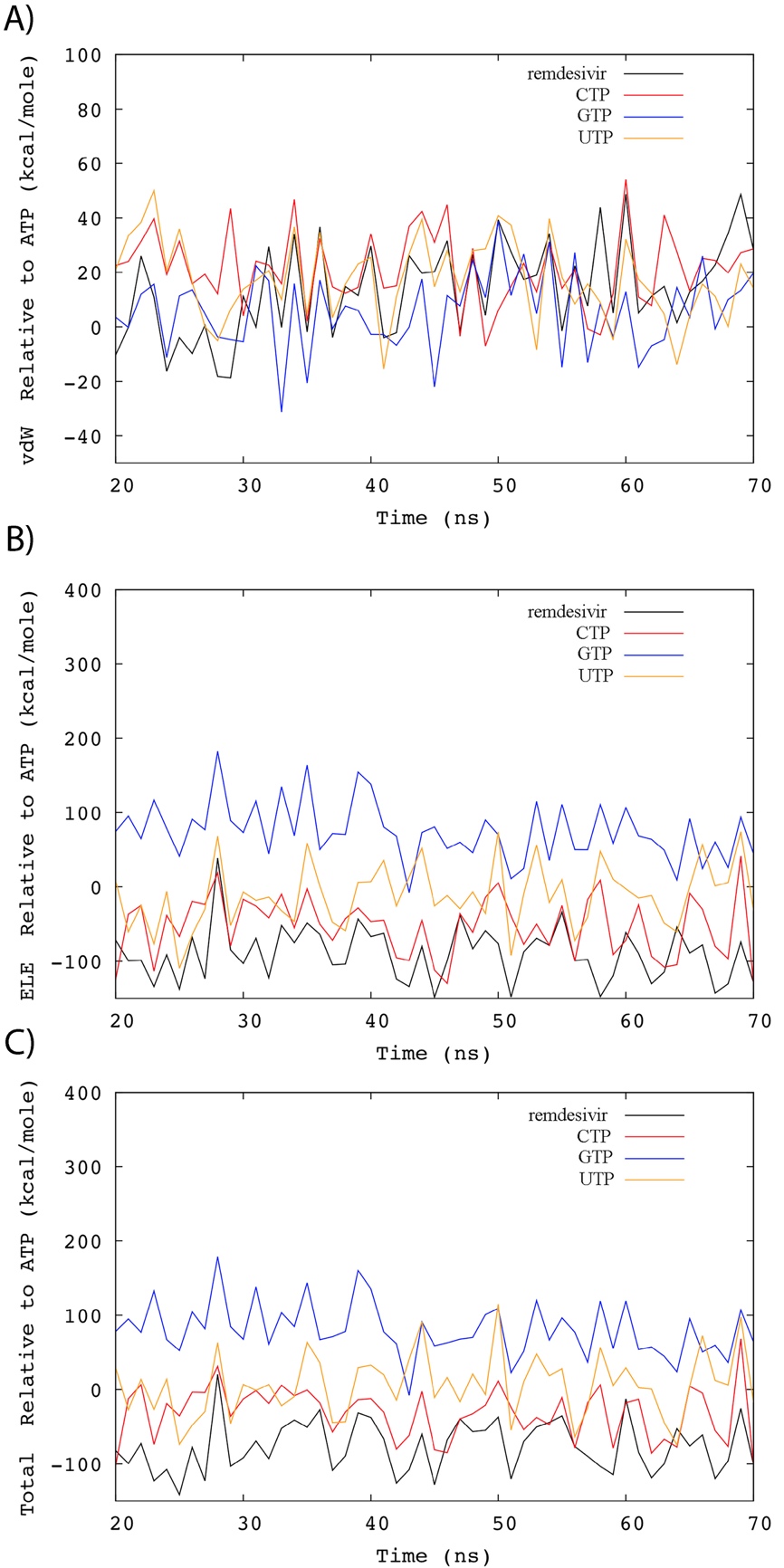

**Figure S26.** Water accessible surface area (SASA) around the different NTPs in the BOUND (**A**) and FREE (**B**) systems.

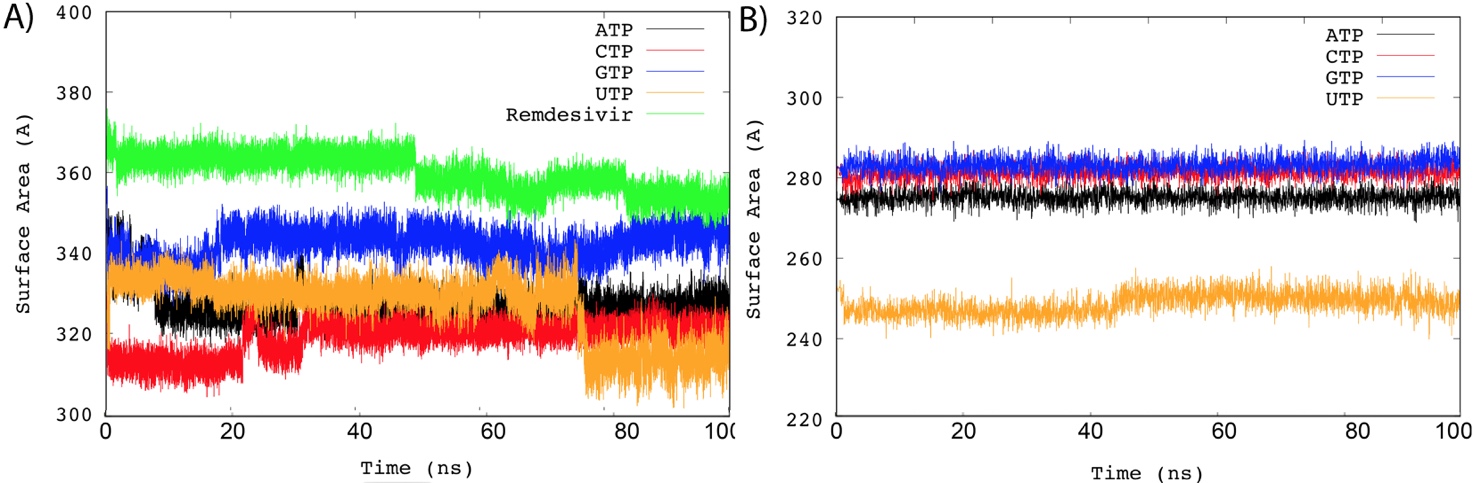

**Figure S27.** Close up on the three identified pockets. **A)** β-hairpin pocket, **B)** pocket 2 and **C)** pocket 3.

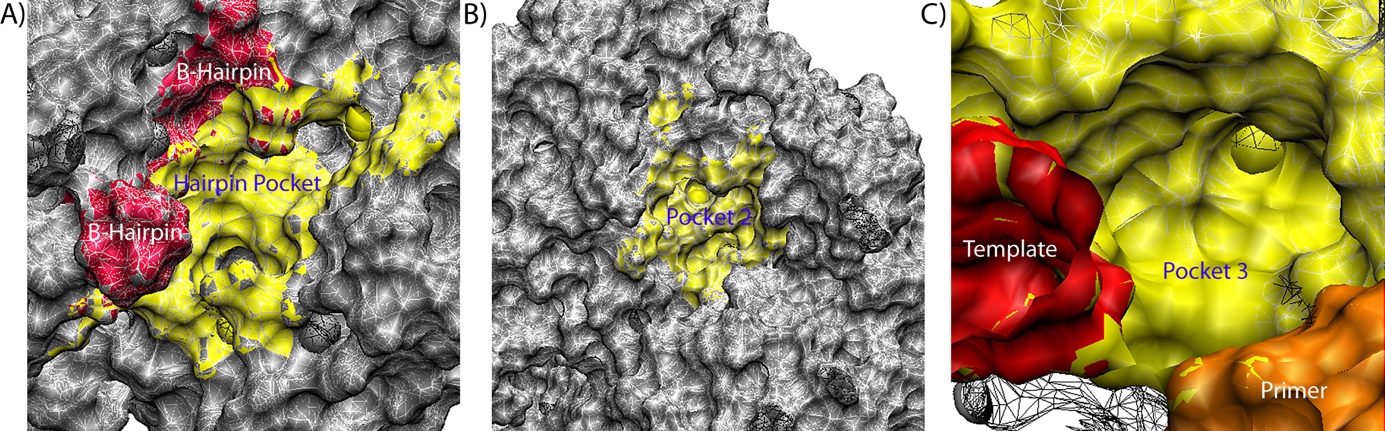

**Figure S28. Water shell analysis for A) catalytic site, B)** β-hairpin pocket, **C)** pocket 2 and **D)** pocket 3.

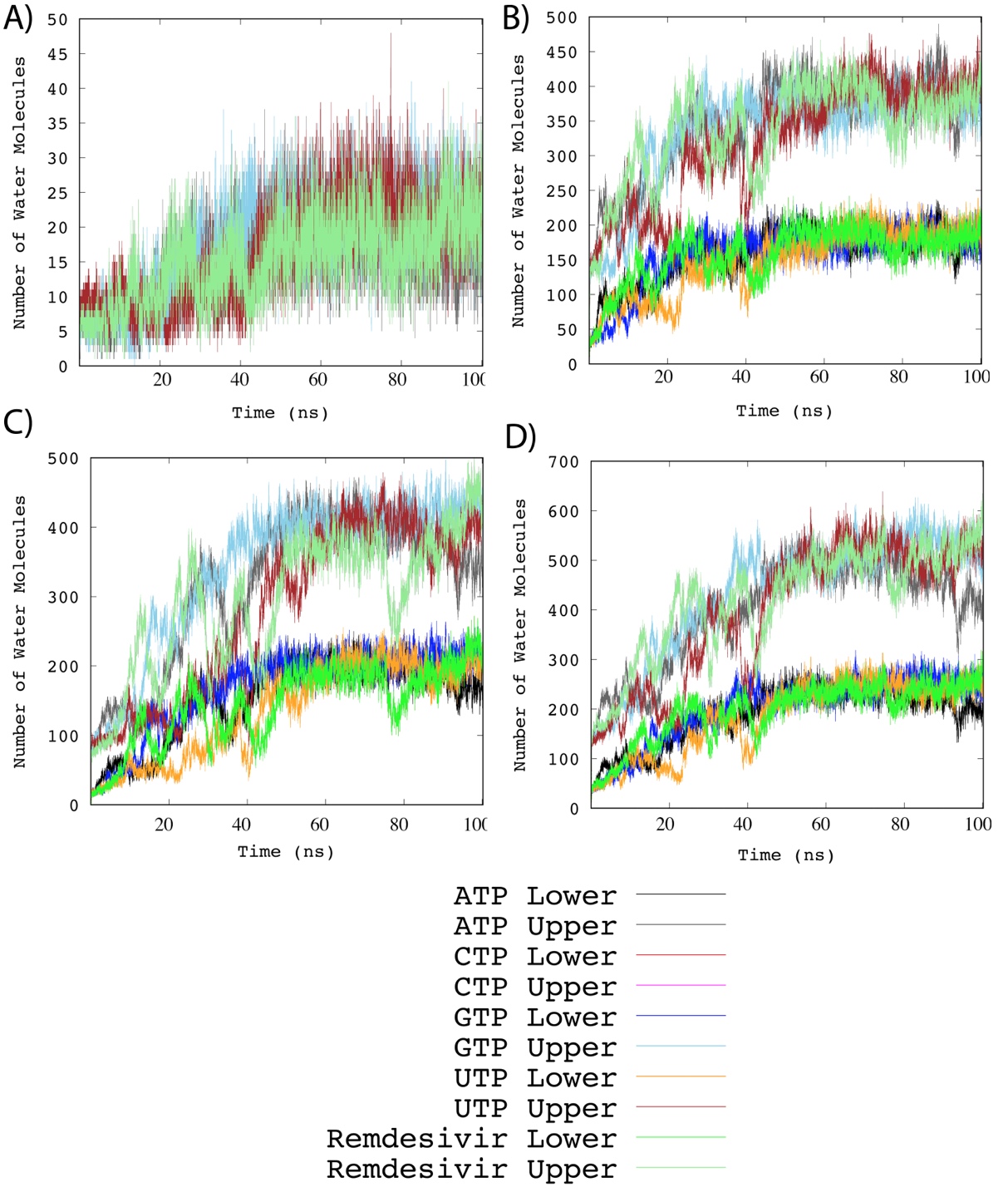

**Figure S29.** Protein heavy atoms RMSD values for the last 80 ns of the classical vs accelerated MD trajectories.

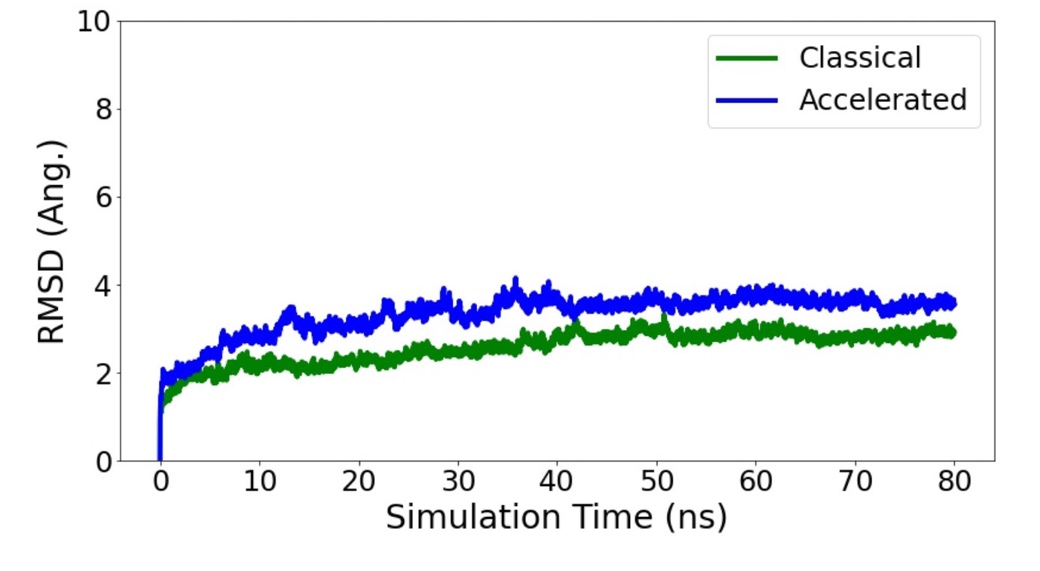

**Figure S30.** Electrostatic potential map for the β-hairpin pocket.

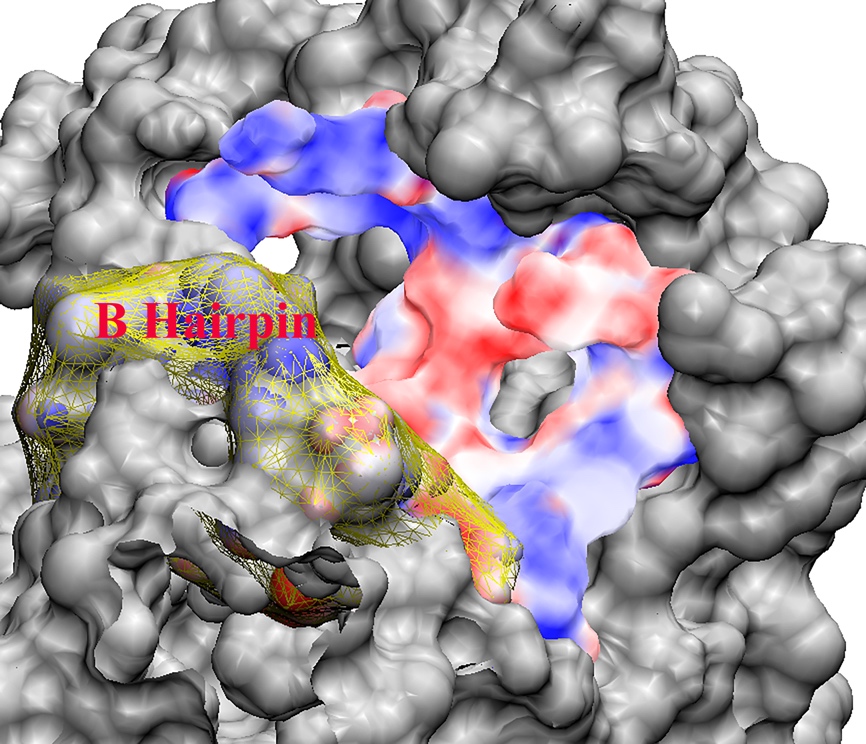

**Figure S31.** RMSD and Atomic fluctuations for NiRAN interface domain for the BOUND (A, B) and FREE (C, D).

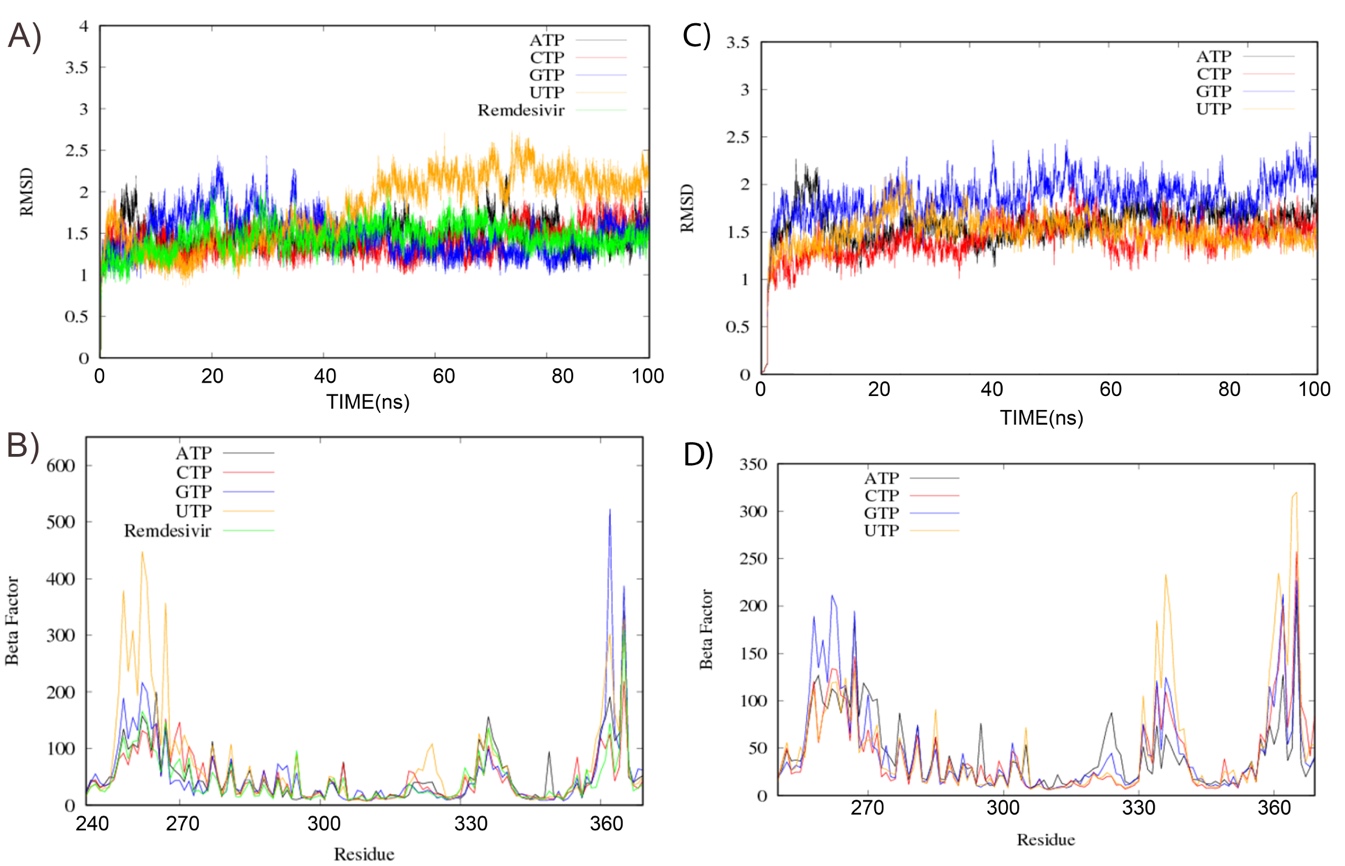

**Figure S32.** Overlay of the HCV NS5B polymerase (green) on top of the nsp12 polymerase (white). The four allosteric binding sites (thumb I, II and palm I, II) are also shown and their positions are compared to the three identified pockets from the current study. While the β-hairpin pocket and pocket 2 are unique to nsp12, pocket 3 in nsp12 overlaps with the palm pockets in NS5B.

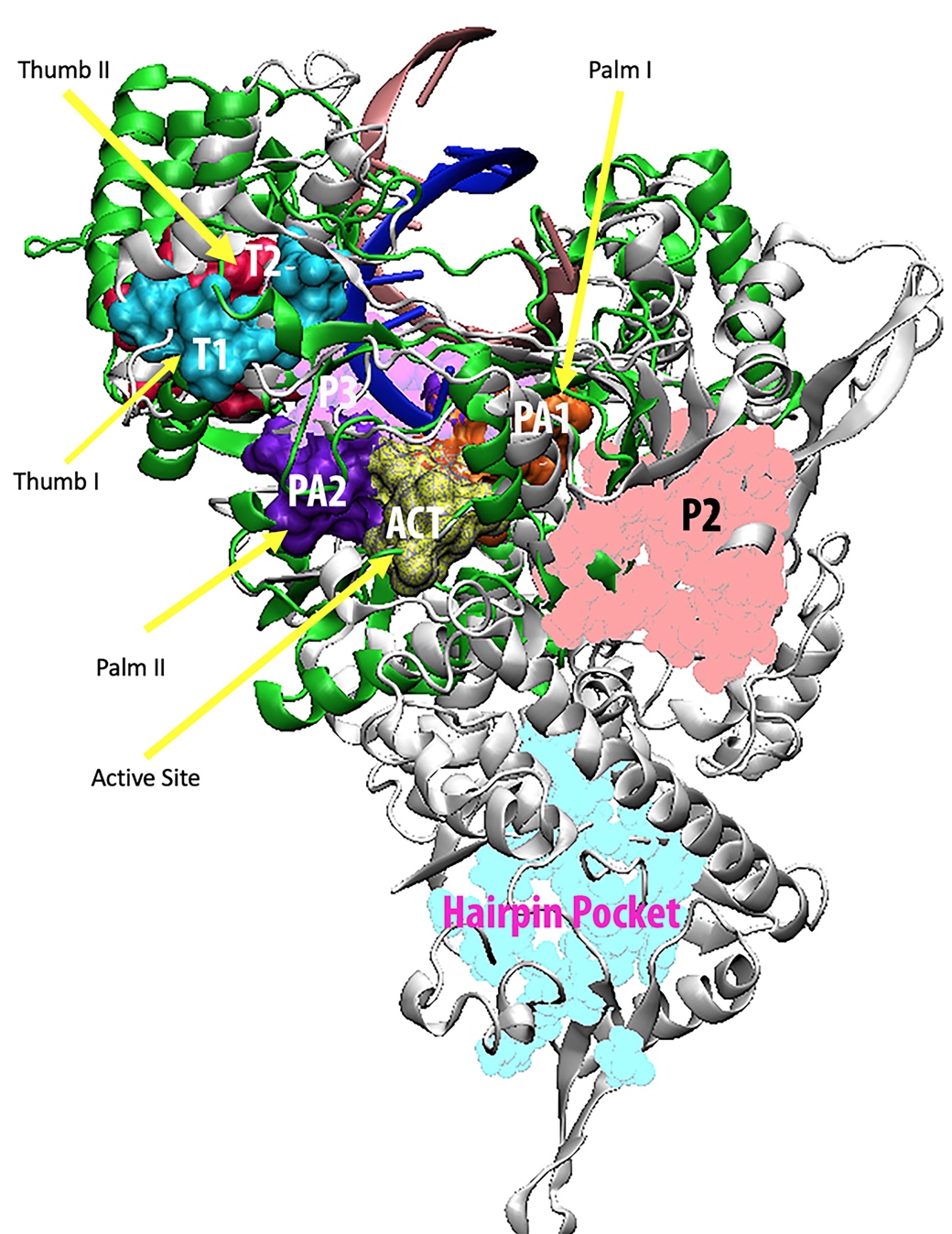

**Figure S33.** Amino acid sequences for the nsp12 (**A**), nsp8 (**B**), and nsp7 (**C**).

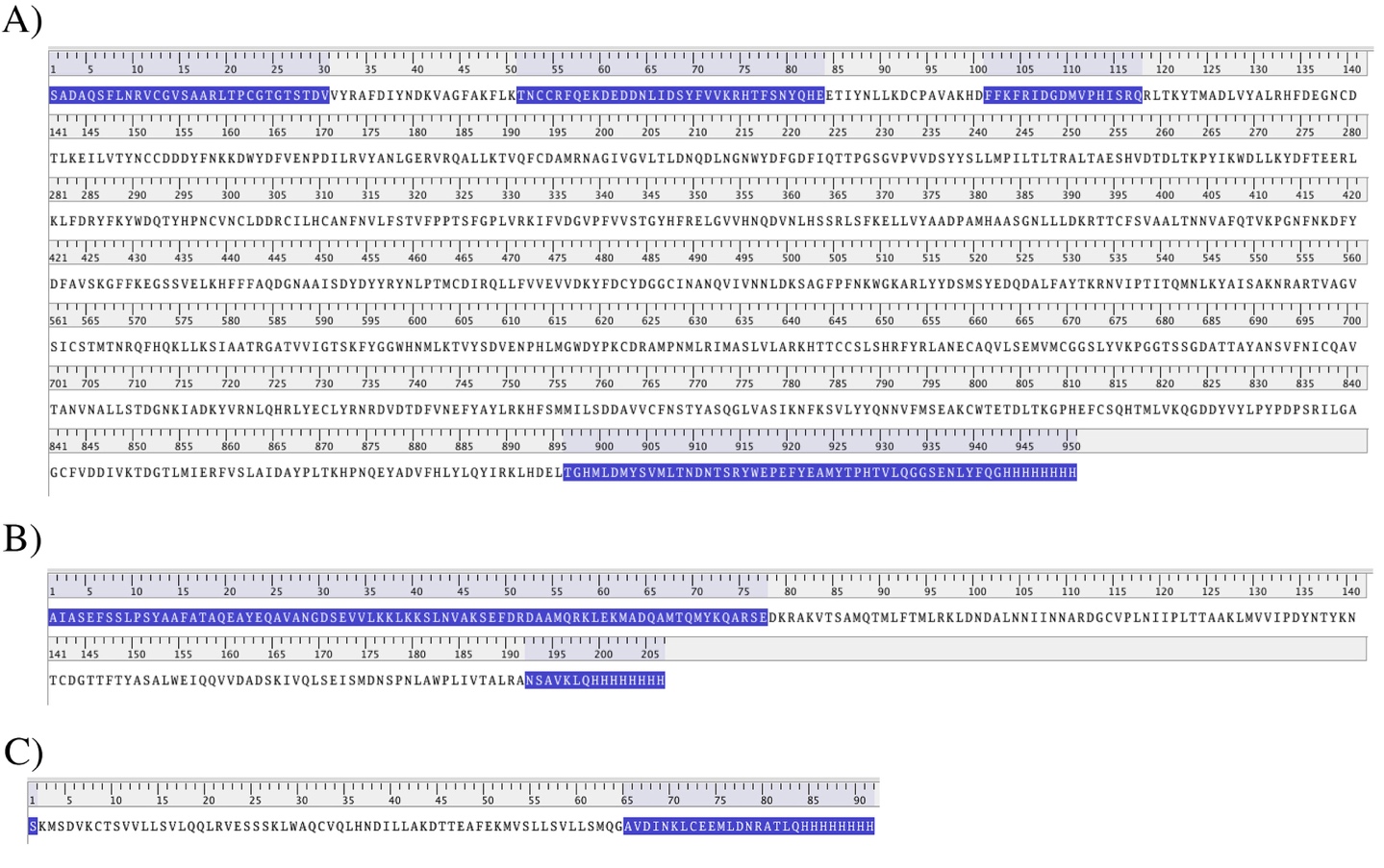
